# Gene regulatory network transitions reveal the central transcription factors in lung adenocarcinoma progression

**DOI:** 10.1101/2025.05.13.651789

**Authors:** Upasana Ray, Adarsh Singh, Debabrata Samanta, Riddhiman Dhar

## Abstract

Transcription factors play a central role in cancer growth, progression, and metastasis, and contribute to intratumor phenotypic plasticity which enable drug tolerance and cancer relapse. Changes in the regulatory activities of transcription factors in cancer may not always be detected by investigating mutational signatures or differential expression of the transcription factor, as done in traditional analysis. In addition, past studies have focused on the activities of transcription factors in tumors as a whole and thus, have not fully captured the heterogeneity in gene regulation among different cell types within the tumor microenvironment. In this work, through an analysis of the transitions in regulatory network architecture and gene regulation dynamics, we identify the central transcription factors associated with lung adenocarcinoma progression. The gene *NR2F1*, associated with neurodevelopment and cancer dormancy, emerge as a key transcription factor in the progression of lung adenocarcinoma. We further identify transcription factors that are only active in cancer samples and uncover how changes in gene regulation dynamics influence intratumor heterogeneity. Taken together, our work elucidates the transitions in gene regulatory network during cancer progression, identifies central transcription factors in this process, and reveals the complex regulatory changes cooccurring in different cell types within the tumor microenvironment.

## Introduction

Cancer undergoes a progressive transformation through different stages that manifest as the hallmarks of cancer, including sustained proliferation, resistance to programmed cell death, immune evasion, and metabolic remodeling^1^. Genetic mutations have a key role in cancer initiation and progression^2,3^. In addition, non-mutational processes also contribute greatly to this process^4-6^, and thus, epigenetic reprogramming and phenotypic plasticity are now considered to be hallmarks of cancer ^7^. The conversion of normal cells to cancer cells disrupts the homeostasis and presents new challenges for the cells in the tumor microenvironment^8,9^. Changes in gene regulatory network (GRN) architecture and activities of transcription factors (TFs), enable cancer cells to remodel gene expression programs and respond effectively to these challenges^10-12^. In this regard, the role of TFs is well studied in the epithelial-to-mesenchymal transition (EMT)^13,14^ - a key step in cancer progression that allows cancer cells to migrate and metastasize distant tissues. Thus, the gene regulatory network and the TFs have a central role in progression and phenotypic transformation of cancer.

Transcription factors also contribute to intratumor heterogeneity and phenotypic plasticity which have important implications for cancer progression and anticancer therapy resistance^15-17^. For example, antagonistic activities of TFs allow cancer cells to exist in a spectrum of intermediate hybrid E-M states^18,19^ that increase phenotypic plasticity^20,21^. In addition, several TFs are associated with generation of cancer stemness^22-24^ and thus, can drive phenotypic plasticity, affecting cancer progression and anticancer therapy^25,26^. Further, TFs have been linked to generation of stochastic variation in gene expression^27,28^, which further contributes to phenotypic heterogeneity with implications for cancer progression and therapy resistance^29,30^.

Given the importance of TFs in cancer transformation and phenotypic plasticity, they could be good targets for anticancer therapy^31-33^ and recent research has uncovered many potential ways to target them^34,35^. Thus, there has been an increasing interest in identifying the key regulators of cancer progression and plasticity across different cancer types^36-39^. Traditionally, cancer driver genes have been identified from the studies on mutational signatures^40^, and mutations in the non-coding regions have been used to identify the TFs with altered regulation in cancer^41,42^. Alternatively, key TFs have been identified from changes in gene expression pattern between cancer and normal samples^43^. However, the expression pattern of the TFs driving changes in gene regulation may not always change^44^, as epigenomic changes can alter activities of TFs on their target genes^6^. Therefore, investigating changes in the topology of the GRN and the regulation dynamics of targets of specific TFs becomes essential for identifying the key regulators of cancer ^45-47^.

It is important to note that the tumor microenvironment (TME) comprises of many different cell types, each of which contributes to tumor progression^48,49^. Thus, changes in gene regulation in one cell-type in the microenvironment may elicit a response from the other cell types, through a corresponding change in their regulatory network. Indeed, the stromal and the fibroblast cells present in the tumor microenvironment can drive cancer^50,51^, and targeting them could improve cancer treatment^52,53^. Earlier studies on cancer gene regulatory network have focused on specific cancer cell lines or on whole tumor samples^45-47,54^. Hence, they have not captured the simultaneous changes occurring in different cell types present in the TME, and thereby, have not revealed the full complexity of the regulatory changes occurring in the TME.

In the present work, we identified the key transcription factors by investigating the transitions in the GRN architecture and regulation dynamics occurring with lung adenocarcinoma (LUAD) progression. The transcription factor NR2F1, that has been associated with cancer dormancy, emerged as one of the central TFs in LUAD progression. We also uncovered the coordinated changes in gene regulation occurring across different cell types within the TME and identified specific TFs that are uniquely active in cancer samples and in specific cell types, but not in normal samples. We further investigated the transitions in gene regulation dynamics with cancer progression using a two-state model of gene expression. Here again, NR2F1 emerged as one of the central TFs, thereby, highlighting its key role in LUAD progression. We further linked the changes in regulation dynamics to intratumor expression heterogeneity which can also drive phenotypic plasticity. Taken together, our work uncovered the transitions in the GRN architecture and dynamics through cancer progression, identified the key TFs in this process, and revealed the complex regulatory changes occurring in the TME.

## Results

### Transitions in the gene regulatory network (GRN) architecture through cancer stages reveals the central TFs

To investigate the transitions in the GRN architecture during cancer progression, we reconstructed the active GRNs using single-cell RNA sequencing (scRNA-seq) data obtained from the LUAD patients at different stages of cancer. We obtained single-cell RNA-seq data from 26 samples comprising of 11 adjacent normal samples, 8 stage I samples, one stage II sample, two stage III samples and four stage IV samples. We did not consider samples from metastasis in other tissues, as the differences in tissue environment could affect TF activity and thus, could confound our analysis. We investigated GRN transitions in three cell-types – Epithelial, Myeloid and T/NK cells, which had >100 cells in all samples required for reconstruction of the GRN.

For reconstructing active GRNs from scRNA-seq data, we followed an approach described in the method bigScale2 ^55^, but with major modifications. For every cell-type in a sample, we calculated centered expression correlation between all pairs of genes using bigScale2 (Fig. 1A). However, we investigated the active GRN using frameworks that were different from the approach described in bigScale2 and could better capture the transitions in the GRN with cancer progression. For epithelial cell-type, we could not calculate correlation values for one stage I sample (see Methods) and we had to discard the stage II sample from our analysis due to potential bias in calculation of expression correlation, likely due to a low cell number in this sample. Hence, we performed all analyses on the remaining 24 samples. For myeloid and T-NK cell-types, we included all 26 samples in our analysis.

**Figure 1.**
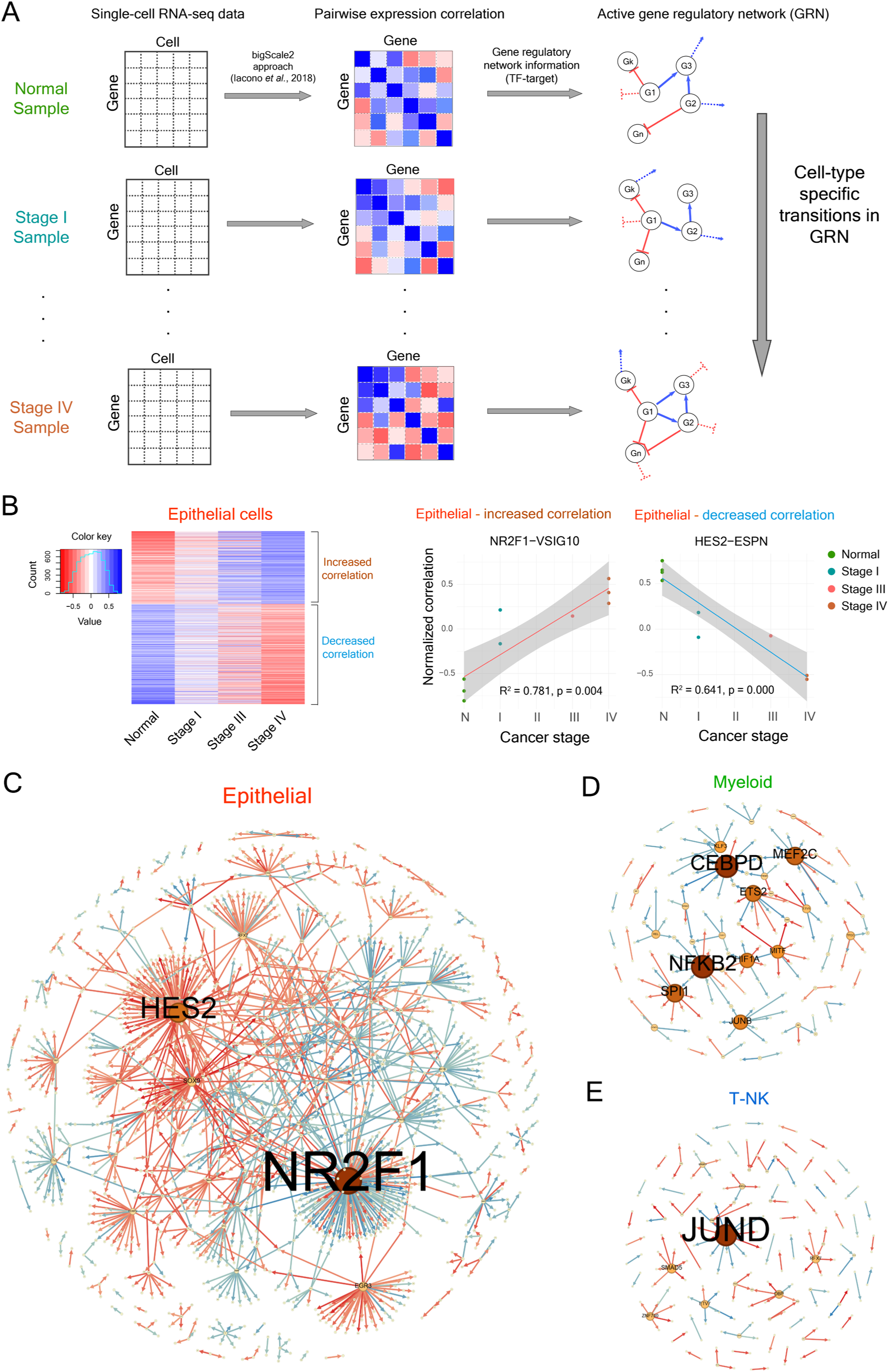
Reconstruction of GRNs from scRNA-seq data in LUAD samples from different stages reveal transitions in the GRN architecture with cancer progression and identify the central TFs. **(A)** For reconstruction of active GRNs in patient samples, pairwise correlation values between genes were calculated using the bigScale2 method (Iacono et al., 2018) for each cell-type across samples from different cancer stages. The correlation values were considered only for experimentally verified TF-target connections for which potential binding motif of the TFs in the target promoters were identified, to ensure inclusion of only direct TF-target regulations in our analysis. **(B)** Heatmap showing the changes in mean correlation values with cancer progression for the TF-target connections that showed linearly increasing (red to blue) or decreasing (blue to red) correlation trends in epithelial cells. These TF-target connections were identified by linear regression where goodness-of-fit (R^2^) was greater than or equal to 0.5 and the slope value was beyond mean ± 2×s.d. values, where mean and s.d. values were calculated from the slope values of the liner regression of all TF-target connections in epithelial cells. **(C-E)** Graphs identifying the hub TFs that were associated with the greatest number of TF-target connections undergoing regulatory transitions in epithelial cells (C), in myeloid cells (D), and in T-NK cells (E). The size of a node and a gene label is proportional to the number of connections associated with the gene which underwent transitions with cancer progression. A red edge represents a negative correlation trend and a blue edge represents a positive correlation trend with cancer progression.

As correlation between two genes may not always indicate direct regulation by TFs, we worked with correlations of only a set of high-confidence TF-target connections obtained after a two-stage filtering process. First, we obtained the list of experimentally validated TF-target relationships from the published literature and databases (see Methods). We then used the binding motif information for the TFs to search for putative TF binding sites in the promoter regions of the target genes, after allowing for up to two mutations in the binding site compared to the motif. We considered only those experimentally verified TF-target relationships where we could find a binding motif of the TF in the promoter of the target (Fig. 1A).

In the next step, we normalized the distributions of correlation values of all samples of a cell-type using quantile normalization (Fig. S1-S3), to account for any technical variation arising in the sample preparation from different patients and in calculation of correlation values from different numbers of cells. To determine the transitions in the GRN of a specific cell-type with cancer stages, for every TF-target connection, we tested for changes in the normalized correlation values with cancer stages using linear regression (Fig. 1B). We calculated the goodness-of-fit (R^2^) and the slope values for all TF-target connections for which we had correlation values in at least two normal samples, at least two stage I samples, at least one stage II or III sample, and at least two stage IV samples. We considered the connections with R^2^ value greater than or equal to 0.5 (Fig. S4) and the slope value beyond 2 standard deviations (s.d.) from the mean slope value (Fig. S5) as the ones undergoing regulatory transitions with cancer progression. Epithelial cells showed extensive transitions in the regulatory network (in 1361 connections) (Fig. 1B), compared to myeloid (in 164 connections) and T-NK cells (in 133 connections) (Fig. S6, S7).

Next, we identified the hub TFs that were overrepresented among the TF-target connections that underwent transitions with cancer progression (Fig. 1C-E). In epithelial cells, the TFs NR2F1 and HES2 showed the greatest number of changes in correlation with their targets, followed by SOX9, EGR3, RFX7, FOXP2 and ETV2 (Table S1). Interestingly, the TF TEAD2, which regulates the expression of NR2F1, showed increased correlation in expression with NR2F1 with cancer progression (Fig. S8A). TEAD2 has been shown to promote cancer progression in hepatocellular carcinoma^56^. Expression of TEAD2 also increased with cancer progression in our dataset (Fig. S8B).

The transcription factor NR2F1 (also known as COUP-TFI) is a key regulator of neuronal development in mammals^57^. In this process, NR2F1 is known to control cell proliferation, adhesion, and migration^58-62^. In addition, NR2F1 has been reported to regulate cancer cell dormancy^63,64^, in which the SOX9 gene is also involved^63^. NR2F1 also prevents dissemination of early-stage breast cancer cells^65^. However, in salivary adenoid cystic carcinoma cell lines and in a xenograft model, overexpression of NR2F1 inhibited proliferation, but substantially increased migration and invasion abilities of cancer cells^66^. Consistent with these observations, silencing of the NR2F1 expression increased proliferation of cancer cells but reduced their invasive abilities^66^. The TFs HES2 and the members of HES family are predicted to be involved in several developmental processes^67^ and HES2 has been suggested to be a tumor suppressors gene in neuroblastoma^68^.

In myeloid cells, the TFs CEBPD, NFKB2, MEF2C, SPI1 and ETS2 showed the greatest number of changes (Table S2), and in T-NK cells, the TFs JUND and SMAD5 showed the greatest number of changes (Table S3). However, the number of changes in the myeloid and T-NK cells were much lower than the number of changes observed in the epithelial cells.

### Gain, loss and switching of regulatory interactions occur during cancer progression

To further understand the role of the hub TFs in cancer progression, we inferred the specific types of regulatory transitions occurring in their target genes. For example, a decreasing correlation trend could arise either from a gain of repression or a loss of activation (Fig. 2A). Similarly, an increasing correlation trend could arise due to a gain of activation or a loss of repression. To identify the types of regulatory transitions, we classified gene regulations into broadly three classes - gain, loss and switching types (Fig. 2A). We inferred these regulation transitions from correlation data using initial correlation strength and the slope of the linear regression line from the correlation trend (see Methods). We classified positive correlation as activation and negative correlation as repression and further classified them into weak and strong categories (see Methods, Fig. 2A). Regulations could also be of switching types, namely, from activation (weak or strong) to repression (weak or strong) and the vice-versa, which we identified from the changes in the sign of the correlation values (Fig. 2A).

**Figure 2.**
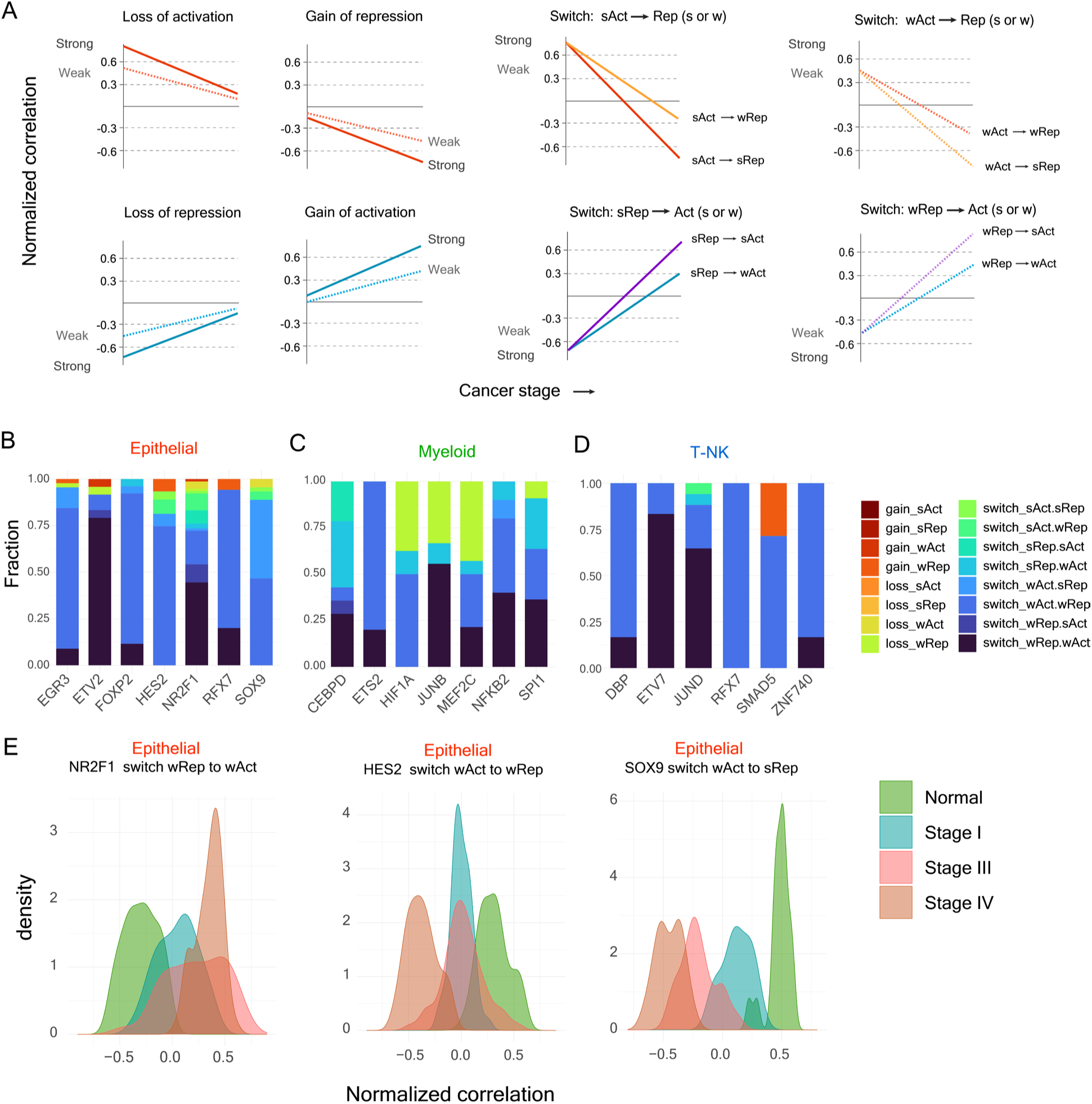
Regulatory transitions predominantly lead to switching between activation and repression. **(A)** Different types of changes in regulation can give rise to negative (top) or positive (bottom) correlation trends across cancer stages. The changes in regulation can include gain of activation or repression, loss of activation or repression, or switching between activation and repression. **(B-D)** Stacked bar plot depicting the fraction of the different types of regulatory changes occurring in the targets of the TFs (in x-axis) with a substantial number of changes in regulatory connections in correlation trend analysis in epithelial cells (B), myeloid cells (C), and T-NK cells (D). Across all cell types, the switching between activation and repression were predominant among all types of regulatory changes. **(E)** Changes in correlation distributions depicting the different types of regulatory changes occurring in the targets of the TFs NR2F1, HES2 and SOX9 in epithelial cells during cancer progression. The distributions in the left show the changes in correlation values of the targets of the TF NR2F1 that underwent switching from weak repression to weak activation; the distributions in the middle show the changes in correlation values of the targets of HES2 that underwent switching from weak activation to weak repression; and the distributions on the right show the changes in correlation values of the targets of SOX9 that underwent switching from weak activation to strong repression.

In epithelial cells, a large fraction of the transitions in the target genes of hub TFs was of switching types – from repression to activation and the vice-versa (Fig. 2B). For the TF NR2F1, ∼65% of the targets exhibited switching from repression to activation (Fig. 2B), of which ∼45%, ∼10%, ∼7% and ∼3% of the targets had switching from weak repression to weak activation (Fig. 2E), weak repression to strong activation, strong repression to strong activation, and strong repression to weak activation, respectively. In addition, ∼29% of the targets of NR2F1 showed switching from activation to repression, of which ∼18% had switching from weak activation to weak repression and ∼9% had switching from strong activation to weak repression (Fig. 2B). For HES2, ∼75% of the targets showed switching from weak activation to weak repression (Fig. 2B,E). For SOX9, ∼46% and ∼41% targets had switching from weak activation to weak repression and weak activation to strong repression respectively (Fig. 2B,E). The TF NR2F1 itself showed a small increase in expression with cancer progression (Fig. S9A), whereas the TFs HES2 and SOX9 did not show any change in expression with cancer progression (Fig. S9B,C).

Next, we analyzed the functions of the target genes that underwent different types of regulatory transitions, as this could reveal the functional transformation occurring through cancer progression. Several target genes of NR2F1 that transitioned from repression to activation were linked to cancer proliferation, progression, and metastasis. Among the target genes of NR2F1 that transitioned from repression to activation, ∼30% of the genes had known association with cancer progression or tumor suppression in different types of cancers. Among these, the *FSTL3* gene was linked to immune cell infiltration in LUAD and its high expression was associated with lower survival in LUAD patients^69^. In addition, the target gene *DIXDC1* was shown to increase invasion and migration ability of non-small-cell lung cancer cells^70^. Further, the target genes *ADAM9* and *PHF20L1* promoted cancer progression or metastasis^71,72^. On the other hand, the target *WWC3* repressed lung cancer invasion and metastasis^73^. Interestingly, one target gene, *TAF6L*, was associated with self-renewal of embryonic stem cells^74^.

Among the target genes that transitioned from activation to repression, ∼20% genes had known association with cancer progression or tumor suppression. *WDR54* has been reported as an oncogene in colorectal cancer^75^. The gene *JOSD2* has been shown to promote lung cancer^76^. Two other genes, *TMBIM6* and *PHLDB3*, were found to promote cancer growth and progression^77,78^. In addition, the gene *DNAJ4* suppressed EMT in nasopharyngeal carcinoma^79^ and the gene *CD82* was identified as a metastasis suppressor^80^.

Most of the targets of *HES2* underwent transitions from activation to repression. Among these targets, ∼42% of the genes were associated with cancer progression or tumor suppression. One gene, *TXNDC12*, was shown to promote EMT and metastasis in hepatocellular carcinoma^81^. Another gene, *PHACTR4*, was a tumor suppressor^82^. Taken together, these results showed that many of the target genes of NR2F1 and HES2 whose regulation changed with cancer stages were associated with cancer progression and metastasis.

In myeloid cells, ∼36%, ∼29% and ∼21% of the targets of CEBPD had switching from strong repression to weak activation, weak repression to weak activation, and strong repression to strong activation, respectively (Fig. 2C). For the TFs HIF1A, JUNB and MEF2C, ∼38%, ∼33% and ∼43% of their targets exhibited loss of repression, respectively (Fig. 2C). In T-NK cells, majority of the targets of the key TFs showed switching from repression to activation and the vice-versa (Fig. 2D). For JUND, ∼65% and ∼24% of the targets showed switching from weak repression to weak activation, and weak activation to weak repression, respectively (Fig. 2D).

In Myeloid cells, several target genes of the TF CEBPD that transitioned from repression to activation, including *REEP3*, *COA6*, *TFRC*, and *NRBF2*, were associated with promoting cancer progression, metastasis or chemoresistance^83-86^. In T-NK cells, among the target genes of the TF JUND that transitioned from activation to repression, *JAML* was involved in immune cell function and higher expression of this gene in tumor infiltrating immune cells was associated with better patient outcome^87^. Among the targets that underwent the repression to activation transition, *GOLM1* has been shown to suppress anti-tumor immunity^88^. *THEMIS2* - another target gene - affected functions of NK cells^89^. In addition, blocking signaling of the target gene *LAIR1* in immune cells inhibited tumor development^90^. Taken together, these results pointed to induction of immune suppression with cancer progression in the T-NK cells and an important role of the TF JUND in this process

### Cancer samples exhibit distinct gene regulation

We next focused on identifying TF-Target connections that may not show linear change in correlation with cancer progression, but may appear only in a subset of the cancer samples. These transitions may also have important implications for cancer and can lead to different trajectories of cancer progression in different patients. To identify sample-specific active GRNs, for each cell type in a sample, we considered those TF-target connections where the magnitude of the normalized correlation value was beyond mean ± 2×s.d. of the correlation values calculated across all connections in that cell type from all samples (Fig. S10). Comparison of overall degree distribution of TFs and targets revealed no substantial difference between cancer samples and normal samples except in myeloid cells, where the number of nodes with high degree decreased in the late-stage cancer samples (Fig. S11).

We then tested whether the cancer samples differ in gene regulation architecture compared to normal cells and whether the GRNs in cancer samples have unique active regulatory connections. To do so, we performed Principal Component Analysis (PCA) on all samples based on the normalized correlation values of the TF-target connections. PCA revealed separation of cancer and normal samples (Fig. 3A-C), despite the samples coming from patients carrying potentially different genetic changes. The separation between cancer and normal samples were clearer for myeloid and T-NK cells (Fig. 3B,C) compared to epithelial cells (Fig. 3A). These results suggested that the cancer samples contained unique TF-target connections that separated them from the normal cells, and despite the different patient origin, the cancer samples likely shared some common regulatory connections.

**Figure 3.**
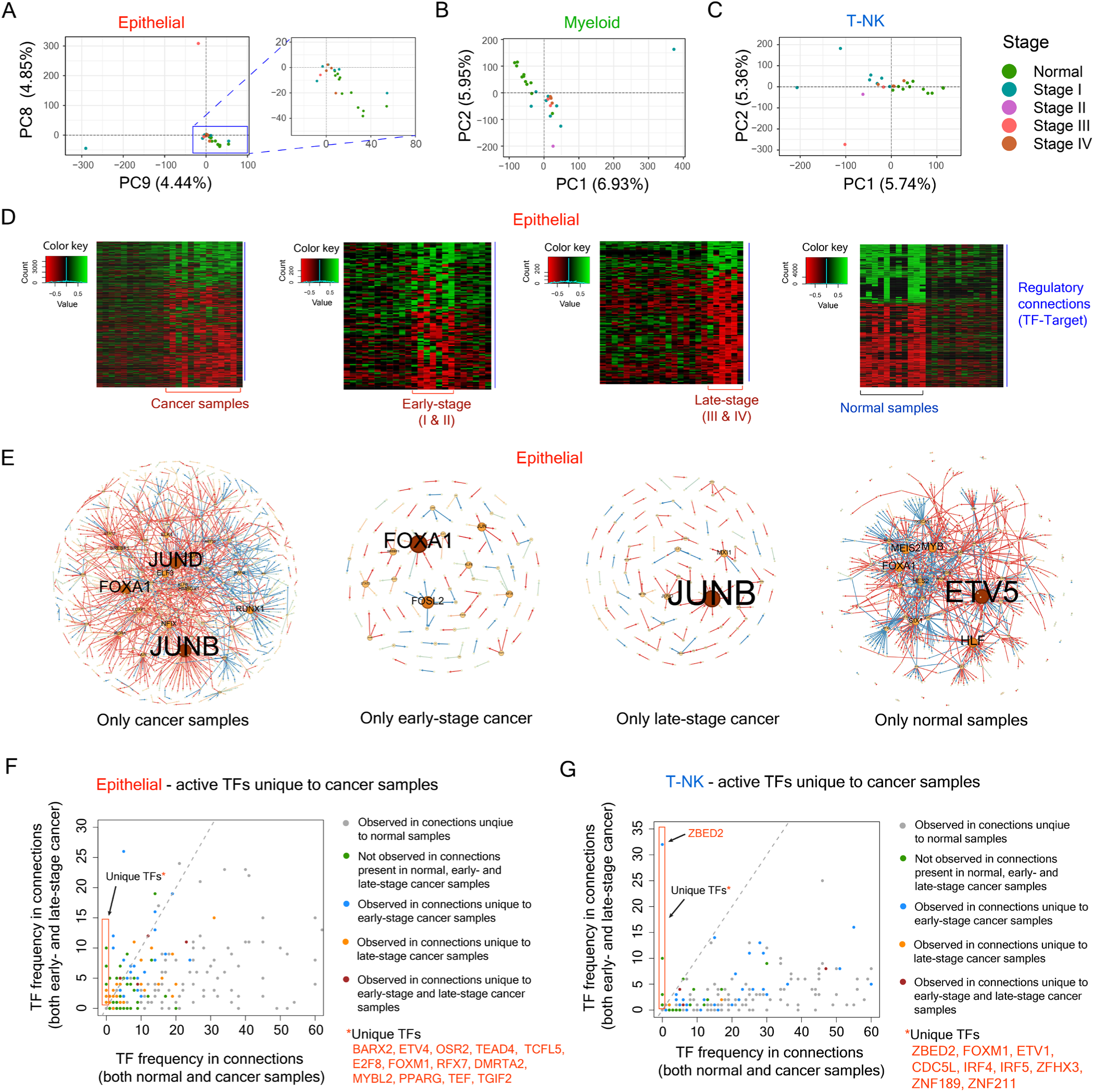
Cancer samples have early- and late-stage specific active regulatory connections and have TFs that are active only in cancer samples. **(A-C)** Principal Component Analysis (PCA) of all samples shows separation of normal and cancer samples in epithelial cells (A), myeloid cells (B) and T-NK cells (C). These results suggest that the cancer samples harbor unique regulatory connections compared to the normal samples. **(D)** From left to right, heatmaps showing the TF-target connections in epithelial cells that are active in only cancer samples (encompassing both early- and late-stage samples), only early-stage cancer samples, only late-stage cancer samples and only normal samples. Green color indicates positive TF-target correlation and red color indicates negative TF-target correlation. **(E)** Graphs showing the degree of genes (both TFs and targets) in epithelial cells that appear in the connections active in only cancer samples, only early-stage cancer samples, only late-stage cancer samples, and only normal samples. Red edges show negative TF-target correlation and blue edges show positive TF-target correlation. **(F-G)** Plot showing the frequency of TFs in connections that occur only in cancer samples and the frequency of the same TFs in connections that occur in both normal and cancer samples in epithelial cells (F) and in T-NK cells (G). Several TFs were only associated with connections that occurred in only normal samples and not in connections that appeared in both normal and cancer samples.

In the next step, we identified connections that were unique to cancer cells and were not observed in the normal cells, and vice-versa. To identify cancer-specific connections, we considered only those TF-target connections that were found in at least three cancer samples, but were not seen in any of the normal samples. In epithelial cells, we observed 1447 TF-target connections that were unique to cancer samples (Fig. 3D). In addition, 115 connections were unique to early-stage (stage I and II) cancer samples and 128 connections were unique to late-stage (stage III and IV) cancer samples (Fig. 3D). We also observed 1071 connections that were present only in normal samples and were not found in any of the cancer samples (Fig. 3D). We found similar unique connections in myeloid and T-NK cells (Fig. S12 and S13).

Next, we identified the TFs that were associated with the unique connections present in cancer samples. The TFs JUNB, JUND, FOXA1 and RUNX1 were overrepresented among the connections that were unique to cancer samples (Fig. 3E). In addition, the TFs FOXA1 and FOSL2 were overrepresented among the unique connections in early-stage cancer samples, whereas the TFs JUNB and MXI1 were overrepresented among the unique connections in late-stage cancer samples (Fig. 3E). The TFs ETV5, HLF, MYB, MEIS2 and FOXA1 were overrepresented among the connections that were unique to normal samples (Fig. 3E). We also identified similar overrepresented TFs in myeloid and T-NK cells (Fig. S14 and S15).

We further identified the TFs which were active only in cancer samples but not in normal samples, as these TFs could have very important role in driving cancer. We identified several such TFs among the connections that were unique to cancer samples. In epithelial cells, the unique TFs included BARX2, ETV4, and TEAD4 among others (Fig. 3F). The BARX2 gene has been shown to promote cancer progression and drive metabolic remodeling in LUAD^91^, even though it was considered a tumor suppressor gene in other cancer types^92^. TEAD4 was shown to drive EMT^93^ and ETV4 was found to promote stemness of breast cancer cells^94^. Similarly, we found a few TFs which were only active in normal epithelial cells (Fig. S16). We also observed unique TFs in cancer samples in myeloid and T-NK cells (Fig. 3G and Fig. S17, S18). Overexpression of the TF BNC2, that was unique to cancer samples in myeloid cells, in cancer stroma has been linked to promotion of EMT and invasion ^95^. ZBED2, one of the TFs unique to cancer samples in T-NK cells (Fig. 3G), was associated with dysfunction or exhaustion of CD8 T cells in melanoma^96^. FOXM1, another unique TF in T-NK cells, was reported to facilitate immune evasion in cancer by recruiting regulatory T cells^97^.

### Coordinated GRN transitions occur in different cell-types within the TME

Different cell types present in the tumor microenvironment (TME) have critical roles in regulating cancer progression and metastasis. For example, metabolic transformation of myeloid cells to a suppressive state can enable cancer progression^98^. Similarly, NK cells can control cancer growth and metastasis^99^. However, in the TME, interaction with cancer cells can transform NK cells to a metastasis promoting state^100^. Thus, cell-cell communication plays an important role in the transformation of the TME^101^.

Thus, to better understand the transformation of the TME, we focused on identifying the cooccurring regulatory changes that were happening between the Epithelial, Myeloid and T-NK cell types at different cancer stages. To do so, we tested for cooccurrence of specific TF-target connections between epithelial-myeloid and epithelial-T-NK cell types in early-stage and late-stage samples. We considered TF-target connections in two different cell types that showed same sign of correlation in at least three early-stage or late-stage samples. We then identified the TFs that were overrepresented in these connections in the two cell-types (see Methods, Fig. 4) and identified the changes in the targets of these TFs in the cooccurring TF-target connections (see Methods).

**Figure 4.**
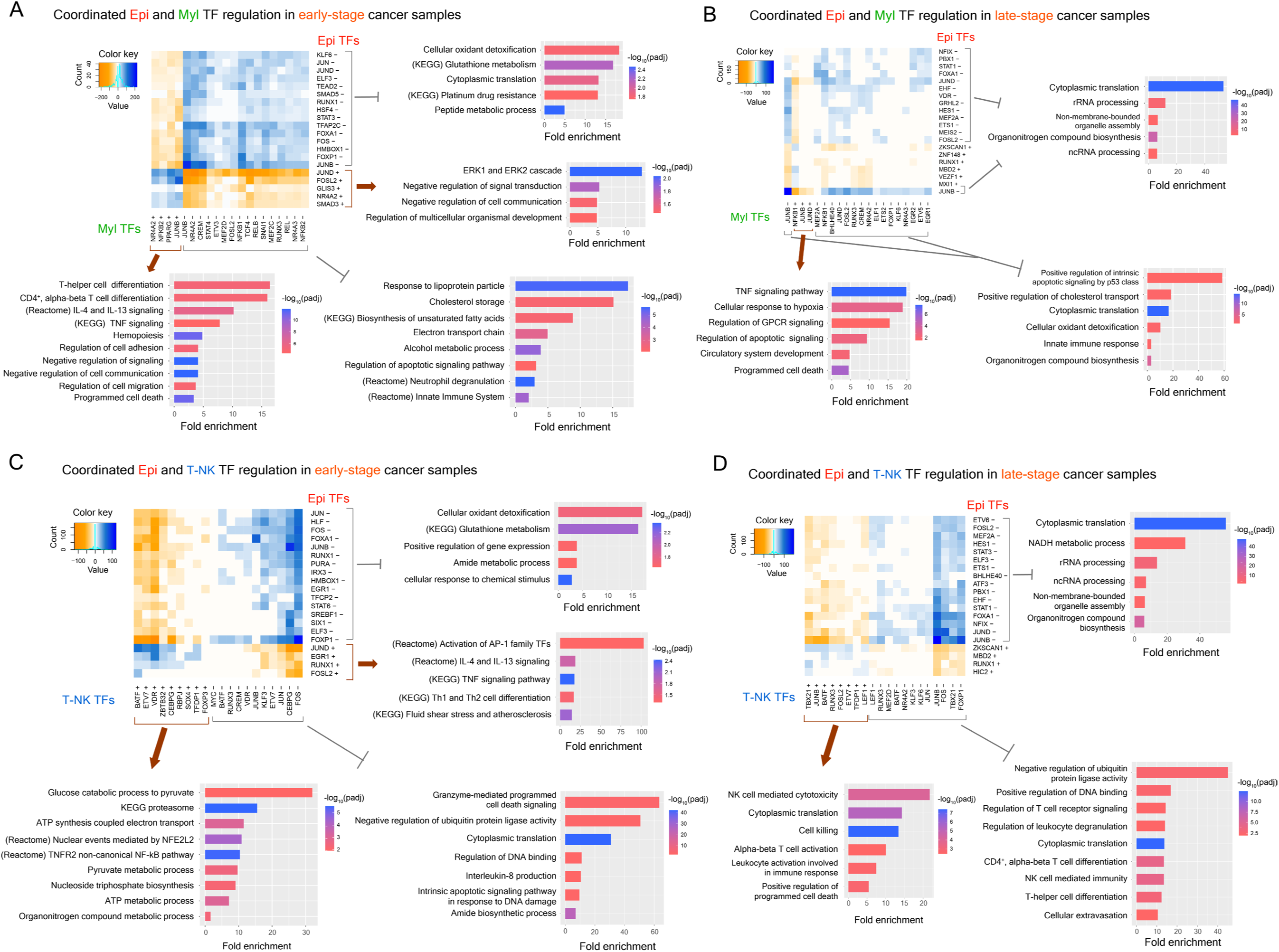
Coordinated changes in TF activities in different cell types transform the cancer microenvironment. **(A-B)** Top 20 TFs in epithelial and myeloid cells with coordinated regulatory activities in early-stage (A) and late-stage (B) cancer samples. **(C-D)** Top 20 TFs in epithelial and T-NK cells with coordinated regulatory activities in early-stage (C) and late-stage (D) cancer samples. The ‘+’ and ‘-’ signs indicate positive and negative correlation of the TFs with target genes. The functional enrichment analysis identified several enriched functional classes whose fold-enrichment and adjusted p-values (after multiple hypothesis testing correction) are shown. The orange color in heatmap indicates opposing signs of TF-target correlations for TFs in different cell types, where activation (repression) of targets by a TF in the first cell type cooccurred with repression (activation) of targets by another TF in the second cell type. The blue color in heatmap indicates same signs of TF-target correlations for TFs in different cell types, where activation (repression) of targets by a TF in the first cell type cooccurred with activation (repression) of targets by another TF in the second cell type.

For each cell-type pair, we identified the top 20 TFs from each cell type that showed coordinated activities at early- and late-stages of cancer (Fig. 4). We observed several TFs that showed coordinated activities between the epithelial and the myeloid cells in early-stage and late-stage samples (Fig. 4A,B), although the specific TFs were different between the early-stage and the late-stage samples (Fig. 4A,B). For epithelial-myeloid pair in early-stage samples, negative TF-target relations of 15 TFs and positive TF-target relations of five TFs in epithelial cells cooccurred with negative TF-target relations of 16 TFs and positive TF-target relations of 4 TFs in myeloid cells (Fig. 4A). In late-stage samples, negative TF-target relations of 14 TFs and positive TF-target relations of 6 TFs in epithelial cells cooccurred with negative TF-target relations of 17 TFs and positive TF-target relations of three TFs in myeloid cells (Fig. 4B). Similarly, for Epithelial - T-NK pair, in early-stage samples, negative TF-target relations of 16 TFs and positive TF-target relations of four TFs in epithelial cells cooccurred with negative TF-target relations of 11 TFs and positive TF-target relations of 9 TFs in T-NK cells (Fig. 4C). In late-stage samples, negative TF-target relations of 16 TFs and positive TF-target relations of 4 TFs in epithelial cells cooccurred with negative TF-target relations of 12 TFs and positive TF-target relations of 8 TFs in T-NK cells (Fig. 4D). We also identified several TFs that showed coordinated regulation between myeloid and T-NK cells (Fig. S19).

To further understand the coordinated functional changes happening across the three different cell-types, we performed functional enrichment analysis of the target genes that were part of the coordinated TF-target relations in each cell-type. As noted, negative relation between a TF and a target would indicate switch to repression or loss of activation, whereas positive relation would indicate switch to activation or loss of repression. For epithelial-myeloid pair, in early-stage samples, activation of negative regulation of cell-cell communication and signaling in epithelial cells cooccurred with activation of negative regulation of cell-cell communication and signaling in myeloid cells (Fig. 4A). In addition, this was accompanied by activation of cell adhesion and cell migration regulation in myeloid cells (Fig. 4A). In late-stage samples, repression of cytoplasmic translation in epithelial cells cooccurred with activation of TNF signaling and circulatory systems development, and repression of apoptotic signaling by p53 and innate immune response. TNF signaling has been reported to help accumulate myeloid derived suppressor cell accumulation^102^, which together with repression of immune response can facilitate cancer progression.

For epithelial – T-NK pair, in early-stage samples, activation of AP-1 family of TFs and fluid shear stress response in epithelial cells cooccurred with activation of electron transport, ATP and pyruvate metabolic process and repression of granzyme-mediated cell death signaling in T-NK cells (Fig. 4C). In late-stage samples, repression of cytoplasmic translation in epithelial cells cooccurred with activation of NK cell mediated cytotoxicity and repression of leukocyte degranulation and cellular extravasation in T-NK cells (Fig. 4D). Taken together, these results pointed to the coordinated nature of changes that occurred within the TME, and identified the TFs in different cell types that are the key regulators of these changes.

### Cancer samples show large-scale alterations in gene regulation dynamics

Gene expression happens in bursts in which a gene transitions between ON and OFF states, and mRNAs are produced in the ON state^103,104^ (Fig. 5A). TFs have a central role in regulation of transcriptional burst kinetics and can impact both the burst frequency (time interval between two consecutive bursting) and burst size (the number of mRNA molecules produced during the ON state)^105,27,28^. Thus, the burst parameters can change in cancer due to changes in activities of TFs. This can, in turn, influence the overall expression pattern of the target genes, as well as give rise to expression heterogeneity among cells within a tumor which can generate phenotypic plasticity.

**Figure 5.**
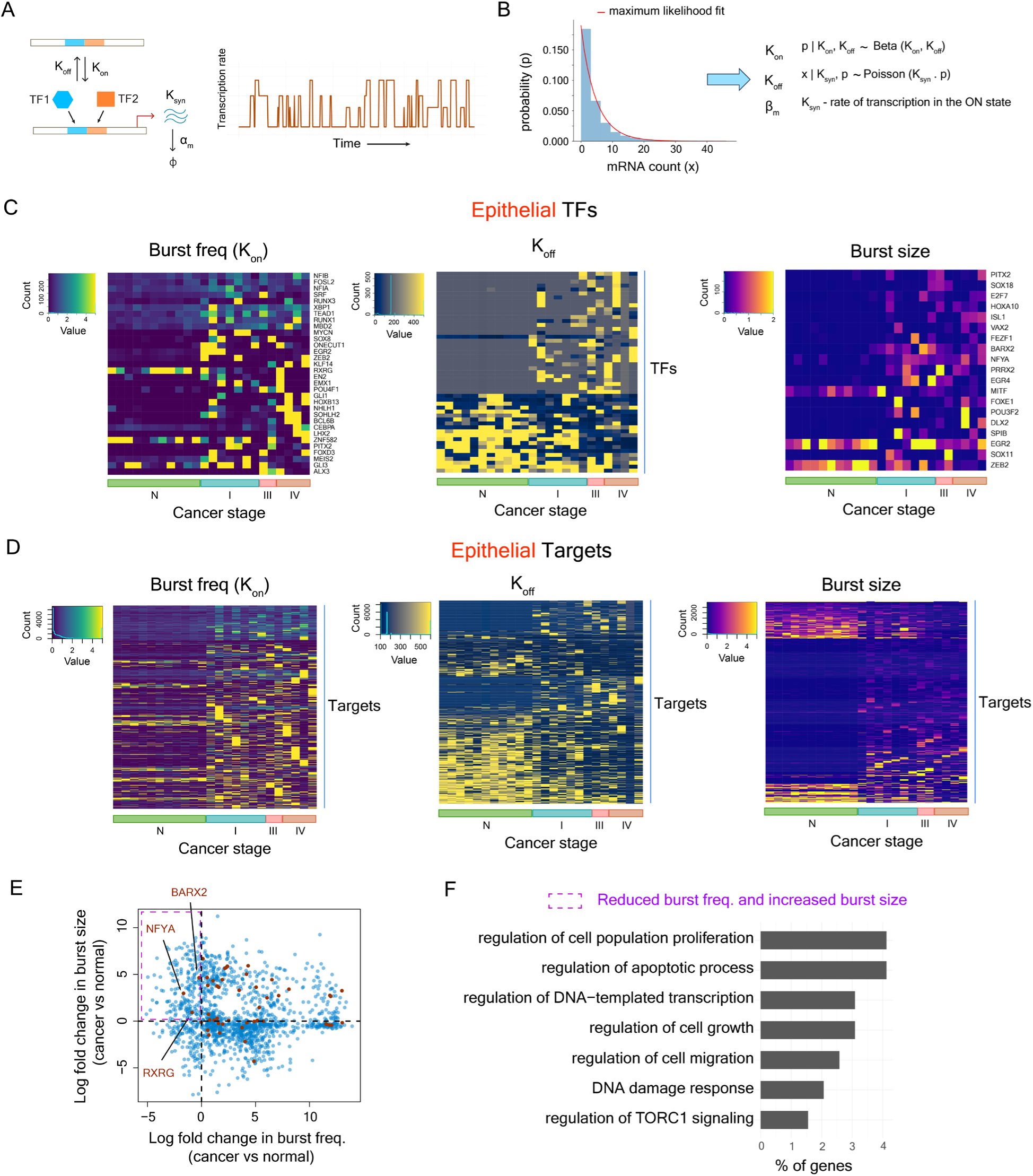
Cancer samples show widespread changes in gene regulation dynamics. **(A)** Two-state model of gene expression with a gene switching between ON and OFF states at the rates K_on_ and K_off_ respectively. **(B)** Estimation of burst parameters from single-cell RNA-seq data using a maximum likelihood approach described by Kim and Marioni (2013). **(C)** Heatmaps showing the changes in burst frequency (left), K_off_ (center), and burst size (right) in epithelial cells of the TFs with altered burst kinetics in cancer samples compared to normal samples. **(D)** Heatmap showing the changes in burst frequency (left), K_off_ (center), and burst size (right) in epithelial cells of the target genes with altered burst kinetics in cancer samples compared to normal samples. Only the target genes for which TF-target map was known were considered. **(E)** Log fold change in burst frequency vs log fold change in burst size in epithelial cells for all genes included in the analysis. **(F)** Functional association with biological processes in Gene Ontology (GO) of genes that showed decreased burst frequency and increased burst size in epithelial cells of cancer samples compared to normal epithelial cells.

To investigate the changes in gene regulation dynamics in cancer, we employed a two-state model of gene expression^104,106^ for all genes in all samples. In each sample, for each cell-type, we estimated the kinetic parameters of gene expression K_on_, K_off_ and K_syn_ from single-cell RNA-seq data using a maximum-likelihood approach^107^ (Fig. 5B). Changes in these parameters in cancer samples, indicated changes in the frequency of switching on, frequency of switching off and the mRNA synthesis rate in the ON state, respectively. We also calculated the parameter burst size that was represented by the ratio of K_syn_ to K_off_.

To identify genes with altered regulation dynamics in each cell type, we normalized the parameter distributions for each parameter across samples (Fig. S20-S28), classified genes into ‘low’ and ‘high’ categories, and identified genes undergoing low/intermediate-to-high or high/intermediate-to-low transitions from normal to cancer samples (see Methods). PCA analysis separated out normal and cancer samples for all cell types based on both burst frequency and burst size (Fig. S29-S31), suggesting alteration in gene regulation dynamics in all cell types of cancer samples compared to normal samples.

In epithelial cells, we identified 32 TFs with changes in burst frequency, of which 29 TFs showed low/intermediate to high transitions (Fig. 5C) and included the TFs TEAD1, XBP1, RUNX1, and ZEB2 (Fig. 5C; Fig. S32A). These TFs have been linked to cancer progression^108-111^. Three TFs (RXRG, GLI3 and ZNF582) showed high/intermediate to low burst frequency transitions (Fig. 5C). In addition, we identified 19 TFs in epithelial cells that showed changes in burst size in cancer cells compared to normal cells (Fig. 5C). Among these, BARX2 showed a low/intermediate to high burst size transition in cancer cells (Fig. 5C; Fig S32B) and ZEB2 showed a high/intermediate to low burst size transition (Fig. S32B). However, the TFs NR2F1, HES2, and SOX9, associated with the greatest number of regulatory transitions in correlation trend analysis in epithelial cells, did not show substantial change in burst frequency or burst size between normal and cancer samples (Fig. S33). In myeloid cells, 33 TFs showed changes in burst frequency and 52 TFs showed changes in burst size in cancer samples (Fig. S34). In T-NK cells, 30 TFs showed changes in burst frequency and 18 TFs showed changes in burst size (Fig. S35).

We also observed changes in burst frequency and burst size in many target genes in all cell types, irrespective of the changes in the burst parameters of their regulatory TFs. In epithelial cells, 755 targets showed changes in burst frequency and 428 targets showed changes in burst size (Fig. 5D). In myeloid cells, 740 targets showed changes in burst frequency and 1817 targets showed changes in burst size (Fig. S36). In T-NK cells, 799 targets showed changes in burst frequency and 417 targets showed changes in burst size (Fig. S37).

Genes with a decrease in burst frequency and no change in burst size or an increase in burst size would be more heterogeneous in their expression in a cell population^27^. In epithelial cells, we identified 194 genes whose burst frequency went down but burst size remained unchanged or increased in cancer samples. Among these, three TFs, namely, BARX2, NFYA and RXRG, were present (Fig. 5E). BARX2 has been associated with LUAD progression^91^, whereas NFYA promoted malignant behavior in breast cancer^112^. The TF RXRG, however, acts as a tumor suppressor in lung cancer^113^. In addition, several other genes in epithelial cells were associated with cell proliferation, migration, and DNA damage response (Fig. 5F). These results pointed to heterogeneous expression of genes that drive and suppress tumor progression.

We identified 585 genes in myeloid cells that showed lower burst frequency in cancer samples compared to normal and showed no change in burst size or an increase in burst size (Fig. S38A). Among these genes, several were associated with regulation of cell proliferation, cell migration and angiogenesis (Fig. S38B). In T-NK cells, we identified 281 genes that showed lower burst frequency in cancer samples compared to normal and showed no change in burst size or an increase in burst size (Fig. S39A). Several of these genes were associated with regulation of cell proliferation, DNA damage response, regulation of interleukin production, and inflammatory response (Fig. S39B). These results pointed to the heterogeneous nature of immune response in the tumor microenvironment.

### Integration of GRN topology and regulation dynamics enables identification of key transcriptional regulators

The changes in burst frequency and burst size involves changes in activities of the TFs that regulate the target genes. Therefore, we identified the TFs that were overrepresented among the genes showing altered regulation dynamics in all cell types. TFs that drive changes in both regulatory network architecture and regulation dynamics could be the most important drivers of cancer progression. To identify those, we calculated the degree of a TF by quantifying the number of its target genes that showed altered burst frequency or burst size. At the same time, we calculated degree of the same TF by quantifying the number of its target genes from correlation trend analysis (Fig. 1B), as well as from the number of targets identified in sample-specific GRN analysis (Fig. 3D,E). Several TFs showed high degree in both GRN topology and dynamics analysis (Fig. 6A), suggesting their central role in cancer progression.

**Figure 6.**
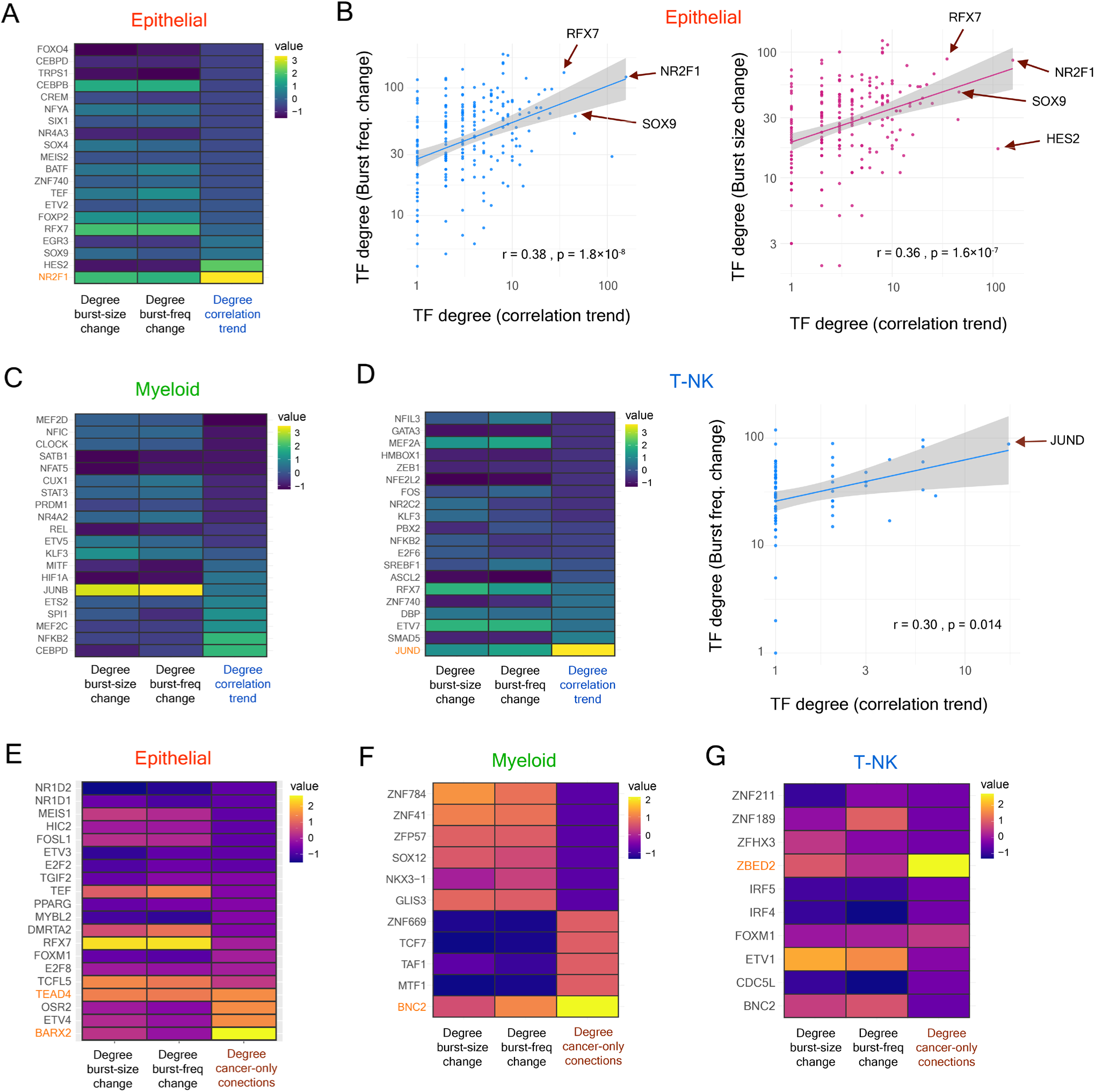
Integrated analysis of the changes in GRN architecture and dynamics identify the key TFs of cancer progression. **(A)** Heatmap showing the degree of the hub TFs identified from correlation trend analysis and the degree of the same TFs identified from the analysis of changes in burst frequency and burst size in epithelial cells. **(B)** Correlation between the degree of the hub TFs identified from the correlation trend analysis and TF degree identified from burst frequency change analysis in epithelial cells. **(C)** Heatmap showing the degree of the hub TFs identified from the correlation trend analysis and the degree of the same TFs identified from the analysis of changes in burst frequency and burst size in myeloid cells. **(D) Left -** Heatmap showing the degree of the hub TFs identified from correlation trend analysis and the degree of the same TFs identified from the analysis of changes in burst frequency and burst size in T-NK cells. **Right -** Correlation between TF degree identified from the correlation trend analysis and TF degree identified from burst frequency change analysis in T-NK cells. **(E-G)** Heatmap showing the degree of the TFs that were associated with TF-target connections unique to cancer samples and the degree of the same TFs identified from the analysis of changes in burst frequency and burst size in epithelial cells (E), in myeloid cells (F), and in T-NK cells (G).

In epithelial cells, the degree of the TFs calculated from the correlation trend analysis showed significant correlations with degree of the TFs calculated from the analysis of changes in burst frequency and burst size (Spearman’s correlation = 0.38, p = 1.8×10^-8^ for correlation with degree from burst frequency analysis and Spearman’s correlation = 0.36, p = 1.6×10^-7^ for correlation with degree from burst size analysis; Fig. 6B,C). Interestingly, in epithelial cells, the TF NR2F1, that emerged as a central TF from correlation trend analysis, also regulated many target genes that showed changes in their burst frequency and burst size (Fig. 6A-C). However, the TF HES2 regulated substantially lower number of target genes with altered regulation dynamics compared to the number of targets it regulated in the correlation trend analysis (Fig. 6B,C).

In myeloid cells, the TF CEBPD that had the highest degree in the correlation trend analysis, did not regulate as many genes with altered regulation dynamics (Fig. 6C). Instead, the TF JUNB regulated the greatest number of target genes with altered regulation dynamics. In addition, in myeloid cells, we did not see significant correlation between the TF degree calculated from the correlation trend analysis and TF degree calculated from burst frequency and burst size change analysis (Fig. S40A,B). In T-NK cells, the TF JUND, that emerged as the central TF in correlation trend analysis, also regulated the greatest number of target genes with altered regulation dynamics (Fig. 6D and S40C).

We also tested whether the TFs that were unique to the cancer samples had any role in regulating genes with altered regulation dynamics. Interestingly, in epithelial cells, the TFs BARX2, OSR2, and ETV4, that were overrepresented among the TF-target connections unique to cancer samples, also regulated many target genes with altered dynamics (Fig. 6E). Similarly, in myeloid cells, the TF BNC2 was overrepresented among the unique TF-target connections in cancer samples, and regulated many genes with altered dynamics (Fig. 6F). In T-NK cells, the TF ZBED2 regulated the greatest number of target genes among the TF-target connections unique to cancer samples and regulated many genes with altered dynamics (Fig. 6G). Taken together, these results identified several TFs that had a central role in not only in transition of the GRN architecture but also in changing gene regulation dynamics.

## Discussion

In summary, our work reveals the transitions in GRN architecture and gene regulation dynamics happening in epithelial, myeloid, and T-NK cells during the progression of lung adenocarcinoma. Our analysis identified several TFs that are central to the transitions of the regulatory network during cancer progression. These central TFs emerged despite the likely existence of genetic differences between the patient samples, suggesting that these TFs could have important implications in LUAD progression. We also showed that the central TFs are cell-type specific, thereby, underscoring the complex cellular changes occurring during cancer progression. We also identified several TF-target connections that were active only in cancer samples and identified TFs that are active only in cancer samples. These TFs are likely to be very important for cancer development and progression and need further investigation. In addition, these TFs are likely to be potentially interesting therapeutic targets as these have low activity in normal samples.

The transcription factor NR2F1 or COUP-TFI has been associated with cancer dormancy and only one study has so far has suggested its potential association with cancer migration and metastasis^66^. However, another member of the same family of TF, NR2F2 (or COUP-TFII) has been associated with cancer growth and metastasis^114,115^. COUP-TFII regulates angiogenesis, and COUP-TFII knockout could inhibit tumor growth and metastasis in mouse^114^. Further, expression of COUP-TFII gene in human lung cancer cells enhanced their invasive property^116^. Another central TF in the epithelial cells, HES2, is likely to be involved in developmental processes^117^ and has been suggested to have a tumor suppressive effect in neuroblastoma cells^68^. However, its role in cancer progression and metastasis in the context of lung adenocarcinoma remains to be elucidated.

We also identified different types of regulatory changes occurring in the tumor cells and the conflicting nature of some of these changes. For example, in epithelial cells the TF NR2F1 activated several genes that were known to promote cancer progression and metastasis in lung cancer and other cancer types. At the same time, NR2F1 also activated several genes that were known suppressors of cancer progression. These conflicting signals could be due to the complex changes occurring within the tumor microenvironment. Understanding this process better will require a deeper understanding of the molecular functions of the target genes, rather than their classification as cancer promoting or suppressing genes. Alternatively, it is also possible that a gene that act as a tumor suppressor in one cancer type can be a promoter of cancer progression in another cancer type, as evident from the existence of dual role TFs in cancer^118-120^. Finally, the conflicting signals might also represent the heterogeneous behavior of different subpopulations of cells in the tumor microenvironment which can eventually decide the fate of the tumor.

Our work also uncovered the coordinated activities of the TFs in different cell types within cancer samples and the simultaneous changes that were occurring within the tumor microenvironment. Here also we observed conflicting signals within one cell type in a cancer sample, again pointing towards the heterogeneous nature of the microenvironment.

Through an analysis of the changes in gene regulation dynamics, we identified genes that were becoming more heterogeneous in their expression with cancer progression. These genes also included several TFs that were associated with cancer progression or tumor suppression. In addition, several target genes with altered dynamics were also associated with tumor cell proliferation and migration. These results highlighted how activities of TFs can contribute to intratumor expression heterogeneity within populations of specific cell types, which can, in turn, generate intratumor phenotypic heterogeneity.

Taken together, our work highlights the large-scale changes in gene regulation taking place in the tumor microenvironment during cancer progression and uncovers the central transcription factors in this process. In this regard, one of the key unresolved questions is how the TFs NR2F1 (COUP-TFI) and SOX9, associated with cancer dormancy in earlier studies, influence LUAD progression. Although dormancy has been linked to drug tolerance, and metastasis in cancer^121,122^, more work is needed to understand whether dormancy itself can facilitate the cancer cell migration process. Alternatively, it is possible that these TFs could facilitate or drive cancer progression and metastasis in lung adenocarcinoma, which needs to be tested in future work. Finally, we note that the central TFs could vary between different patient cohorts and may also depend on the genetic background of the patients. Thus, analysis of more patient data of diverse genetic origin and at different stages of cancer will enable us to do a more robust identification of key TFs in cancer.

## Materials and Methods

### Dataset

The single-cell RNA-seq data for lung adenocarcinoma patients were obtained from (NCBI GEO GSE131907)^123^. Specifically, 26 samples from patients of different cancer stages were used, including data from 11 samples from adjacent normal lung tissue collected alongside tumor samples, seven stage I samples, one stage II sample, two stage III samples, and four stage IV samples (Table S4). Metastatic samples in other organs (e.g., lymph node or brain) were not considered in our analysis, as differences in tissue environment could activate specific gene regulation pathways and could confound our analysis. The dataset contained data for seven cell types, namely, epithelial, endothelial, fibroblast, mast, myeloid, B and T-NK cells. In addition, a set of cells in the dataset were unannotated. For the unannotated cell types, Seurat v4^124^ and SciBet^125^ were used for cell type annotation. For SciBet, the annotated cells in the dataset were used as the training set to predict the cell-types of the unannotated cells. Cells that were assigned same cell type from both these methods were included in the analysis of which 98.3% were annotated as T-NK cells (Fig. S41). Cell-types for which at least 100 cells were present in all samples were considered for gene regulatory network (GRN) analysis, as it was essential for a robust reconstruction of the GRN architecture and estimation of regulation dynamics. Three cell types - epithelial, myeloid, and T-NK cells – satisfied this criterion and were considered for all subsequent analyses. All analyses were performed individually for the three cell types to obtain a more nuanced view of the cell-type specific regulatory transitions occurring in the tumor microenvironment.

### Calculation of pairwise expression correlation

For identification of the active GRN, pairwise expression correlation of all genes in each cell-type within individual samples was calculated. The bigScale2 pipeline^55^ was used for this purpose and centered correlation values were calculated (Supplementary file 1). In epithelial cells, the correlation values for the LUNG_T08 sample (stage I) could not be calculated. In addition, the distribution of the correlation values in the LUNG_T09 sample (stage II) showed almost a uniform distribution for correlation values between -1.0 and 0.5 and with peaks above correlation value of 0.5. This suggested that in this sample, the correlation calculation was biased by the low cell number (119 cells). Pairwise correlation values were likely to peak around zero due to presence of correlation values between many genes that were not targets of each other and hence, were unlikely to show high correlations. Thus, this sample was discarded from all subsequent analysis of GRN architecture. For myeloid and T-NK cells, no such issue was observed (Supplementary file 2 and 3) and all samples were considered for GRN analysis.

As expression correlation may be present between genes without any direct regulation, only high confidence TF-target connections were considered. To do so, a two-step filtering process was followed. In the first step, the experimentally validated TF-target connections were obtained from various studies and databases including^126-132^. In the second step, a search was carried out for known binding sites of the TFs in the promoter regions of the target genes. To do so, non-redundant binding motifs in the form of position weighted matrices (PWM) for the human TFs were collected from hocomoco (v11)^133^ and JASPAR (2020)^134^ databases. All possible combinations of nucleotides from the PWMs for a TF were considered for searching in the region from 1000bp upstream of the start codon to 100bp downstream of the start codon. Sequences containing up to two mutations compared to the search sequence were considered as TF binding sites. Among all experimentally validated TF-target connections, only those with at least one mapped TF binding site were considered for all downstream analyses. The correlation values corresponding to only these TF-target connections were considered for the analysis of GRN architecture. For each cell-types, the correlation distributions of the samples were normalized by quantile normalization method using *normalize.quantiles* function from the R package ‘*preprocessCore’* ^135^. The correlation distributions before and after normalization were visualized using boxplots created with the R package *‘ggplot2’* ^136^ (Fig. S1-S3).

### Correlation trend analysis

The TF-target connections undergoing transitions during cancer progression were identified through a correlation trend analysis. Specifically, for every filtered TF-target connection in a cell-type, a linear regression model was fit to the normalized correlation values (y-axis) across the cancer stages (x-axis). The slope of the regression line and the goodness-of-fit (R^2^) were calculated for each regression model. The TF-target connections with at least two finite correlation values in adjacent normal samples, at least two finite correlation values in stage I samples, at least one finite correlation value in stage II/stage III samples and at least two finite correlation values in stage IV samples were considered. This was done to ensure that the results from the correlation trend analysis were robust and the high goodness-of-fit (R^2^) value did not arise merely due to a low number of finite correlation values in the samples.

Among the TF-target connections which satisfied this criterion, the ones with R^2^ values greater than or equal to 0.5 (Fig. S4) and with the regression slope value beyond the mean ± 2×s.d. of all slope values, calculated for all TF-target connections in that cell type across all samples (Fig. S5), were taken forward for the next steps. This enabled identification of TF-target regulations undergoing the most substantial transitions with cancer progression. For these connections, the correlation heatmaps with mean correlation across different cancer stages was generated with the *heatmap.2* function from the *‘gplots’* package in R^137^ (Fig. 1B, S6A, S7A). The identified TF-target connections were then used to calculate indegree and outdegree of all the TFs and targets. The TFs with the most outdegree (and total degree) represented the hub TFs for cancer progression, as these TFs showed the most widespread changes in regulation of their targets. The network representations of the TF-target connections undergoing transitions were generated using Gephi (https://gephi.org/) (Fig. 1C-E), where the nodes represented genes and the edges represented regulation between genes. The size of a node was proportional to the total degree of the node, and the thickness of an edge represented the magnitude of the slope of the fitted regression line between the two genes (TF-target), thereby, showing the magnitude of the change in correlation pattern with cancer progression. A positive slope, represented by the blue color, indicated an increase in correlation with tumor progression. Conversely, a negative slope, represented by the red color, indicated a decrease in correlation with tumor progression. In addition, hub TFs did not merely emerge because of their overrepresentation in our TF-target maps (Fig. S42).

### Inferring the type of regulatory change from correlation trends

Changes in the expression correlation during cancer progression can arise due to different types of changes in the regulation of a target genes by a TFs. For example, a positive correlation trend (thus, a positive slope value) can arise due to emergence of a new activation, a loss of repression or a switching from a repression to activation. All these transitions will lead to an increase in expression correlation value with cancer progression. Thus, the TF-target connections identified from the correlation trend analysis were further analyzed to infer the type of regulatory change occurring during cancer progression. This was essential to better understand the functional transformation occurring in different cell types in the tumor microenvironment.

The regulation changes were classified into the following classes: gain of activation (weak /strong), gain of repression (weak /strong), loss of activation (weak /strong), loss of repression (weak /strong), switching from activation (weak /strong) to repression (weak /strong) and switching from repression (weak /strong) to activation (weak/strong). As activation and repression were further subclassified into weak or strong types, this generated a total of 16 types of regulatory transitions (Fig. 2A-D). The mean correlation values in the normal samples and in the stage IV samples, as well as the slope of the regression line from the correlation trend analysis were considered for classification of TF-target connections into different types of regulatory transitions. The slope values of 0.2 and -0.2 were set as thresholds for classifying strong or weak transitions (Fig. S43).

A TF-target connection was considered to have undergone a gain of activation if the mean correlation value for this pair was non-negative in the normal samples (mean_corr_normal_ ≥ 0) and the slope of the regression line was positive (slope_regr_ > 0). It was defined as a gain of strong activation if slope_regr_ > 0.2, and as a gain of weak activation if 0 < slope_regr_ ≤ 0.2. A TF-target connection was considered to have undergone a gain of repression if mean_corr_normal_ ≤ 0 and slope_regr_ < 0. It was defined as a gain of strong repression if slope_regr_ < -0.2 and as a gain of weak repression if -0.2 ≤ slope_regr_ < 0.

A TF-target connection was considered to have undergone a loss of strong activation if, for this pair, mean_corr_normal_ ≥ 0, mean_corr_stageIV_ ≥ 0, and slope_regr_ < -0.2. A TF-target connection was considered to have undergone a loss of weak activation if mean_corr_normal_ ≥ 0, mean_corr_stageIV_ ≥ 0, and -0.2 ≤ slope_regr_ < 0. Similarly, a TF-target connection was considered to have undergone a loss of strong repression if mean_corr_normal_ ≤ 0, mean_corr_stageIV_ ≤ 0, and slope_regr_ > 0.2. A TF-target connection was considered to have undergone a loss of weak repression if mean_corr_normal_ ≤ 0, mean_corr_stageIV_ ≤ 0, and 0 < slope_regr_ ≤ 0.2.

A TF-target pair was considered to have switched from strong activation to strong repression if mean_corr_normal_ > 0.6, mean_corr_stageIV_ < 0 and slope_regr_ < -0.2. A TF-target pair was considered to have switched from strong activation to weak repression if mean_corr_normal_ > 0.6, mean_corr_stageIV_ < 0 and -0.2 ≤ slope_regr_ < 0. A TF-target pair was considered to have switched from weak activation to strong repression, if 0<mean_corr_normal_ ≤ 0.6, mean_corr_stageIV_ < 0, and slope_regr_ < -0.2. A TF-target pair was considered to have switched from weak activation to weak repression if 0<mean_corr_normal_ ≤ 0.6, mean_corr_stageIV_ < 0, and -0.2 ≤ slope_regr_ < 0.

Conversely, a TF-target pair was considered to have switched from strong repression to strong activation if mean_corr_normal_ < -0.6, mean_corr_stageIV_ > 0 and slope_regr_ > 0.2. A TF-target pair was considered to have switched from strong repression to weak activation if mean_corr_normal_ < -0.6, mean_corr_stageIV_ > 0 and 0 < slope_regr_ ≤ 0.2. Further, a TF-target pair was considered to have switched from weak repression to strong activation if -0.6 ≤ mean_corr_normal_ < 0, mean_corr_stageIV_ > 0 and slope_regr_ > 0.2. A TF-target pair was considered to have switched from weak repression to weak activation if -0.6 ≤ mean_corr_normal_ < 0, mean_corr_stageIV_ > 0 and 0 < slope_regr_ ≤ 0.2.

### Identifying sample- and stage-specific gene regulations in cancer

TF-target correlation trend analysis might not be able to identify all regulatory connections that are likely to be important for cancer, as some of the TF-target connections could be active only in some stages during cancer progression and may not be present in other stages. In addition, some of the TF-target connections could be active only in a subset of samples, but nevertheless could have important role in cancer progression.

To identify sample- and stage-specific TF-target regulations during cancer progression, sample-specific active GRNs were identified for each cell type. First, for each cell type, the correlation values of all TF-target connections across all samples were considered and the mean as well as the standard deviation (s.d.) were calculated (Fig. S10). Next, for a cell type within each sample, the TF-target connections for which the correlation values were beyond the mean ± 2 s.d. values of that cell type were identified and considered as part of the active GRN. This process was repeated for all three cell types across all samples.

To test whether the normal and cancer samples contain unique active GRNs, for each cell type, principal component analysis (PCA) on all samples was performed using the R package *‘PCAtools’* ^138^(Fig. 3A). In the next step, the TF-target connections unique to normal samples, cancer samples, early-stage (stage I & II) cancer samples, and late-stage (stage III & IV) cancer samples were identified. A TF-target connection was unique to cancer samples if the connection was active in at least three cancer samples spanning early-stage (stage I & II) and late-stage (stage III & IV) samples, but not in any normal sample. A TF-target connection was considered unique to early-stage cancer samples if the connection was active in at least three early-stage (stage I & II) cancer samples and not in any normal sample or late-stage cancer sample. A TF-target connections was considered unique to late-stage cancer samples if the connection was active in at least three late-stage (stage III & IV) cancer samples and not in any normal sample or early-stage cancer sample. Similarly, a TF-target connection was considered unique to normal samples if the connection was active in at least three normal samples and not in any cancer sample.

Considering all unique TF-target connections in normal and cancer samples, the indegree, outdegree and total degree of each gene were calculated. TFs that showed zero degree in normal samples but non-zero degree in cancer samples were identified as cancer-specific active TFs. Similarly, TFs that showed zero degree in cancer samples but non-zero degree in normal samples were identified as normal sample-specific active TFs.

### Coordinated changes in gene regulation across cell types

The tumor microenvironment contains different cell types and the phenotypic transformation of the tumor is driven by molecular changes occurring across these different cell types. Molecular changes, driven by altered gene regulation, in one cell type can facilitate or inhibit molecular and regulatory changes in other cell types within the tumor microenvironment. A better understanding of these coordinated changes in gene regulation, and eventually cellular phenotypes, with cancer progression can provide deeper insights into cancer transformation.

The coordinated changes occurring across different cell types in the tumor microenvironment were investigated through a comparison of regulatory changes occurring between Epithelial-Myeloid cells and Epithelial-T-NK cells across all samples. To start with, the cooccurrence of a pair of TF-target connections between a pair of cell types within a sample were identified. Next, the frequency of such pairs that occurred in more than one sample between the same pair of cell types and with the same correlation signs were counted. Subsequently, the pairs of connections were marked for stage-specific cooccurrences where at least three samples in that stage showed cooccurrence of these connections and no sample in other stages showed cooccurrence. This led to identification of TF-target pairs between cell types that cooccurred only in normal samples, only in early-stage cancer samples and only in late-stage cancer samples. Finally, the overrepresented TFs were identified and the frequency of their connections to targets with positive and negative correlation values were counted. The top 20 overrepresented TF pairs along with their correlation signs with their targets for every pair of cell type at early- and late-cancer stages were represented by heatmaps (Fig. 4).

To further understand how coordinated changes in gene regulation across different cell types influence the cancer microenvironment and affect cancer progression, functional enrichment analysis of the target genes of the top 20 TFs, identified from the previous step, was performed. The target genes that showed positive correlation and negative correlation with a TF were analyzed separately. The functional enrichment analysis was done using ‘*gprofiler*’ ^139^ and the functional enrichment for Biological Processes in Gene Ontology (GO) database^140,141^, KEGG pathways^142,143^ and Reactome pathways^144^ were performed. The significantly enriched classes were obtained by filtering classes with adjusted p-value less than 0.1 (10% False Discovery Rate) and their fold enrichment values were calculated.

### Inferring gene regulation dynamics from single-cell RNA-seq data

The two-state model of gene expression enabled estimation of the parameters of transcriptional bursts from single-cell RNA-sequencing data using a maximum likelihood approach, as shown by Kim and Marioni, 2013^107^. According to the model, a gene transitions into ON and OFF states with the rates K_on_ and K_off_ respectively. Transitions to the ON state results in the production of mRNA at a rate K_syn_. These mRNA molecules undergo removal at a constant rate of K_deg_.

Specifically,

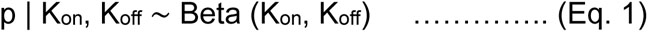

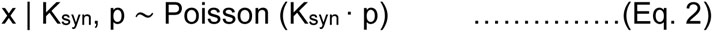

where ‘x’ is the number of RNA transcripts of a gene within a cell and ‘p’ is an auxiliary variable that follows a beta distribution. The mean of the parameter ‘p’ is equal to the fraction of time that the gene spends in the active state. The resultant marginal distribution is known as Poisson-Beta distribution, denoted by P (x | K_on_, K_off_, K_syn_), and gives the steady-state distribution for the mRNA copy numbers across cells. All parameters are further normalized by K_deg_. The parameters K_on_ and K_off_ control the shape of the Beta distribution and represent the probability of a gene being in ON and OFF states, respectively. K_syn_ is the mean of Poisson distribution and represents the average rate of gene expression when the gene is in the ON state.

The single cell RNA-seq dataset for each cell type in each sample was given as the input to the model. The two-state model parameters 𝜃 = (𝐾_𝑜n_, 𝐾_𝑜ff_, K_syn_) were inferred by maximizing the likelihood of the parameters of Poisson-Beta distribution for a given set of observed data points X.

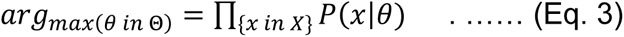

This was further represented as maximization of the log-likelihood, which was equivalent to minimization of the negative log-likelihood:

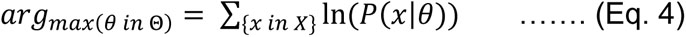

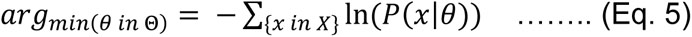

The negative log-likelihood was minimized using L-BFGS-B approach^145^, and the resulting distributions for parameters in 𝜃 were calculated.

### Identifying TFs associated with altered regulation dynamics

Analysis of gene regulation dynamics provides insights into the dynamic changes in gene regulation occurring with cancer progression and enable a better understanding the emergence of gene expression heterogeneity and consequently, phenotypic heterogeneity. To identify genes with altered regulation dynamics, the following steps were followed. In the first step, for each cell type, the distributions of each burst parameter (K_on_, K_off_ and burst size) for all genes were obtained across all samples. The distributions of each parameter were normalized separately with the quantile normalization method using the R package ‘*preprocessCore*’ ^135^. Next, in each sample and for each parameter, a gene was classified into either ‘low’ or ‘high’ category if the value of the parameter for that gene was among the bottom 25% or top 25% values of the parameter distribution, respectively. Subsequently, for a specific burst parameter, a gene was identified as transitioning to ‘high’ category from normal to cancer stage if the parameter value was classified as ‘high’ in at least three cancer samples and in none of the normal samples. Similarly, a gene was identified as transitioning to ‘low’ category, if the parameter value was classified as ‘low’ in at least three cancer samples and in none of the normal samples. For the analysis of target genes, only the targets with at least one regulatory TF in our TF-target were considered for analyses, as the eventual goal was to identify the TFs driving the dynamic changes in regulation of these target genes.

A reduction in burst frequency, with or without an increase in burst size, can lead to increased gene expression heterogeneity^27^. To identify genes that showed a reduction in burst frequency with or without an increase in burst size in cancer samples compared to normal samples, the parameters log fold change in burst frequency and log fold change in burst size were defined. The log fold change in burst frequency was calculated as the log_2_(mean burst frequency in cancer samples/mean burst frequency in normal samples). Similarly, the log fold change in burst size was calculated as the log_2_(mean burst size in cancer samples/mean burst size in normal samples). Genes with log fold change in burst frequency < 0 and log fold change in burst size ≥ 0 were identified as genes with increased expression heterogeneity in cancer samples and their association with specific biological processes in Gene Ontology (GO) database were investigated.

### Integration of GRN architecture and dynamics analysis

Results from the analyses of GRN architecture and regulation dynamics were integrated to identify the TFs that affect both. For integration, the following steps were followed. From the TF-target connections undergoing transitions in the correlation trend analysis, the degrees of the TFs were obtained. A higher degree suggested association with more transitioning regulatory connections, thus, implying a central role in the remodeling of GRN architecture during cancer progression. In addition, the degree values of the TFs that were identified to be active in only cancer samples in the sample- and stage-specific analysis of GRN architecture were also considered.

From the analysis of changes in gene regulation dynamics, the genes with a change in burst frequency or burst size or both were considered. For genes with a change in burst frequency, the TFs that regulate these genes were identified from our TF-target map and the degree of these TFs were calculated. A higher degree suggested association with changes in burst frequency of more target genes, and thus, a central role in changing regulation dynamics in cancer. Similarly, the degree values of the TFs regulating the genes with altered burst size were calculated.

The degrees of the TFs obtained from the analysis of the correlation trend analysis were compared with the degrees of the TFs calculated from the analysis of the GRN dynamics. The degree values of the TFs obtained from the analysis of correlation trend, as well as from the analysis of cancer-specific active GRN, along with the degree values obtained from the analysis of changes in burst frequency and burst size were represented through heatmaps. In addition, the spearman’s correlation between the degree values were calculated and the TFs showing high degree values in both analyses were identified as the key TFs.

## Author contributions

Conceptualization: RD

Methodology: UR, AS, DS, RD

Investigation: UR, AS, DS, RD

Visualization: UR, AS, DS, RD

Funding acquisition: RD

Project administration: RD

Supervision: RD

Writing – original draft: UR, AS, DS, RD

Writing – review & editing: UR, AS, DS, RD

## Data availability

All datasets used in the manuscript are publicly available.

## Code availability

All codes for data analysis and models are available in github: https://github.com/riddhimandhar/GRN_LUAD

## Supplementary material

### Supplementary figures

Figure S1 to S43

### Supplementary tables

**Table S1:** Degree values of the genes associated with the TF-target connections that underwent regulatory transitions with cancer progression in epithelial cells and were identified from correlation trend analysis.

**Table S2:** Degree values of the genes associated with the TF-target connections that underwent regulatory transitions with cancer progression in myeloid cells and were identified from correlation trend analysis.

**Table S3:** Degree values of the genes associated with the TF-target connections that underwent regulatory transitions with cancer progression in T-NK cells and were identified from correlation trend analysis.

**Table S4:** Patient sample information

### Supplementary files

Supplementary file 1: Histograms showing gene expression correlation values between all gene pairs in epithelial cells.

Supplementary file 2: Histograms showing gene expression correlation values between all gene pairs in myeloid cells.

Supplementary file 3: Histograms showing gene expression correlation values between all gene pairs in T-NK cells.

