## Supplementary figures for "Gene regulatory network transitions reveal the central transcription factors in lung adenocarcinoma progression"

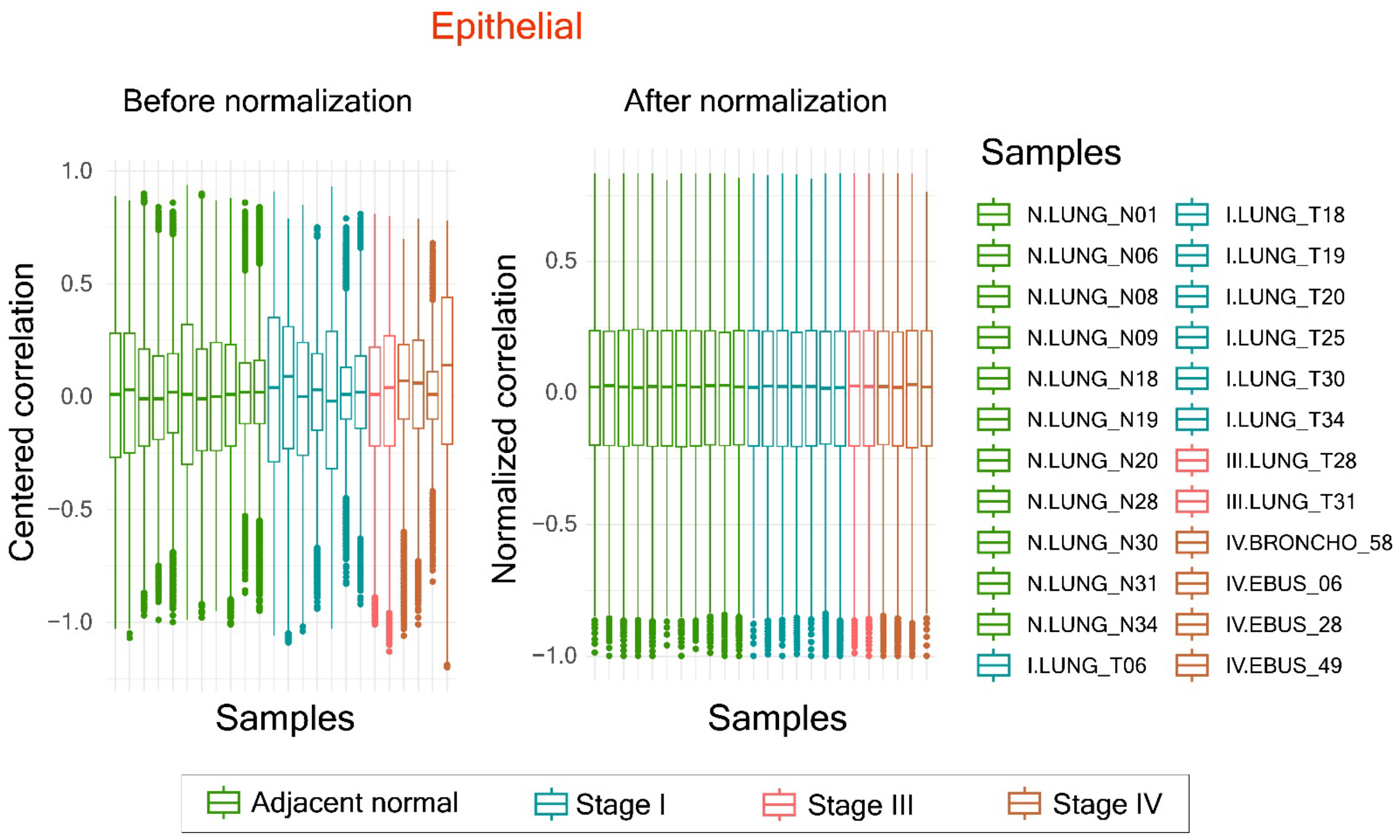


**Figure S1 –** Boxplots showing distribution of centered correlation values (obtained from the bigScale2 pipeline) before normalization (left) and the normalized correlation values obtained after normalization in epithelial cells. The colors indicate the cancer stages.

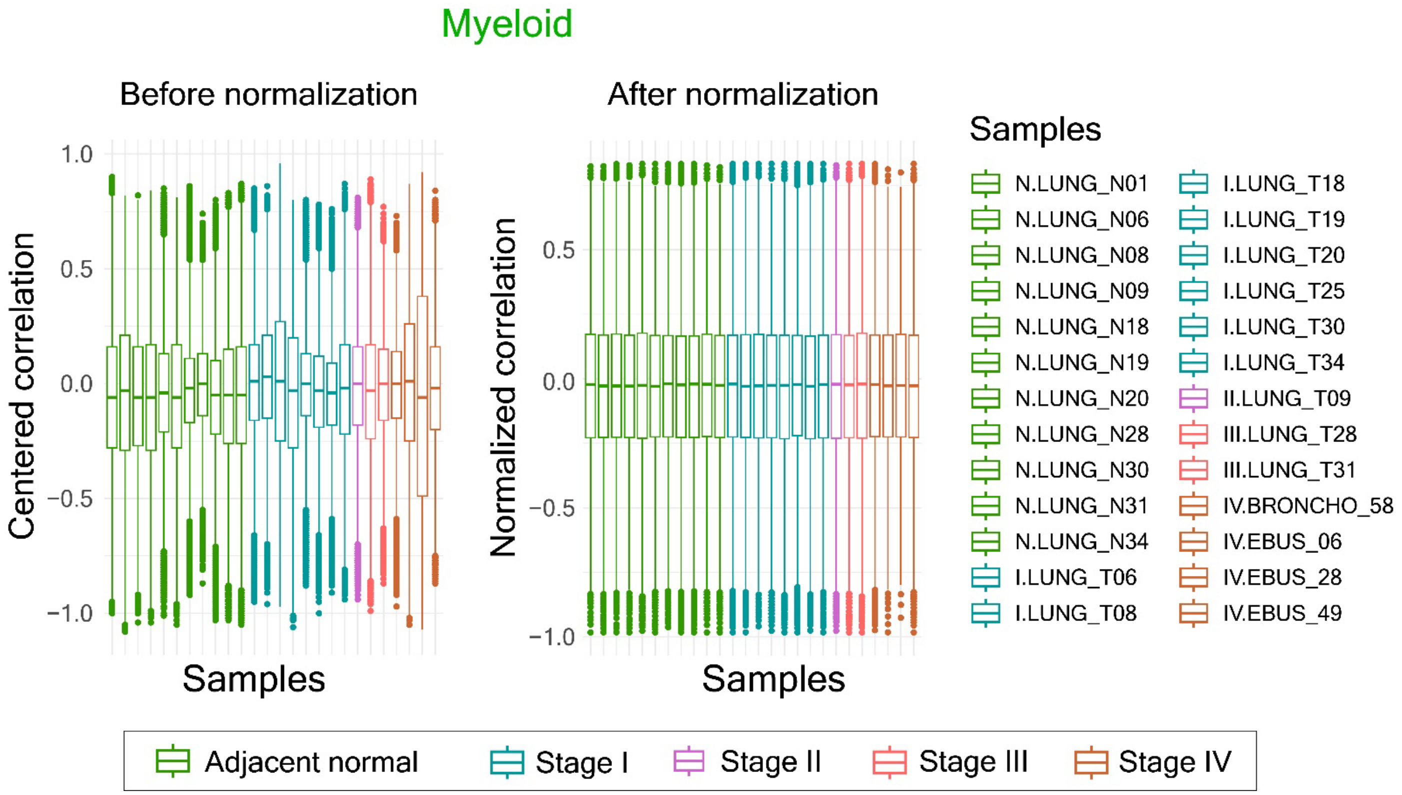


**Figure S2 -** Boxplots showing distribution of centered correlation values (obtained from the bigScale2 pipeline) before normalization (left) and the normalized correlation values obtained after normalization in myeloid cells. The colors indicate the cancer stages.

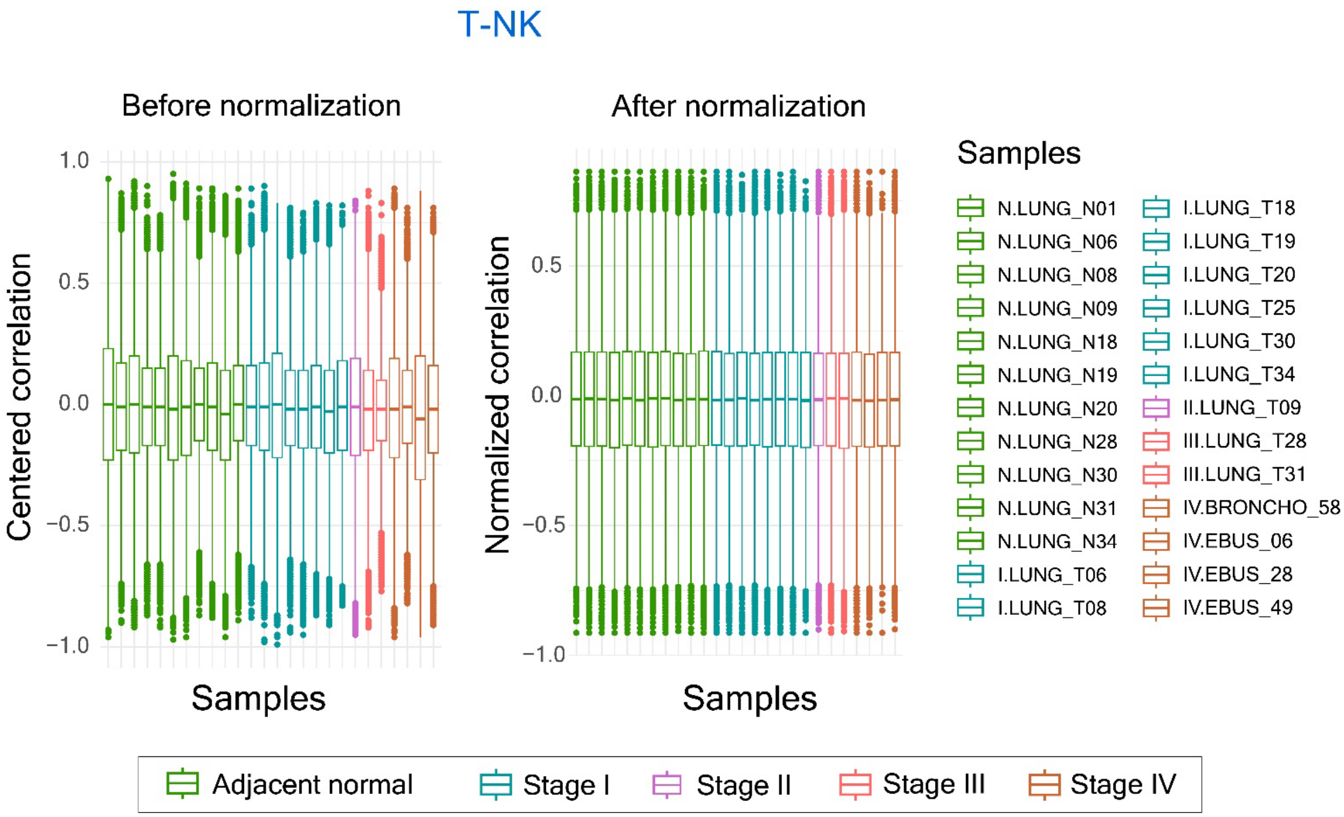


**Figure S3 -** Boxplots showing distribution of centered correlation values (obtained from the bigScale2 pipeline) before normalization (left) and the normalized correlation values obtained after normalization in T-NK cells. The colors indicate the cancer stages.

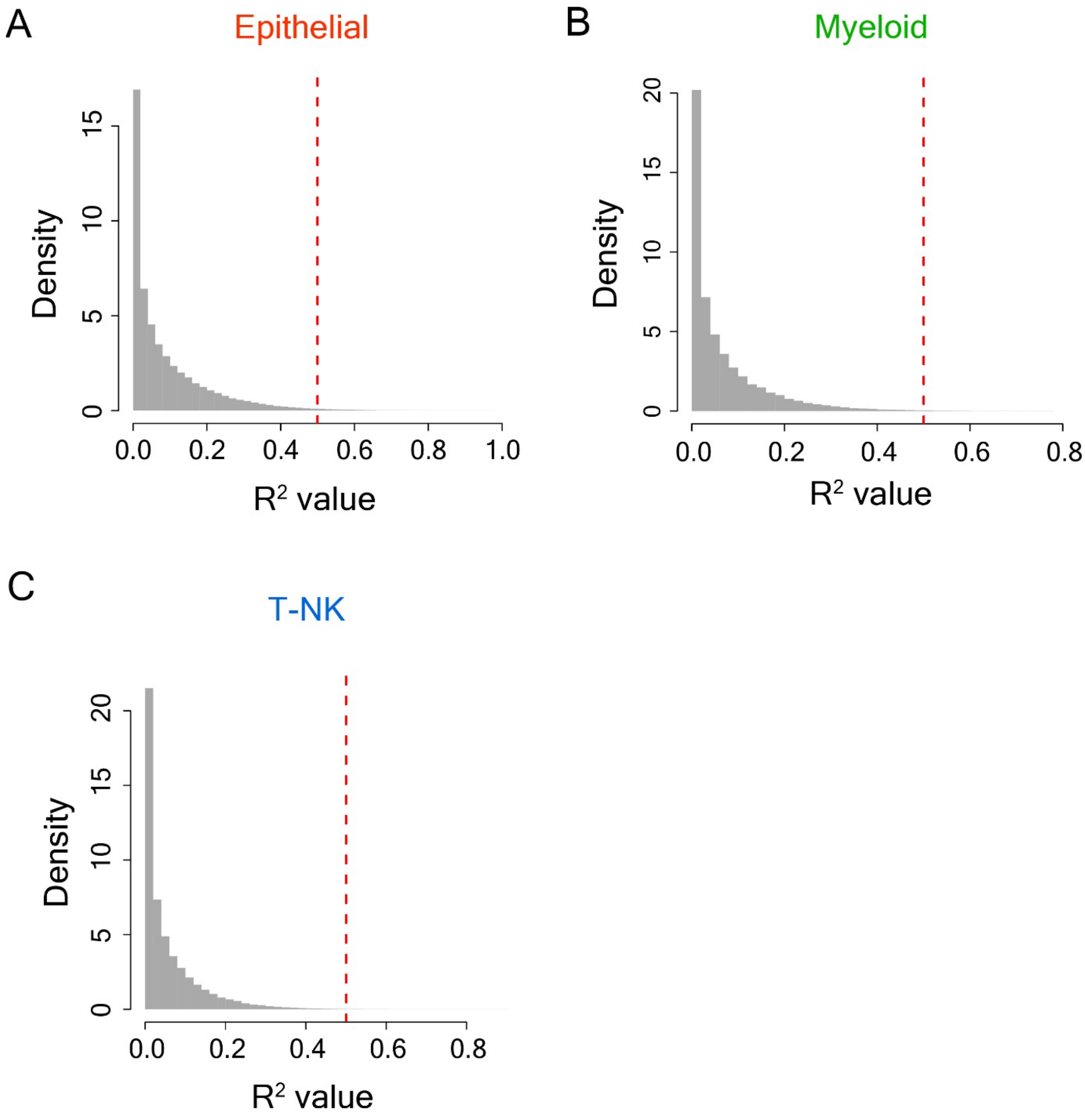


**Figure S4 –** Histograms showing distribution of the goodness-of-fit ($R^2$) values of the linear regression lines for correlation trend analysis for all considered TF-target connections in epithelial cells **(A),** in myeloid cells **(B)** and in T-NK cells **(C)**. The red dotted lines show the $R^2$ threshold, and only the TF-target connections with $R^2 \geq 0.5$ were considered to exhibit a transition in gene regulation with cancer progression.

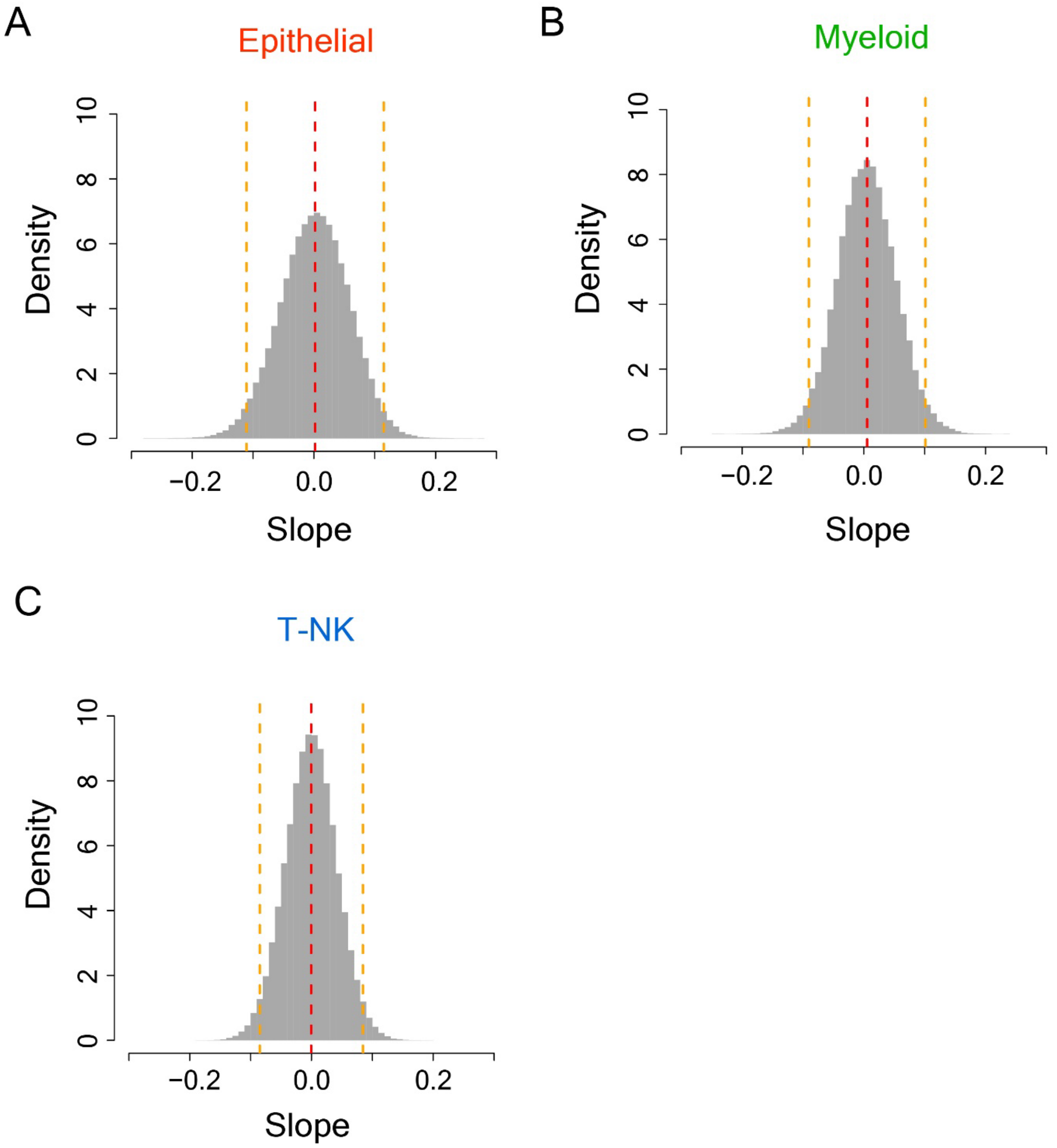


**Figure S5 -** Histograms showing distribution of the slope values of the linear regression lines for correlation trend analysis for all considered TF-target connections in epithelial cells **(A)**, in myeloid cells **(B)** and in T-NK cells **(C)**. The red dotted lines show the mean values, and the orange dotted lines show the mean ± 2×s.d. values.

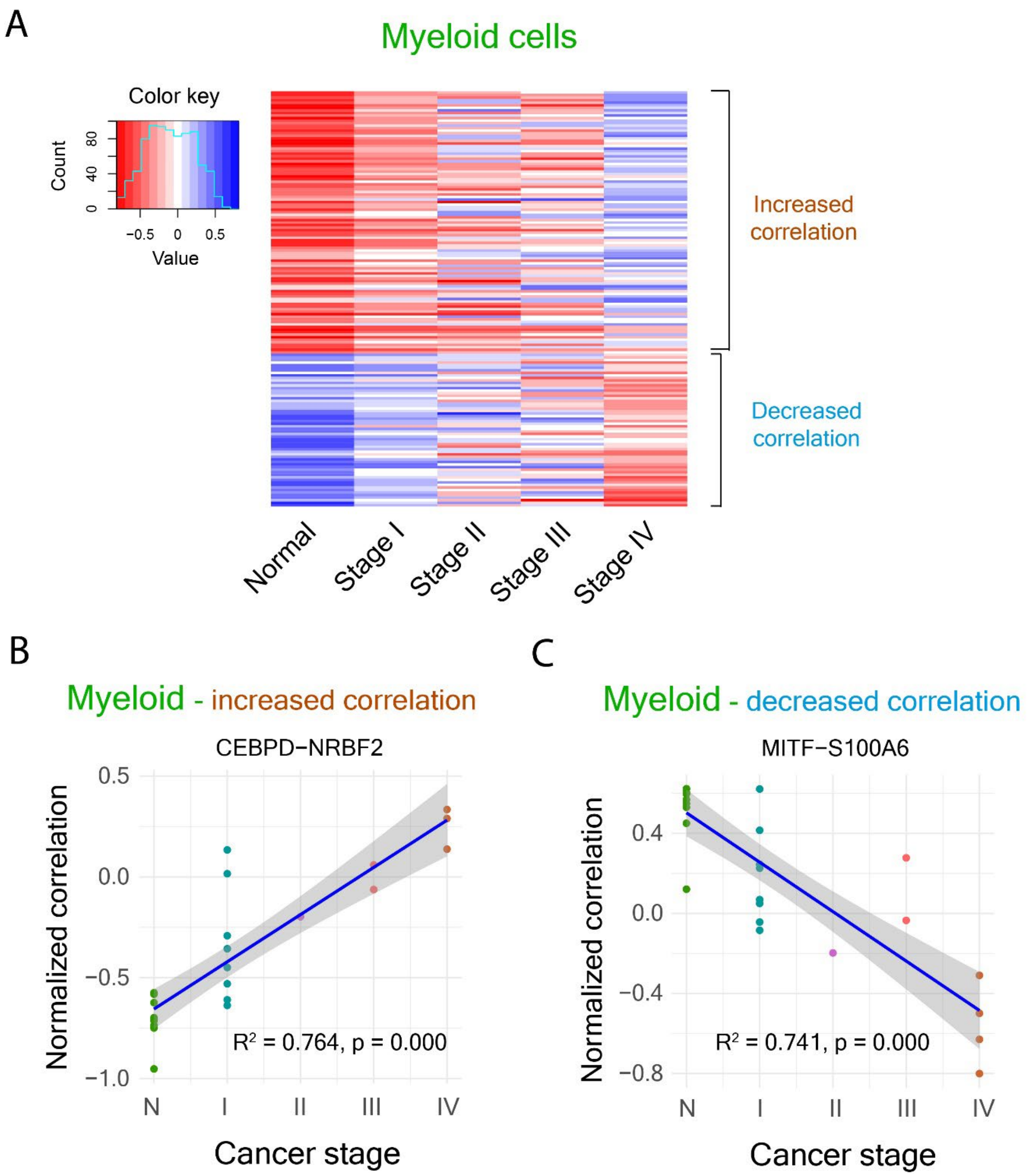


**Figure S6 - TF-target connections undergoing transitions with cancer progression in myeloid cells. (A)** Heatmap showing the changes in mean correlation values with cancer progression for the TF-target connections that showed linearly increasing (red to blue) or decreasing (blue to red) correlation trends in myeloid cells. These TF-target connections were identified by linear regression with $R^2$ greater than or equal to 0.5 and the slope value was beyond mean ± 2× s.d. values, where mean and s.d. values of the slope were calculated from all TF-target connections in myeloid cells. **(B-C)** Examples of TF-target connections showing a positive correlation trend (increasing correlation) (B) and a negative correlation trend (decreasing correlation) (C) with cancer progression in myeloid cells.

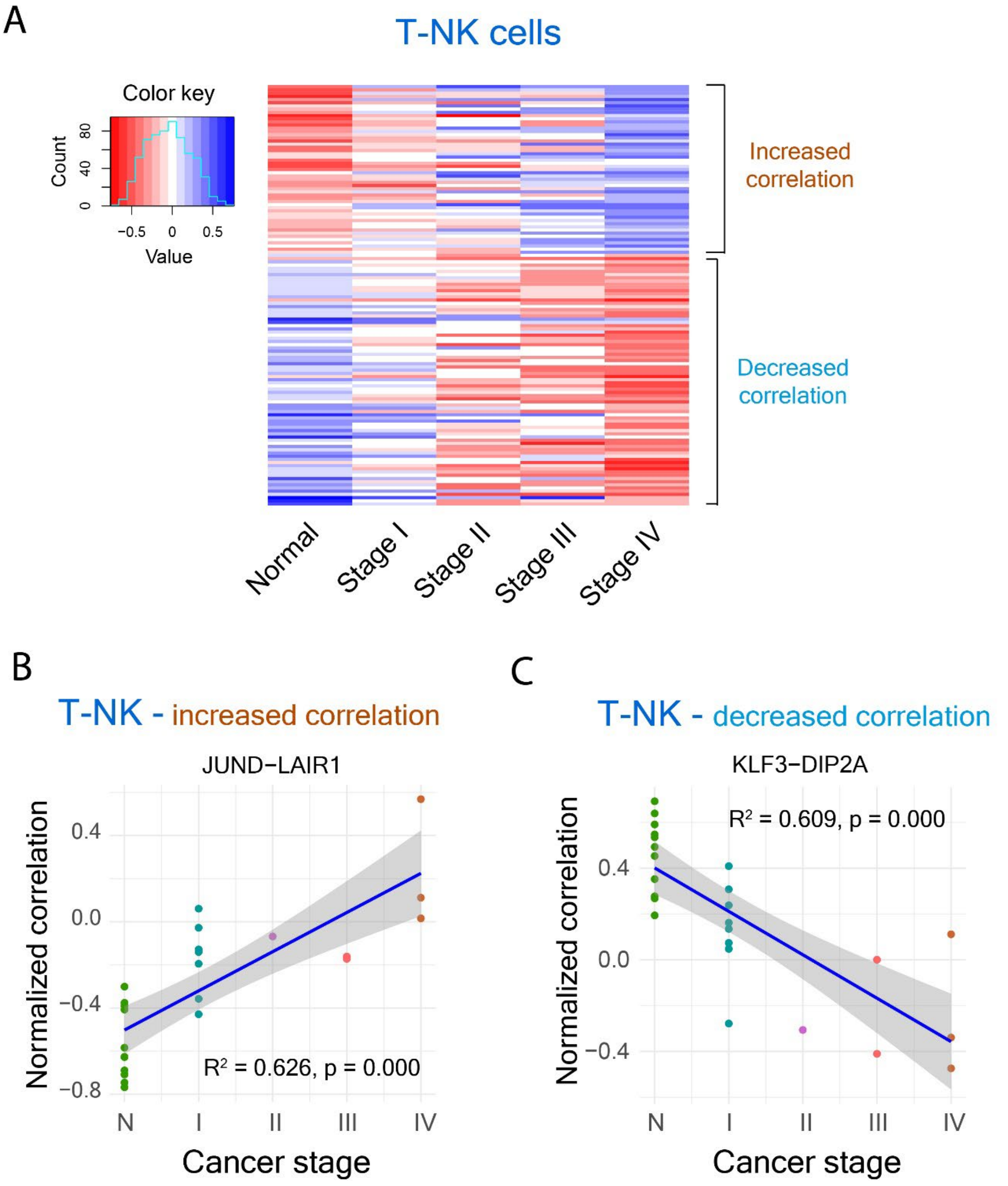


**Figure S7 - TF-target connections undergoing transitions with cancer progression in T-NK cells. (A)** Heatmap showing the changes in mean correlation values with cancer progression for the TF-target connections that showed linearly increasing (red to blue) or decreasing (blue to red) correlation trends in T-NK cells. These TF-target connections were identified by linear regression with $R^2$ greater than or equal to 0.5 and the slope value was beyond mean ± 2× s.d. values, where mean and s.d. values of the slope were calculated from all TF-target connections in T-NK cells. **(B-C)** Examples of TF-target connections showing a positive correlation trend (increasing correlation) (B) and a negative correlation trend (decreasing correlation) (C) with cancer progression in T-NK cells.

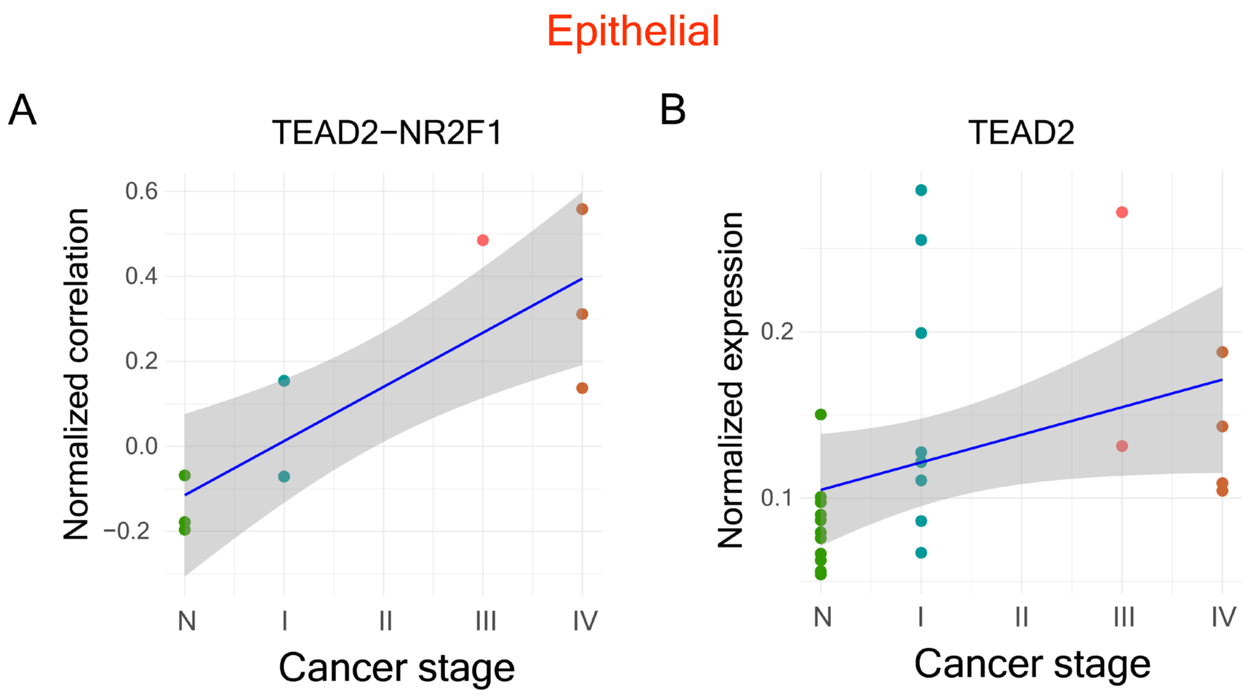


**Figure S8 – (A)** Increased expression correlation between the TF TEAD2 and its target NR2F1 in epithelial cells with cancer progression. **(B)** Increased normalized mean expression values of the TF TEAD2 in epithelial cells with cancer progression.

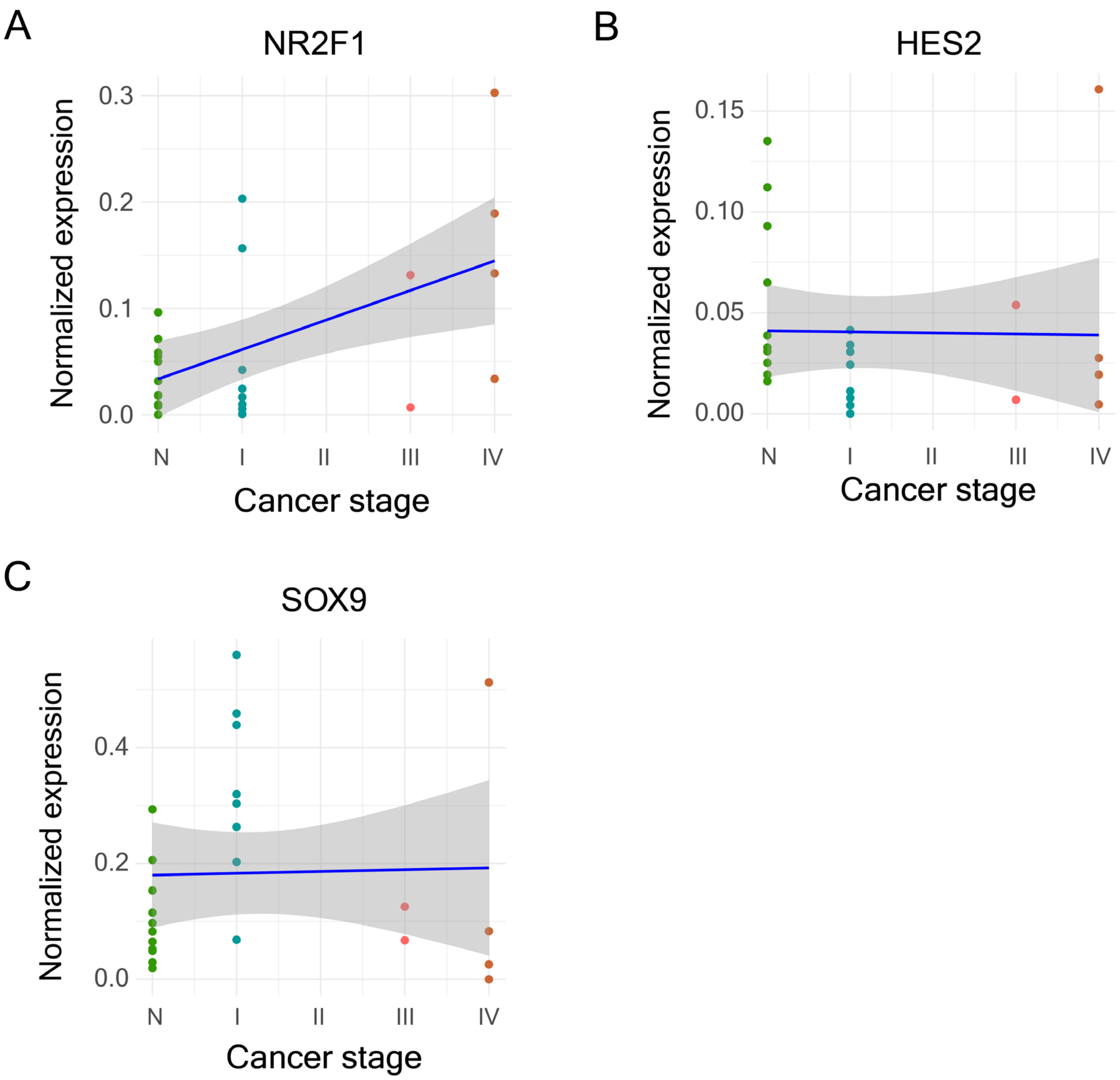


**Figure S9** – Normalized mean expression values of the TFs NR2F1 **(A)**, HES2 **(B)**, and SOX9 **(C)** in epithelial cells of normal samples and cancer samples at different stages.

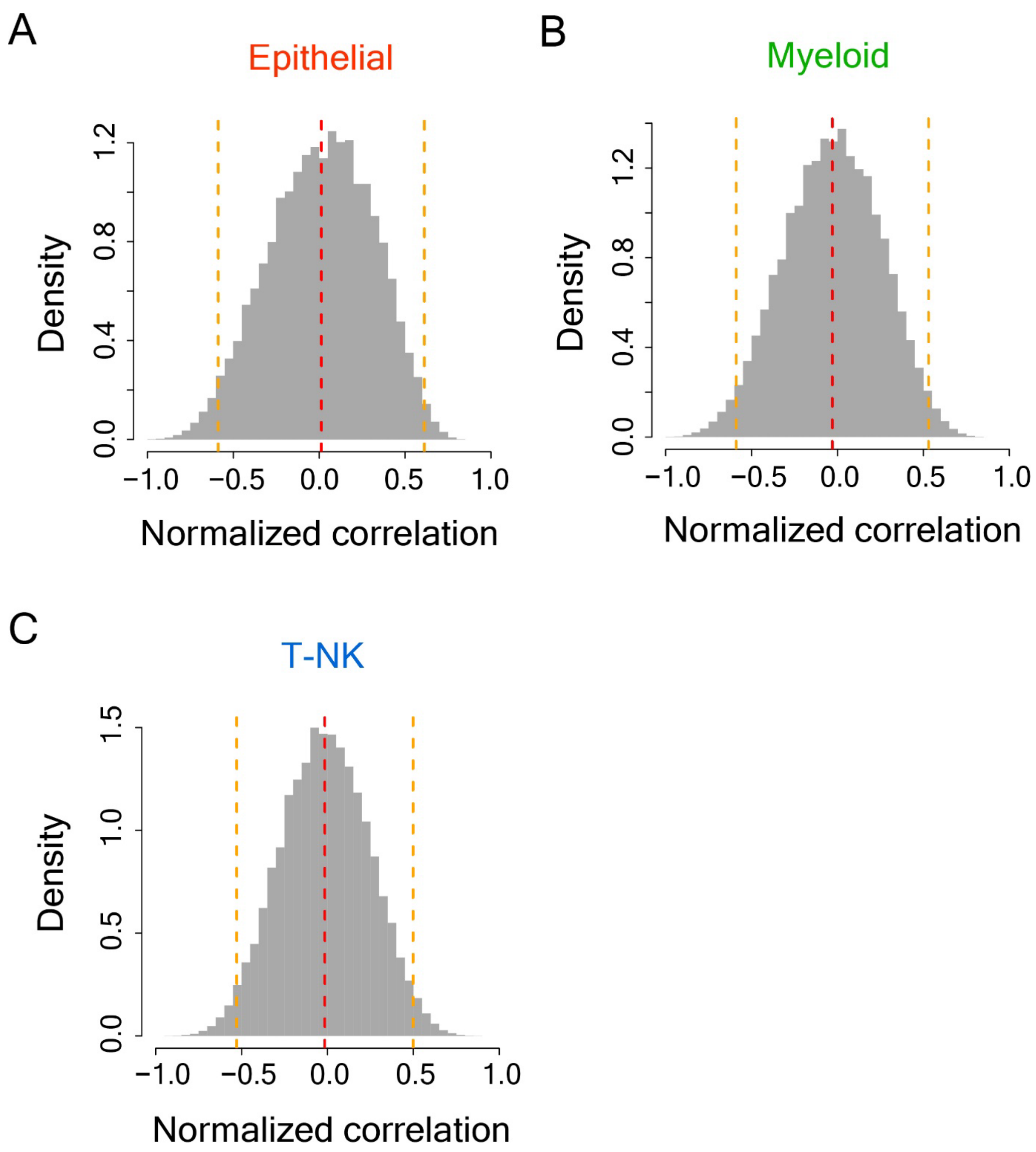


**Figure S10 -** Histograms showing distributions of normalized correlation values of all TF-target connections in epithelial cells **(A)**, myeloid cells **(B)**, and T-NK cells **(C)**. The red dotted lines show the mean correlation values of the distributions and the orange dotted lines show mean ± 2×s.d. values.

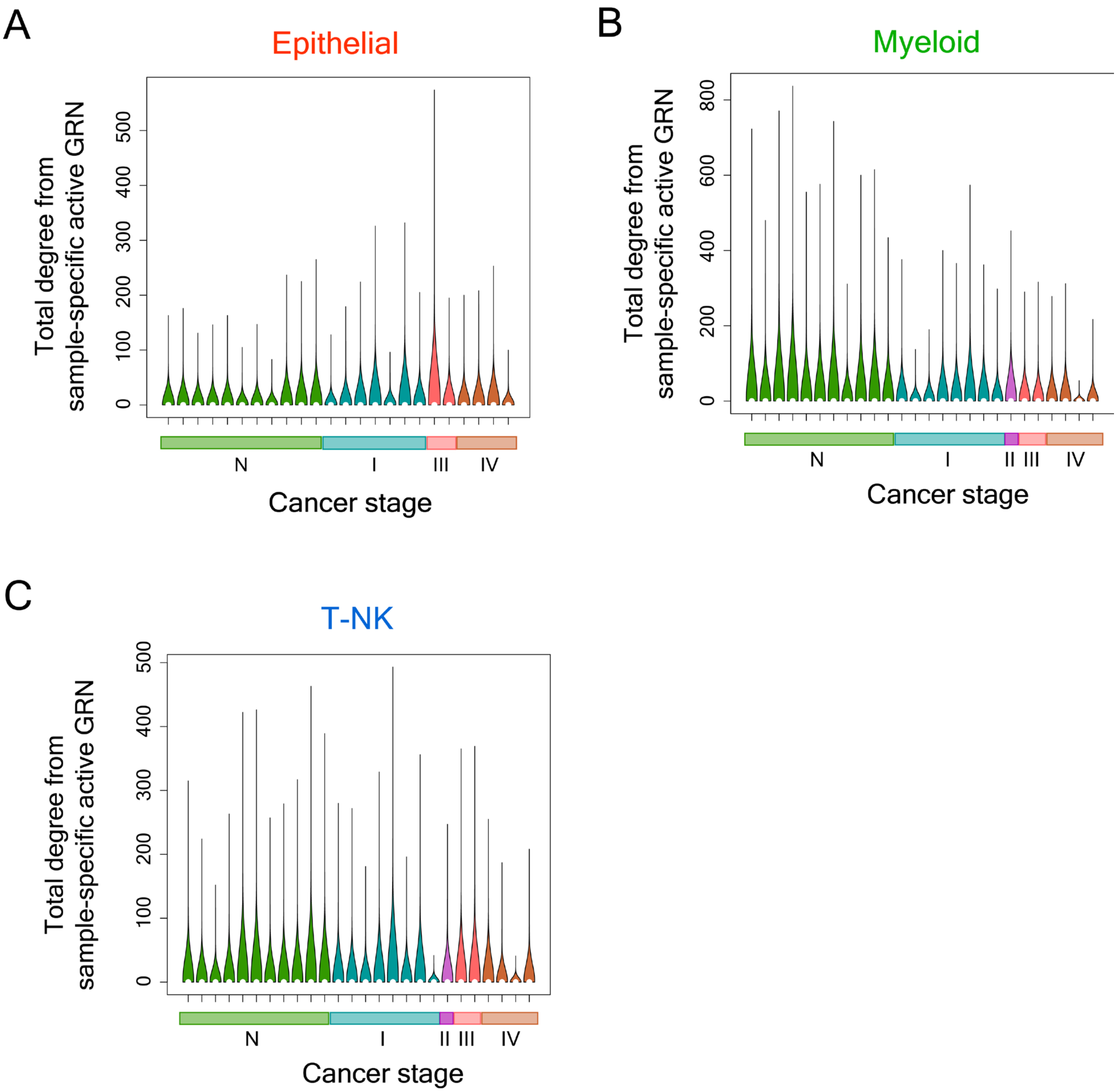


**Figure S11 –** Violin plots showing the distributions of total degree values of active GRNs in normal as well as cancer samples of different stages in epithelial cells **(A)**, myeloid cells **(B)**, and T-NK cells **(C)**.

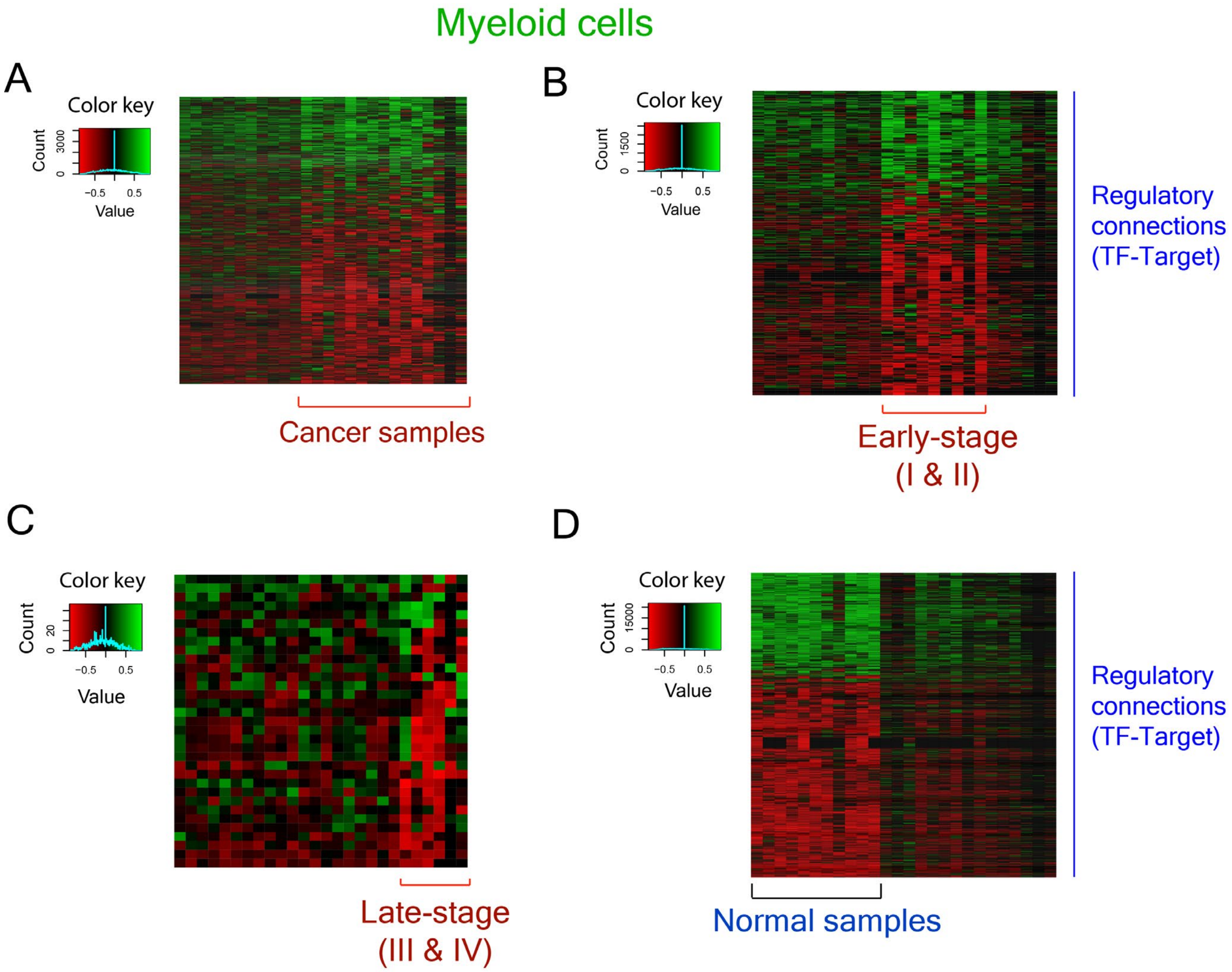


**Figure S12 –** Heatmaps showing the TF-target connections in myeloid cells that are active in only cancer samples (encompassing both early- and late-stage samples) **(A)**, only early-stage cancer samples **(B)**, only late-stage cancer samples **(C)** and only normal samples **(D)**. Green color indicates positive TF-target correlation and red color indicates negative TF-target correlation.

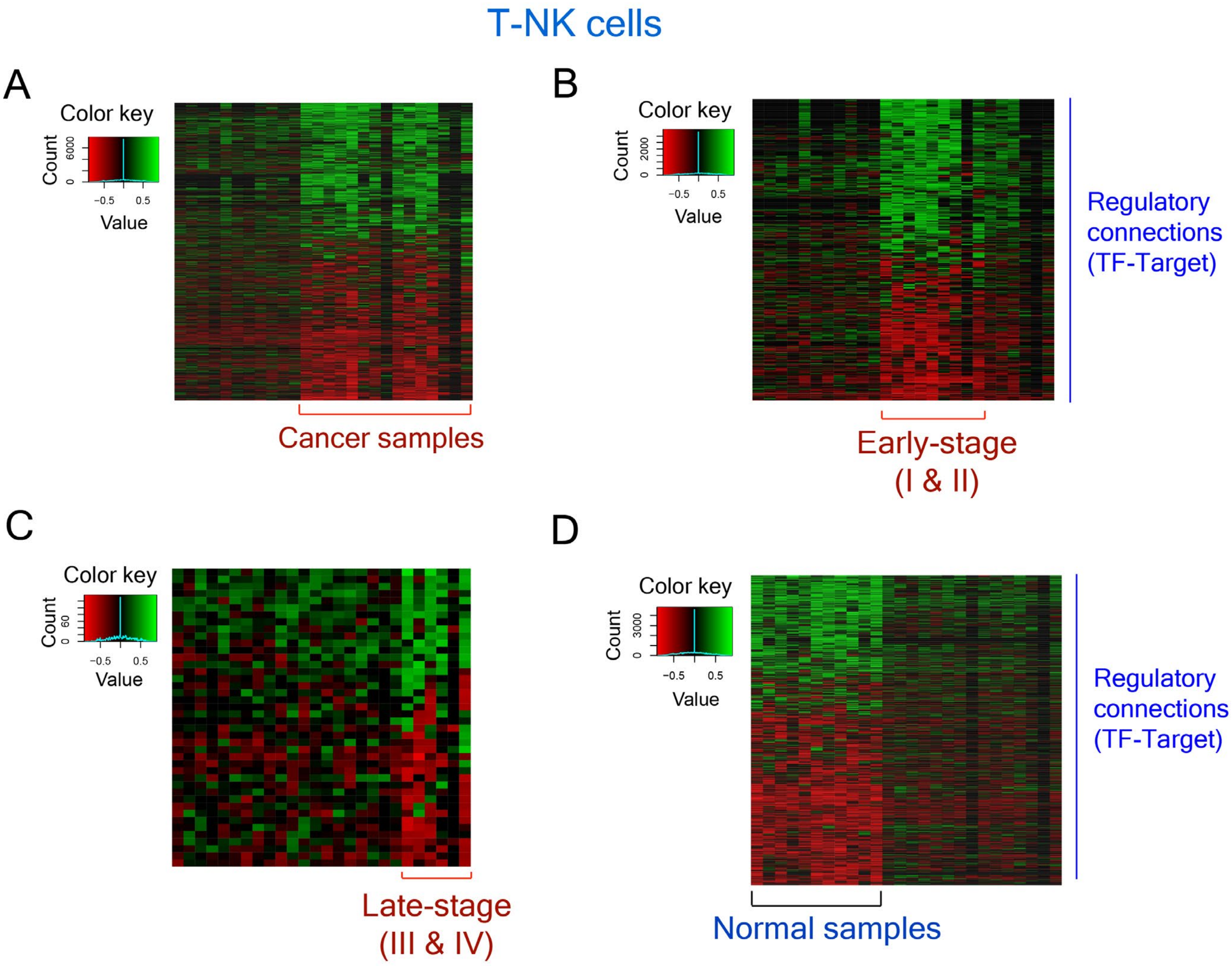


**Figure S13 -** Heatmaps showing the TF-target connections in T-NK cells that are active in only cancer samples (encompassing both early- and late-stage samples) **(A)**, only early-stage cancer samples **(B)**, only late-stage cancer samples **(C)** and only normal samples **(D)**. Green color indicates positive TF-target correlation and red color indicates negative TF-target correlation.

### Myeloid cells

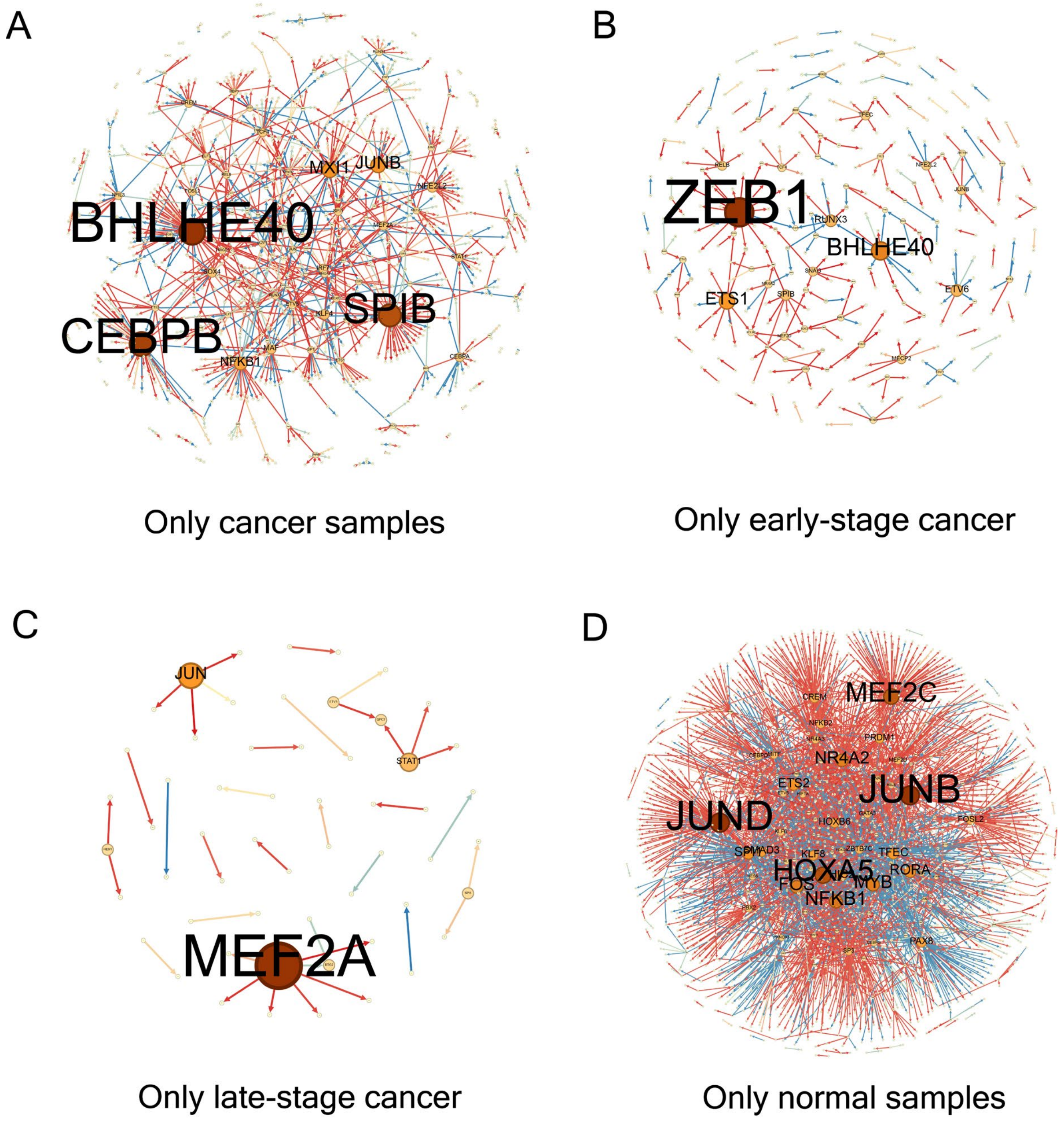


**Figure S14 -** Graphs showing the degree of genes (both TFs and targets) in myeloid cells that appear in the connections active in only cancer samples **(A)**, only early-stage cancer samples **(B)**, only late-stage cancer samples **(C)**, and only normal samples **(D)**. Red edges show negative TF-target correlation and blue edges show positive TF-target correlation.

#### T-NK cells

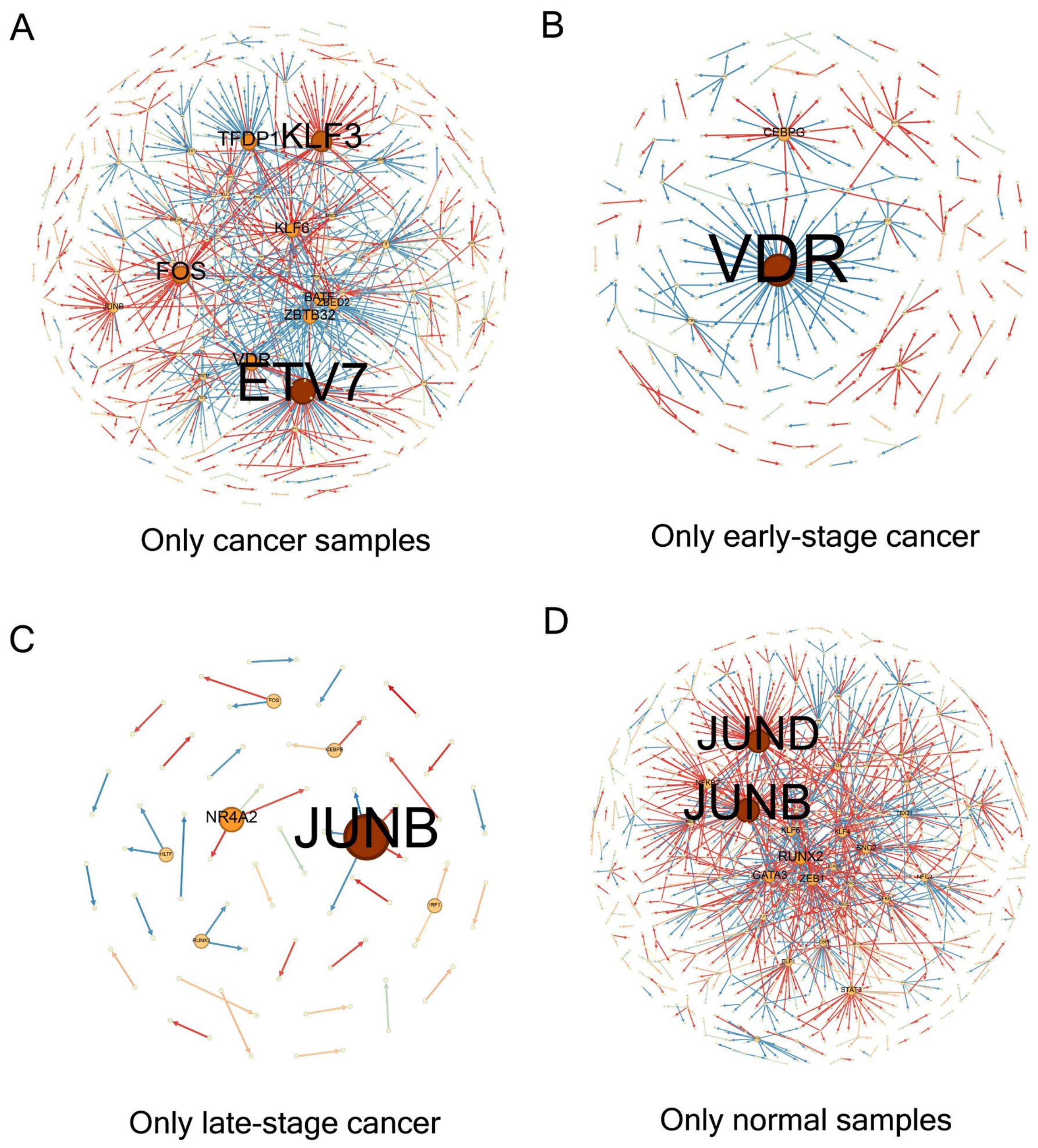


**Figure S15 -** Graphs showing the degree of genes (both TFs and targets) in T-NK cells that appear in the connections active in only cancer samples **(A)**, only early-stage cancer samples **(B)**, only late-stage cancer samples **(C)**, and only normal samples **(D)**. Red edges show negative TF-target correlation and blue edges show positive TF-target correlation.

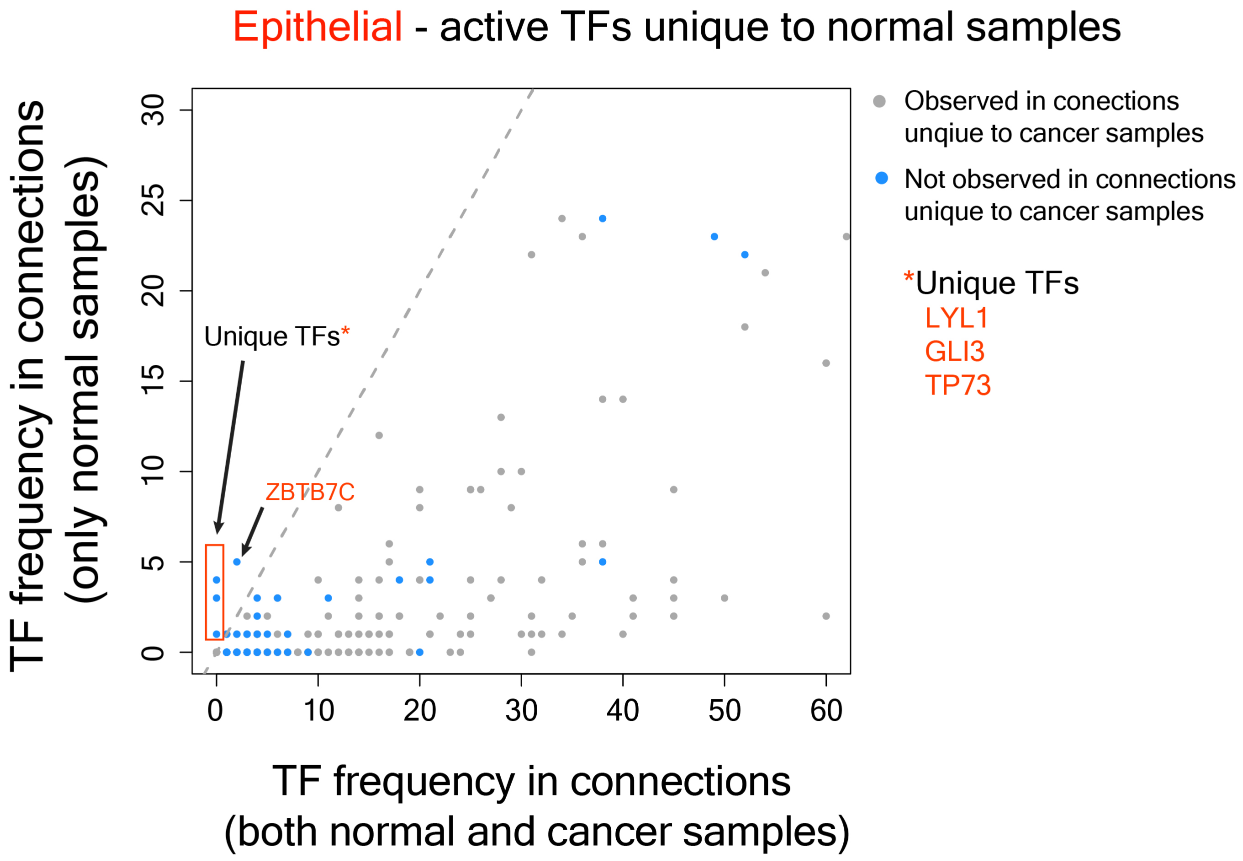


**Figure S16 -** Plot showing the frequency of TFs in connections that occur only in normal samples and the frequency of the same TFs in connections that occur in both normal and cancer samples in epithelial cells.

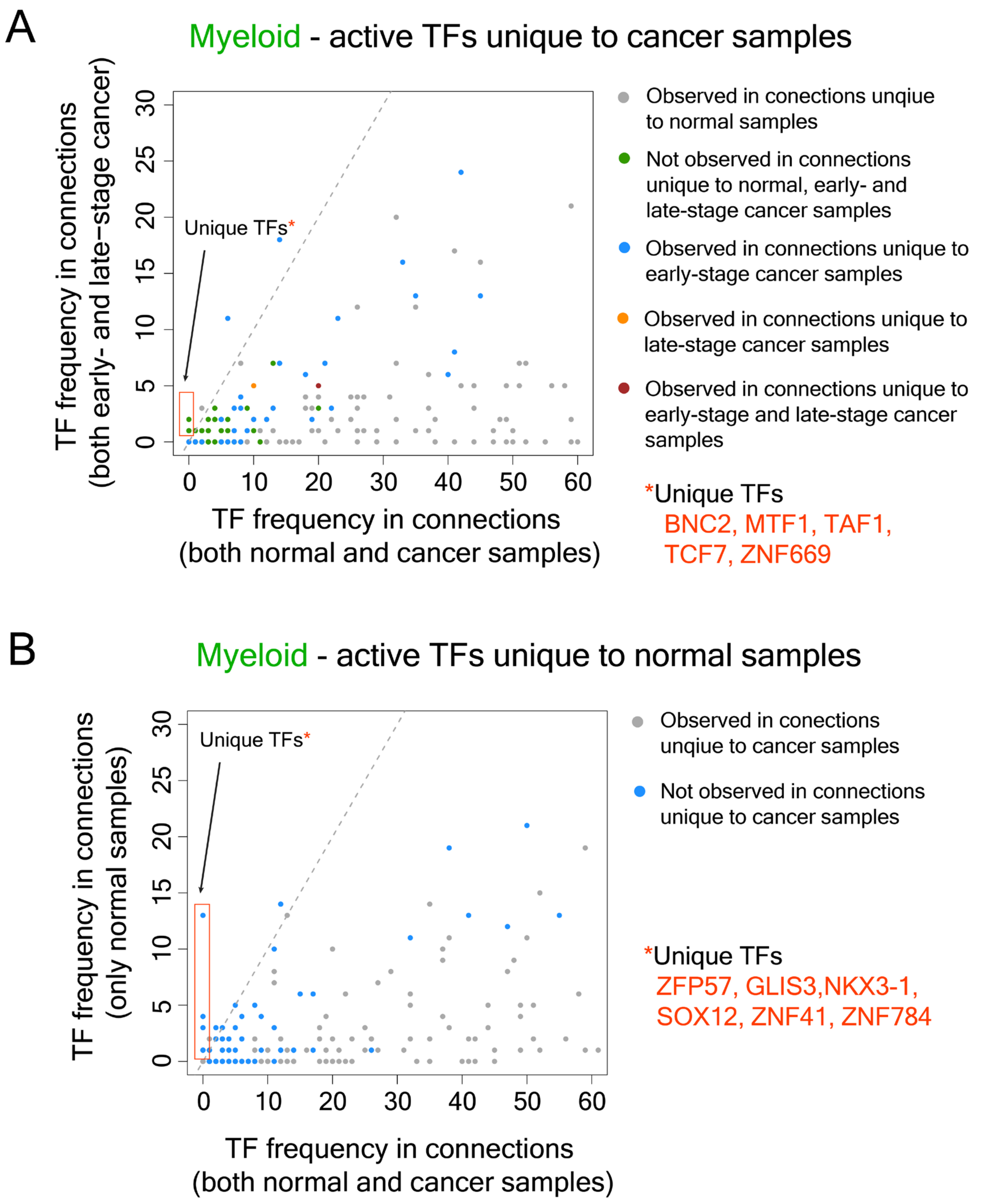


**Figure S17 – (A)** Plot showing the frequency of TFs in connections that occur only in cancer samples and the frequency of the same TFs in connections that occur in both normal and cancer samples in epithelial cells. **(B)** Plot showing the frequency of TFs in connections that occur only in normal samples and the frequency of the same TFs in connections that occur in both normal and cancer samples in epithelial cells.

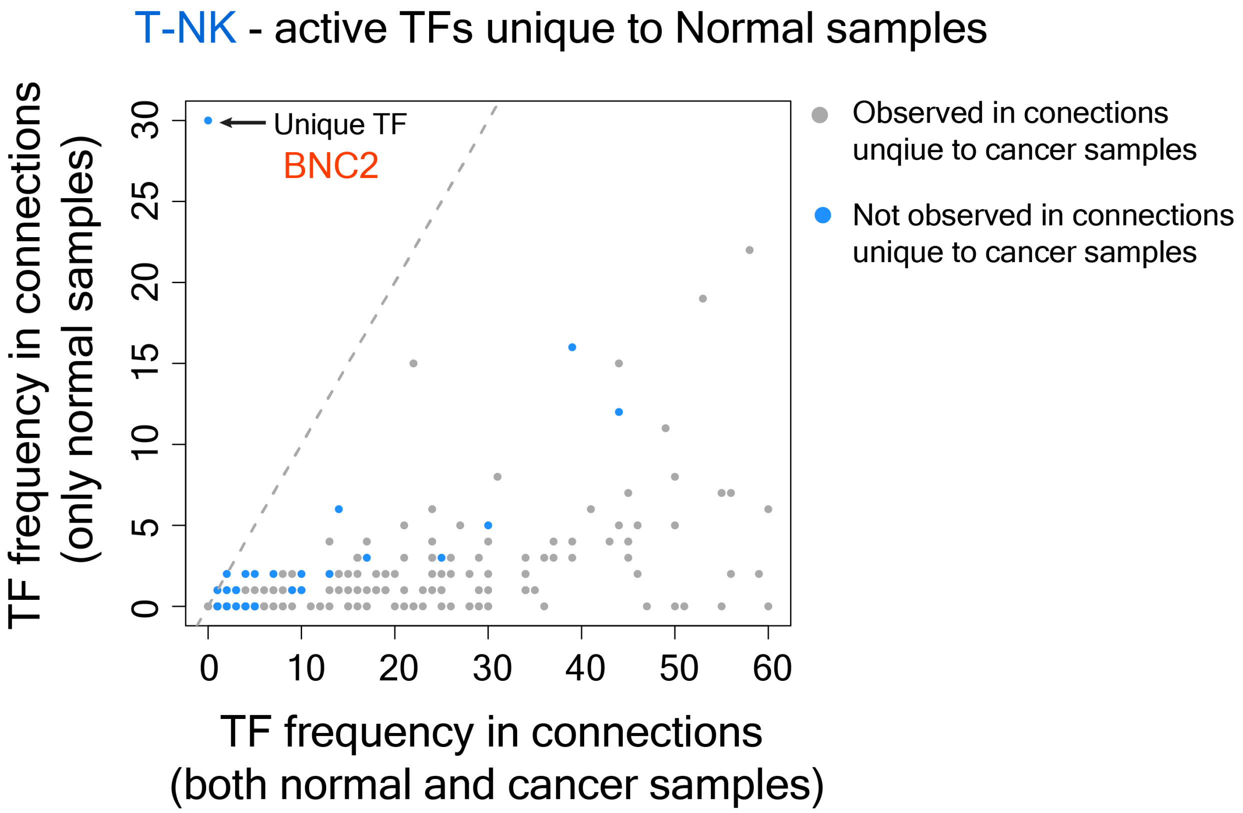


**Figure S18 -** Plot showing the frequency of TFs in connections that occur only in normal samples and the frequency of the same TFs in connections that occur in both normal and cancer samples in T-NK cells.

A

Coordinated Myl and T-NK TF regulation in early-stage cancer

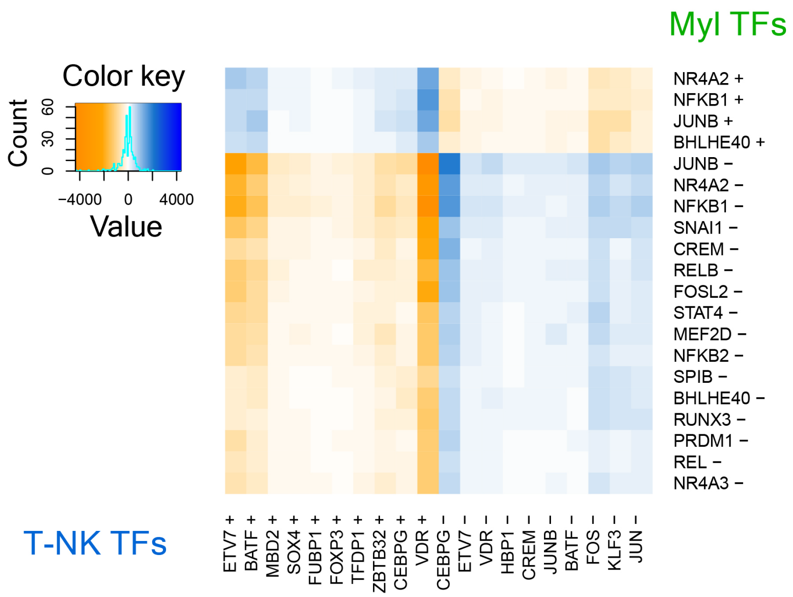


B

Coordinated Myl and T-NK TF regulation in late-stage cancer

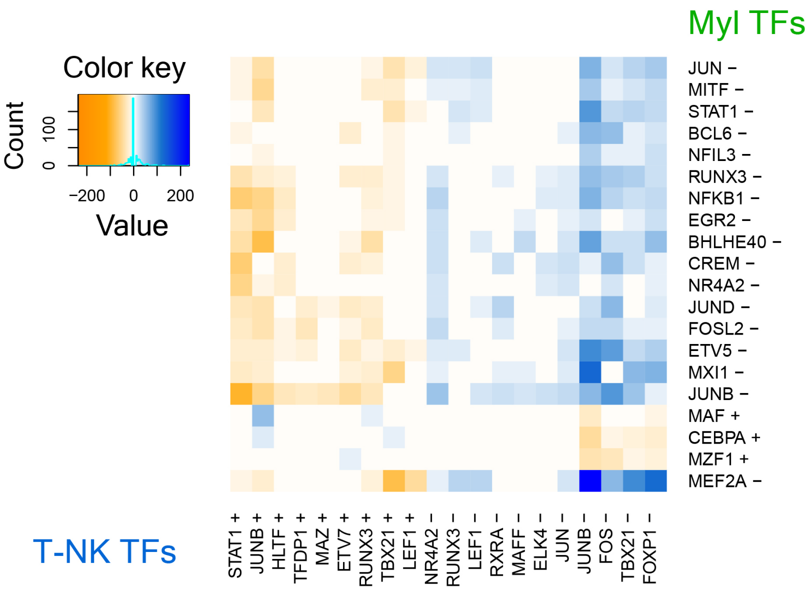


**Figure S19 -** Top 20 TFs in myeloid cells and T-NK cells with coordinated regulatory activities in early-stage **(A)** and late-stage **(B)** cancer samples.

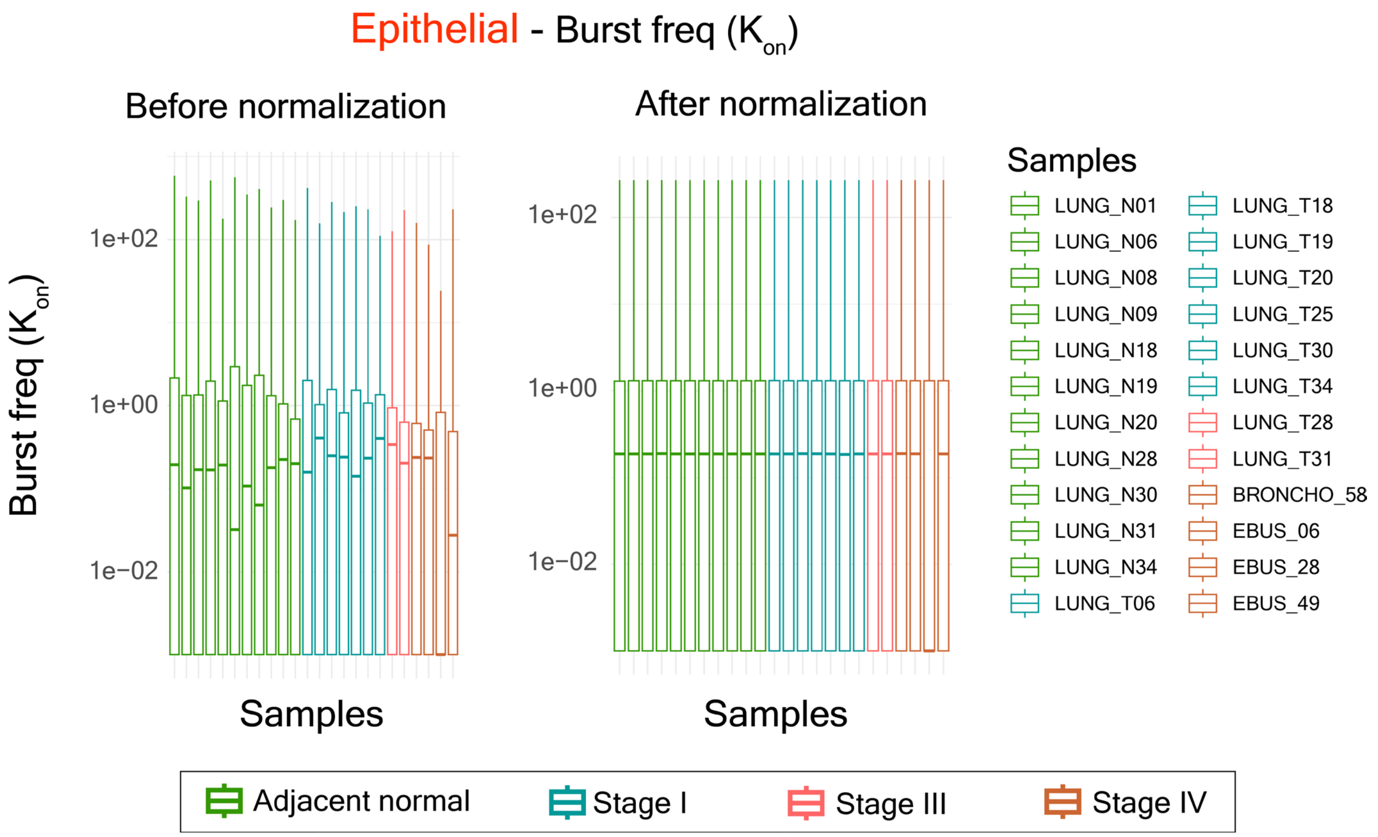


**Figure S20 –** Boxplots showing distributions of burst frequency ($K_{on}$) values in epithelial cells of normal samples and cancer samples before normalization (left) and after normalization (right).

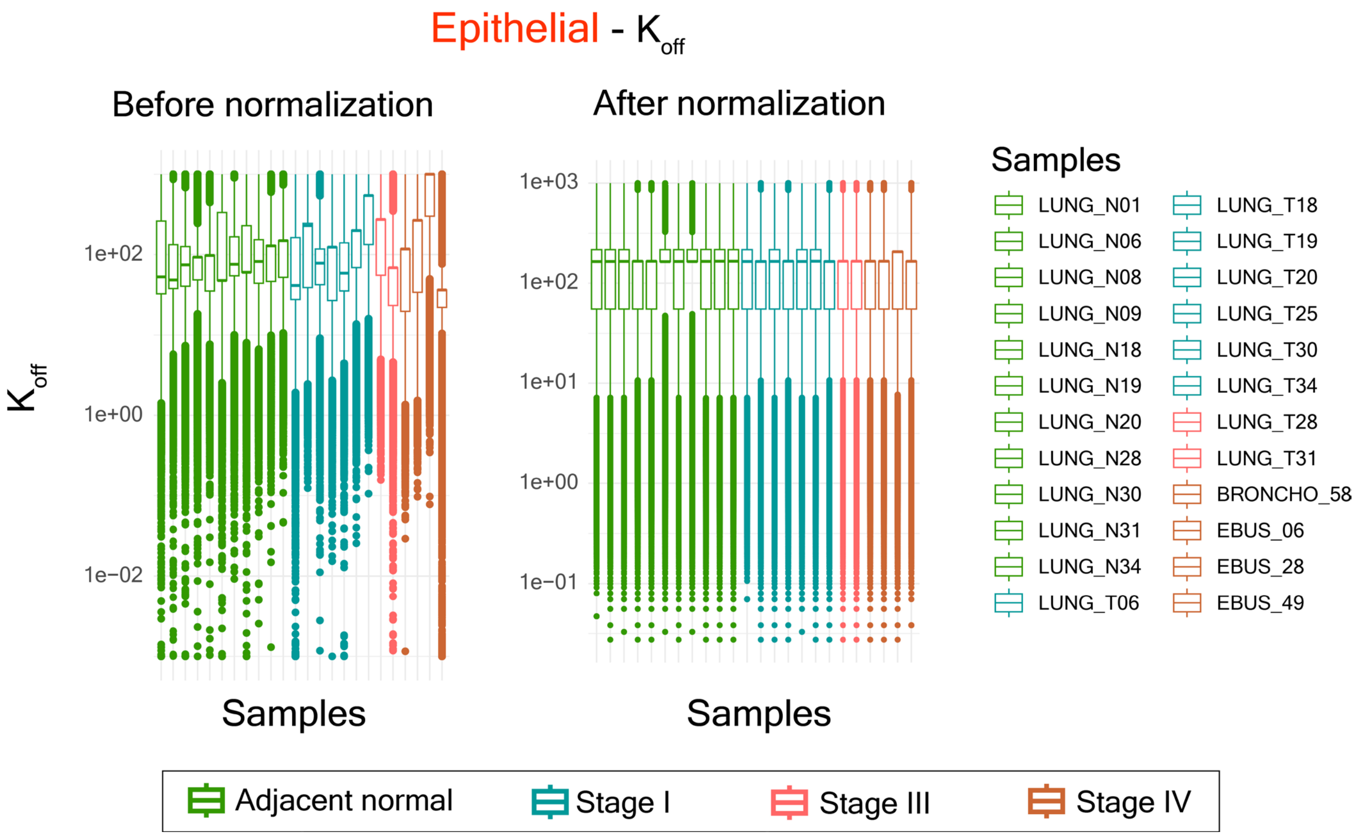


**Figure S21 -** Boxplots showing distributions of $K_{off}$ values in epithelial cells of normal samples and cancer samples before normalization (left) and after normalization (right).

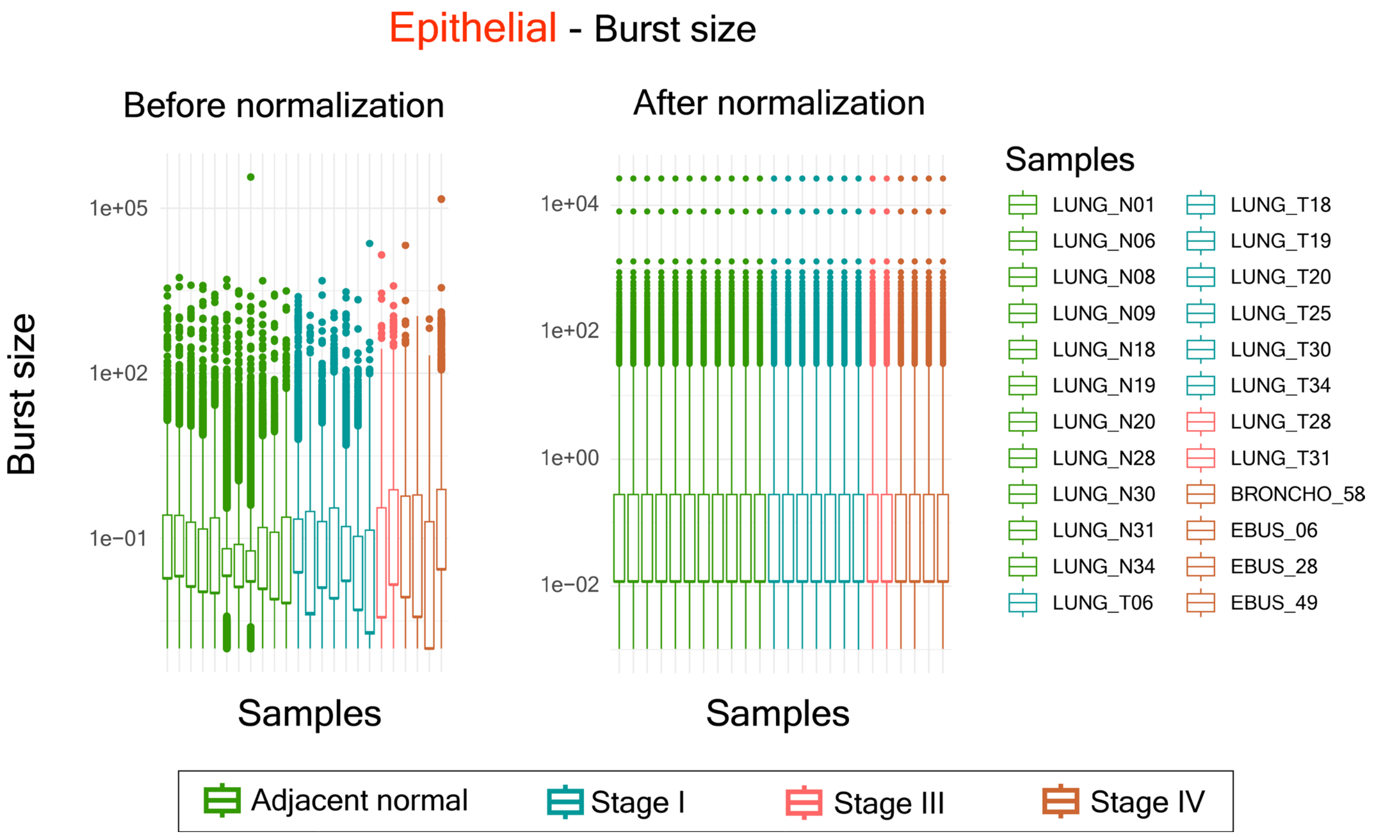


**Figure S22 -** Boxplots showing distributions of burst size values in epithelial cells of normal samples and cancer samples before normalization (left) and after normalization (right).

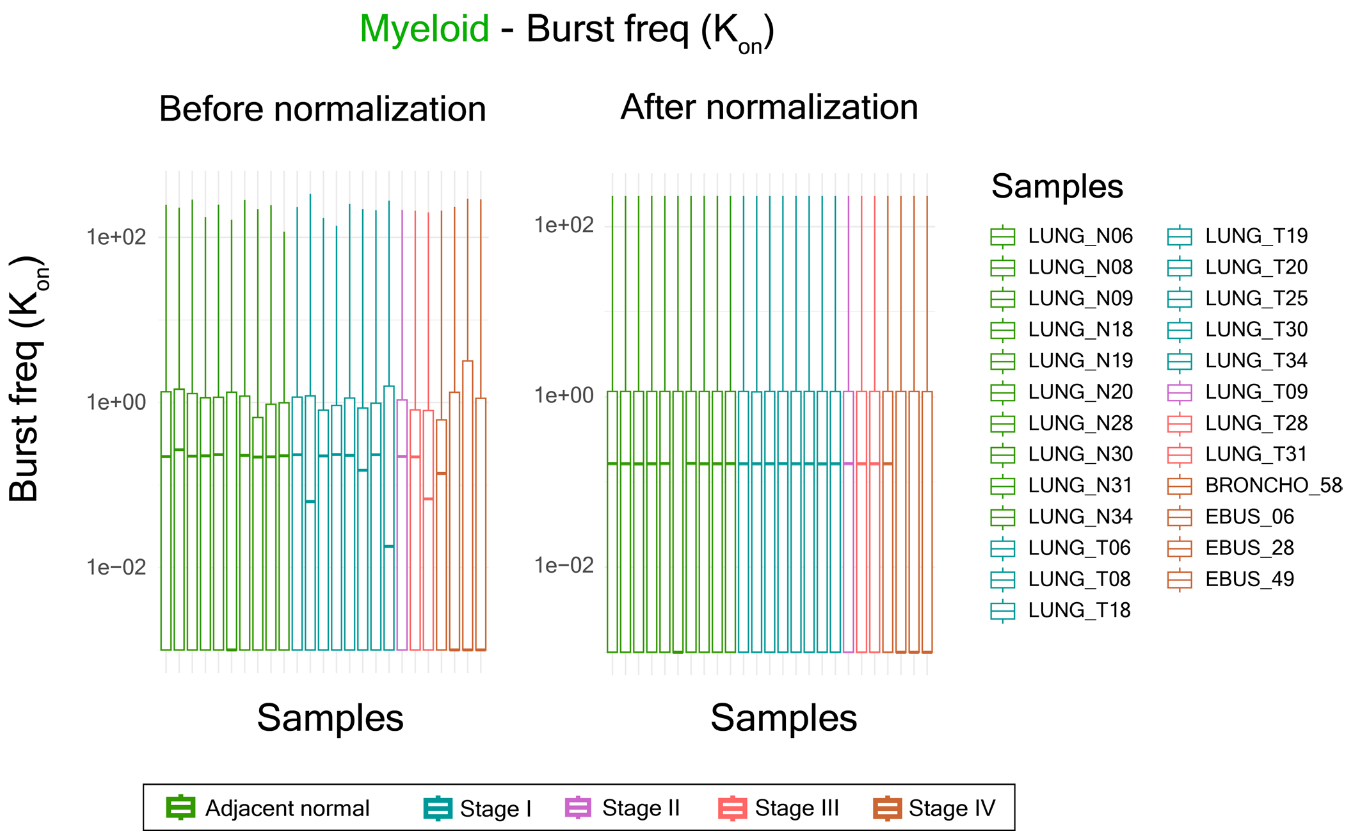


**Figure S23 -** Boxplots showing distributions of burst frequency ($K_{on}$) values in myeloid cells of normal samples and cancer samples before normalization (left) and after normalization (right).

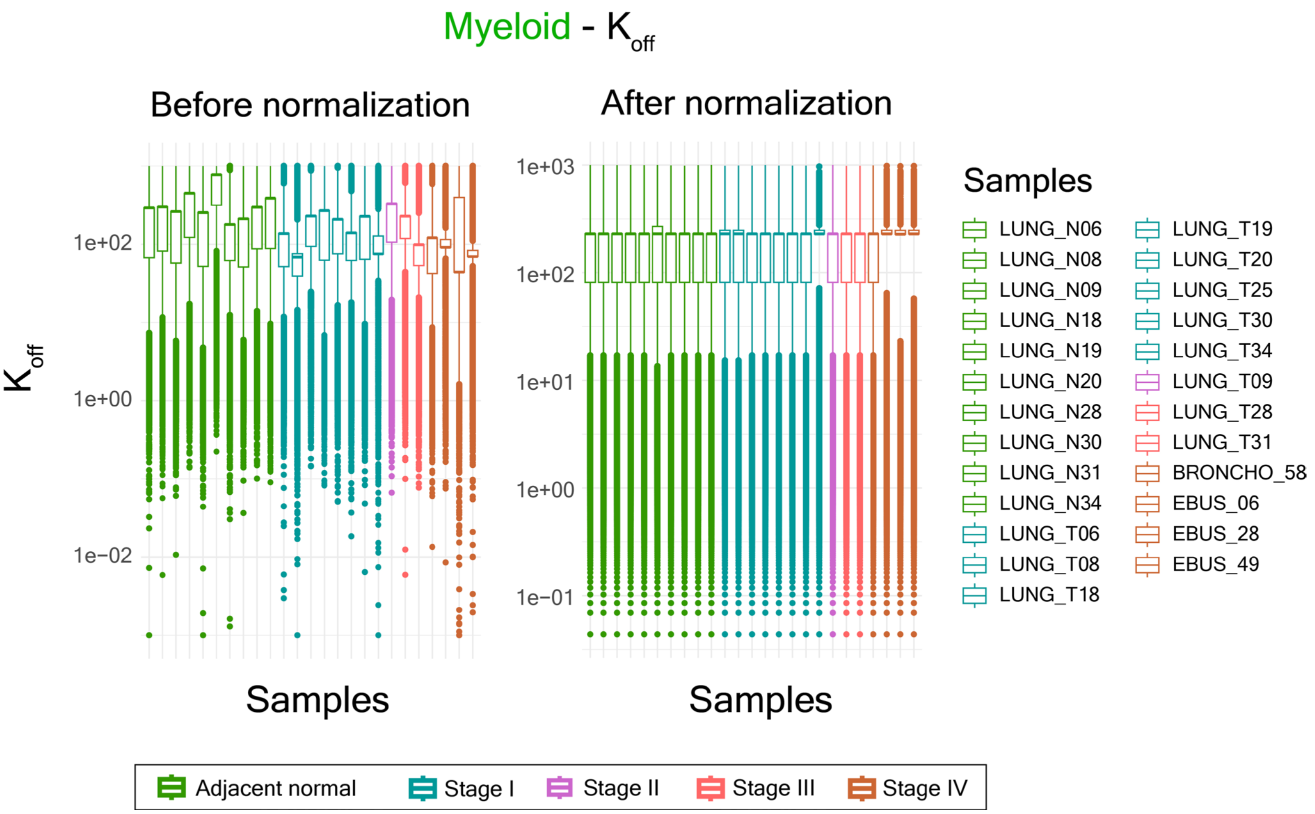


**Figure S24 -** Boxplots showing distributions of $K_{off}$ values in myeloid cells of normal samples and cancer samples before normalization (left) and after normalization (right).

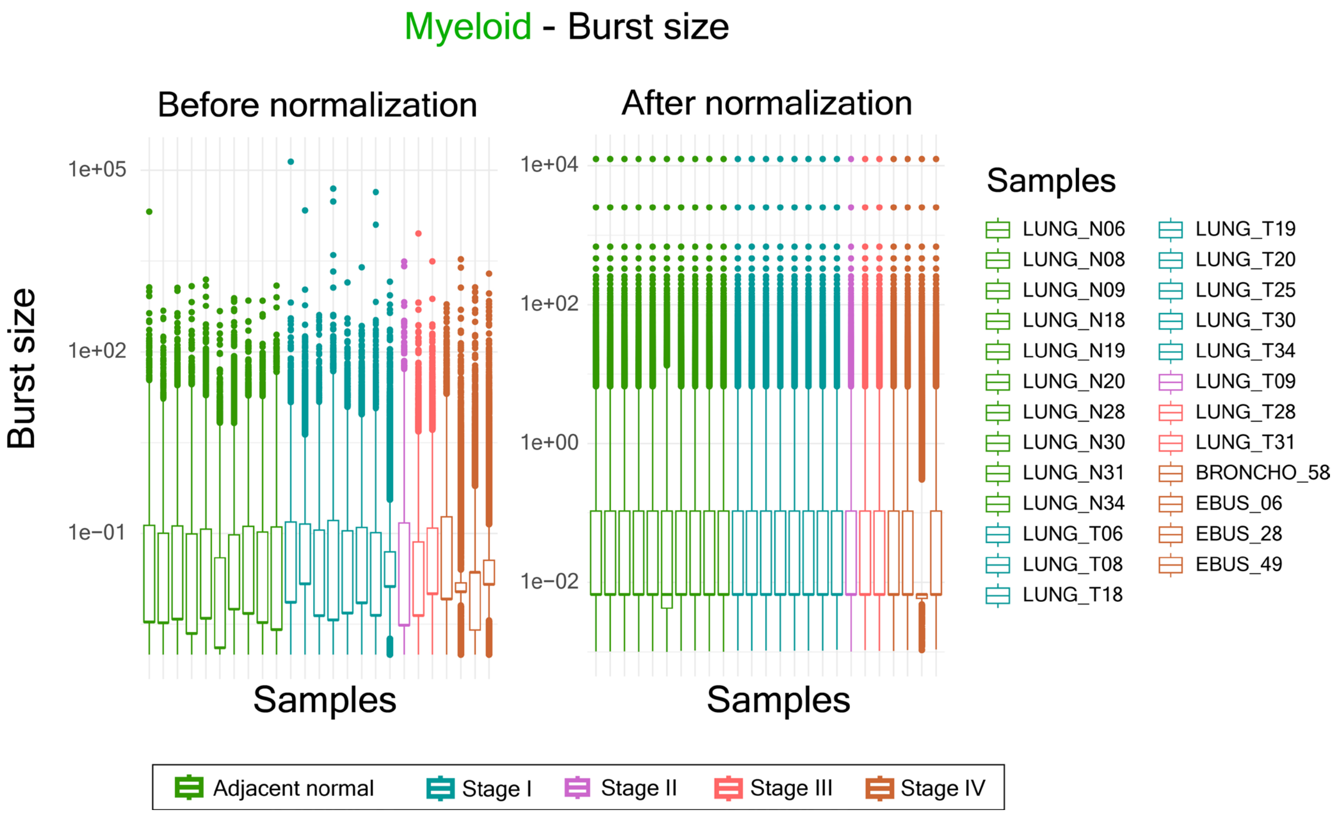


**Figure S25 -** Boxplots showing distributions of burst size values in myeloid cells of normal samples and cancer samples before normalization (left) and after normalization (right).

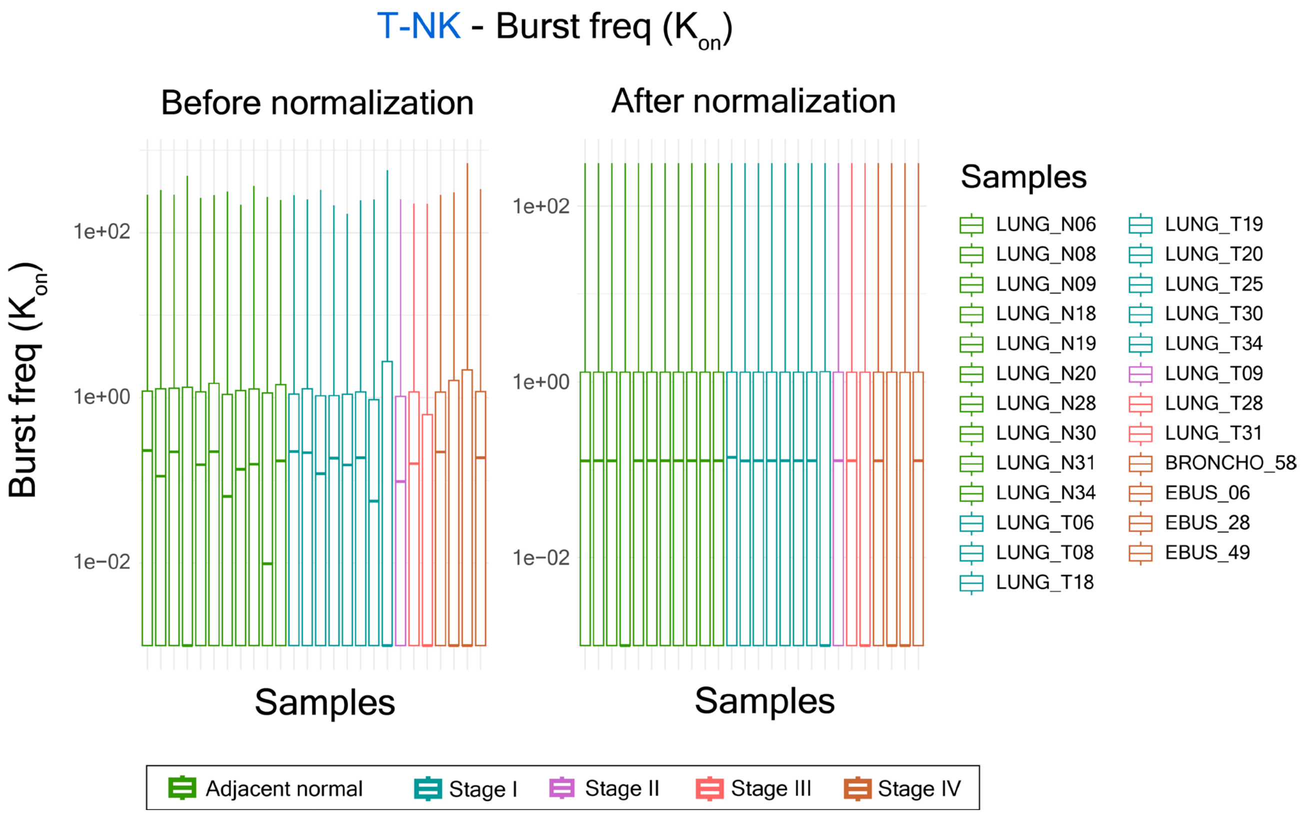


**Figure S26 -** Boxplots showing distributions of burst frequency ($K_{on}$) values in T-NK cells of normal samples and cancer samples before normalization (left) and after normalization (right).

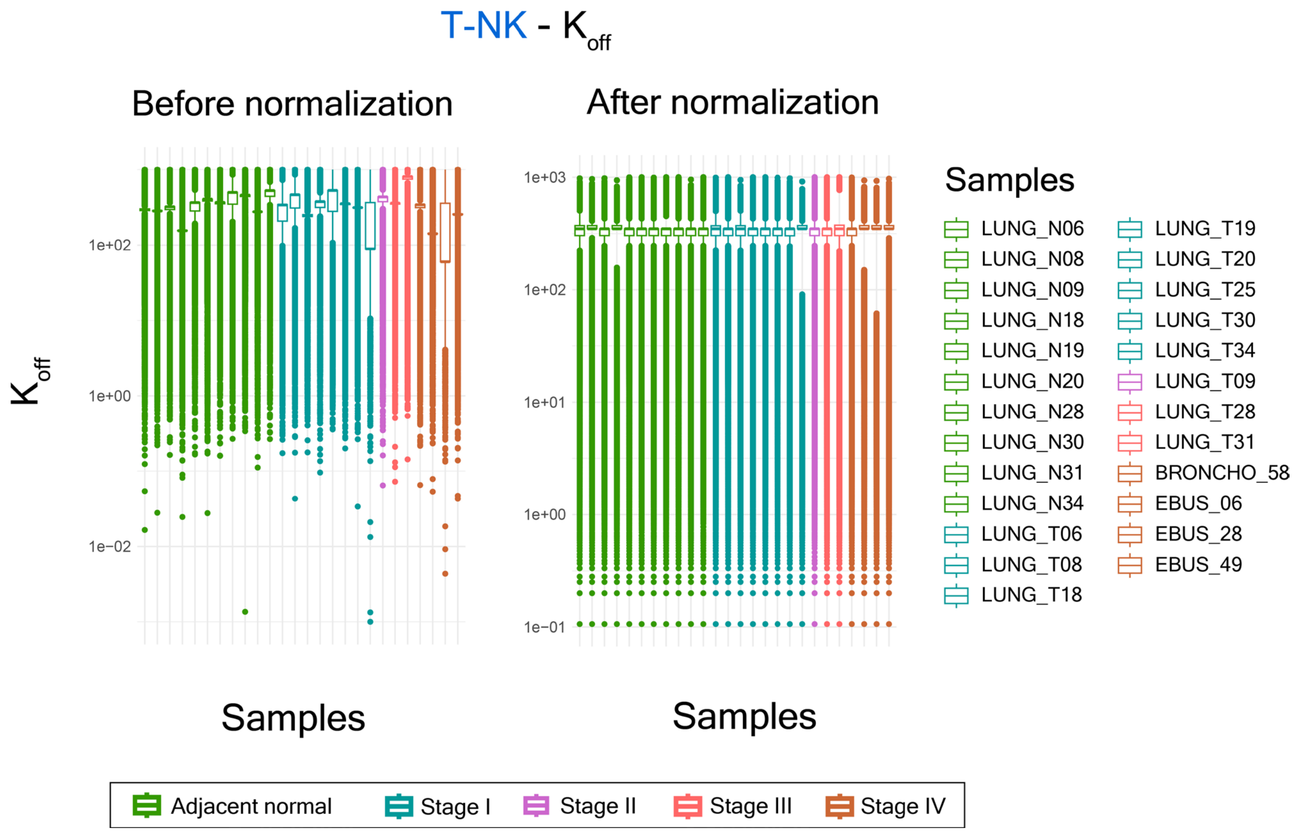


**Figure S27 -** Boxplots showing distributions of $K_{off}$ values in T-NK cells of normal samples and cancer samples before normalization (left) and after normalization (right).

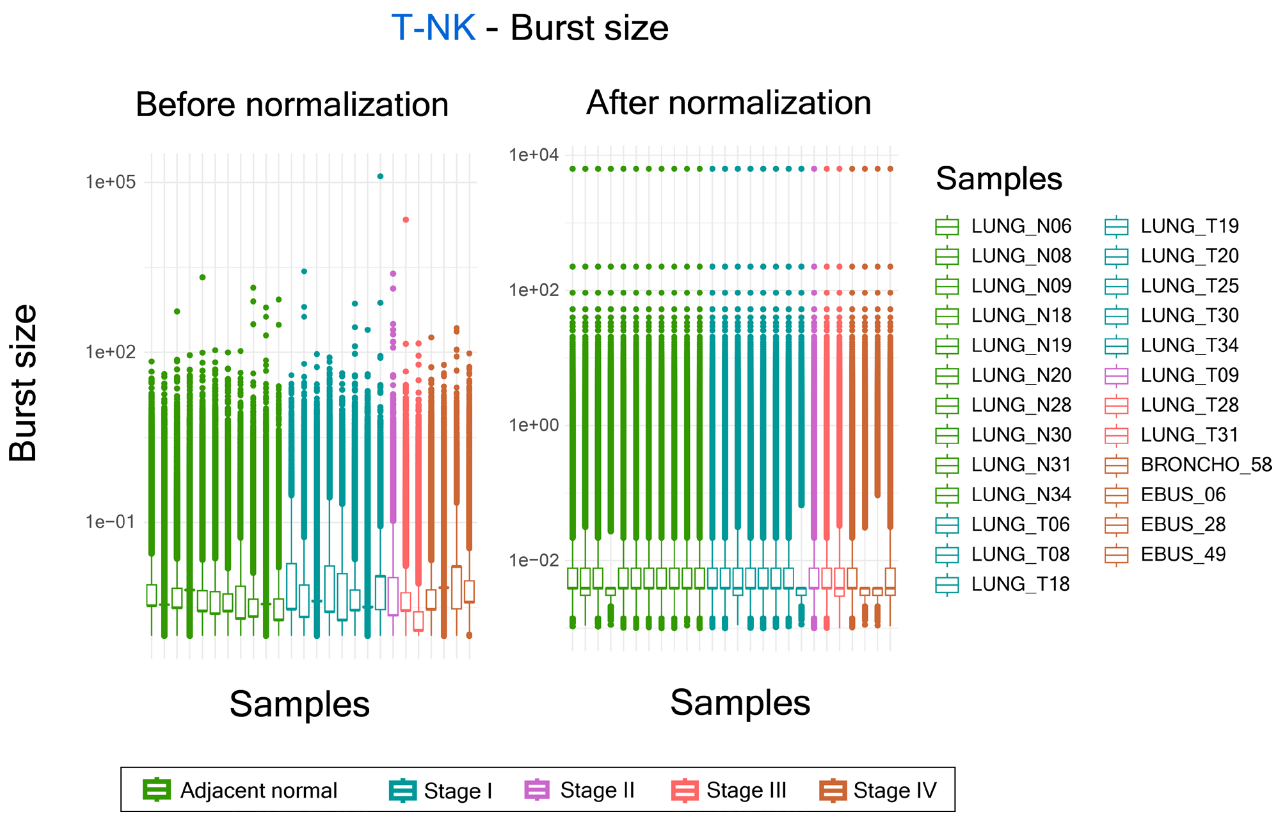


**Figure S28 -** Boxplots showing distributions of burst size values in T-NK cells of normal samples and cancer samples before normalization (left) and after normalization (right).

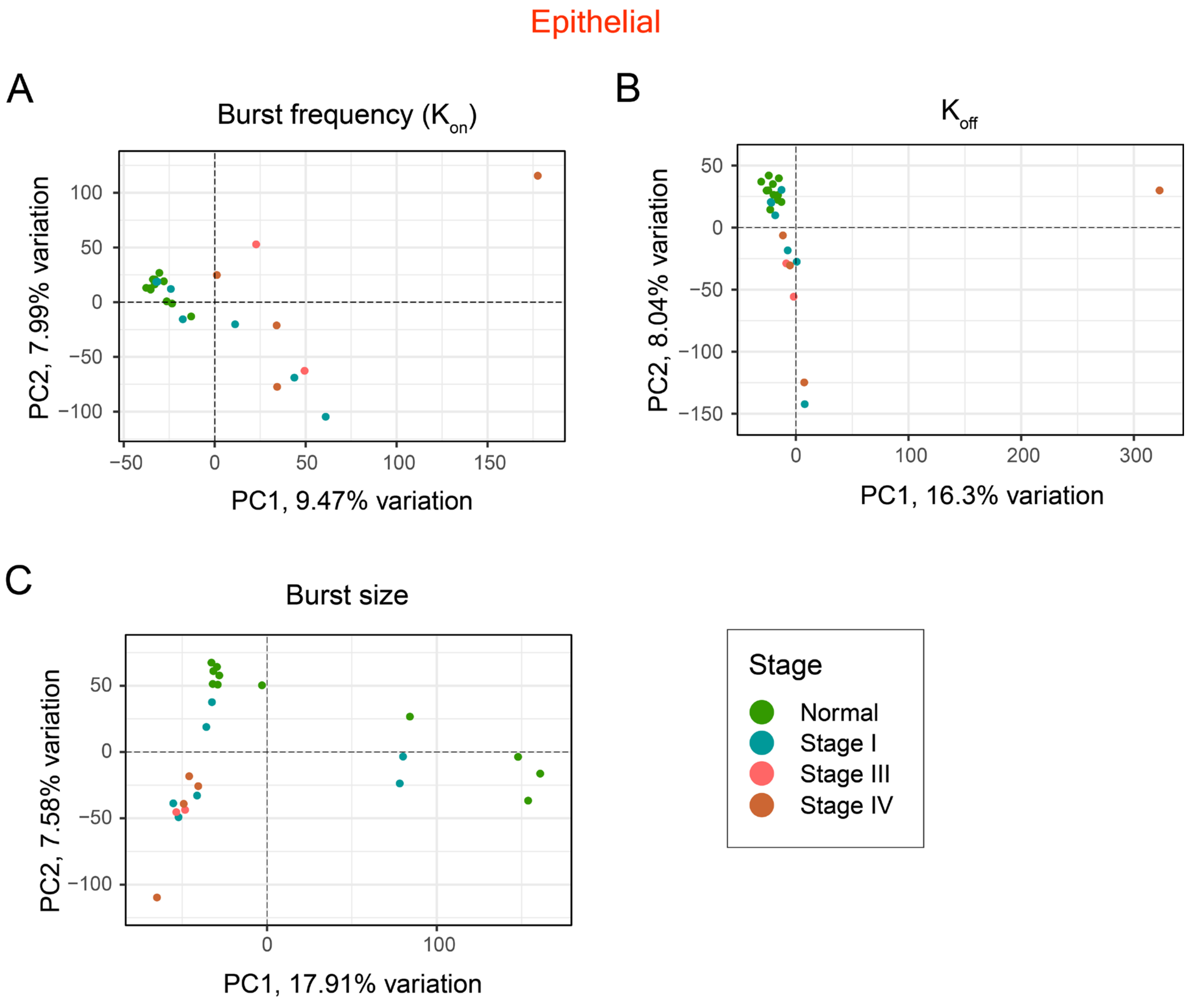


**Figure S29 -** Principal Component Analysis (PCA) of all samples based on burst frequency ($K_{on}$) **(A)**, $K_{off}$ **(B)**, and burst size **(C)** shows separation of normal and cancer samples in epithelial cells.

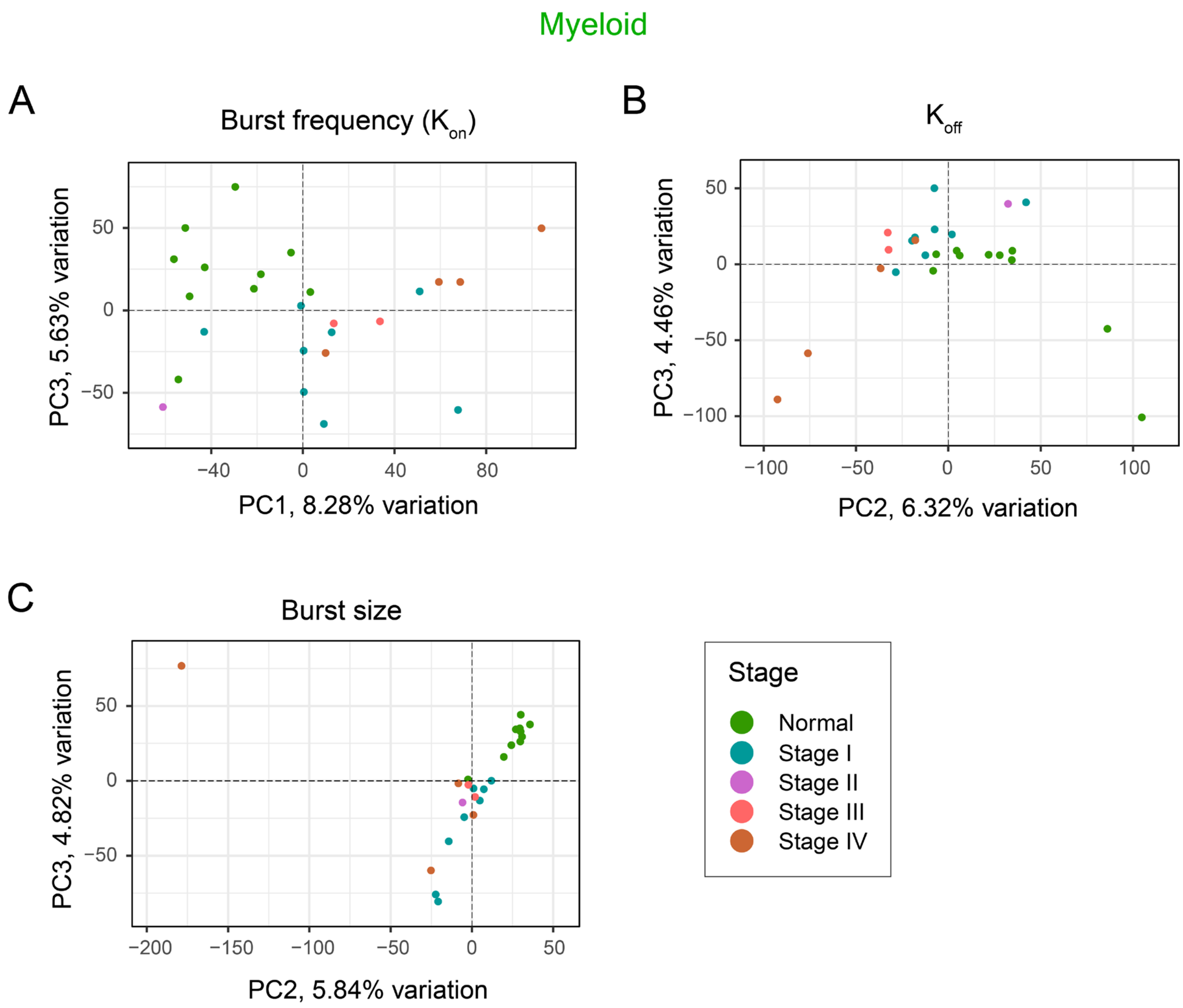


**Figure S30 -** Principal Component Analysis (PCA) of all samples based on burst frequency ($K_{on}$) **(A)**, $K_{off}$ **(B)**, and burst size **(C)** shows separation of normal and cancer samples in myeloid cells.

## T-NK

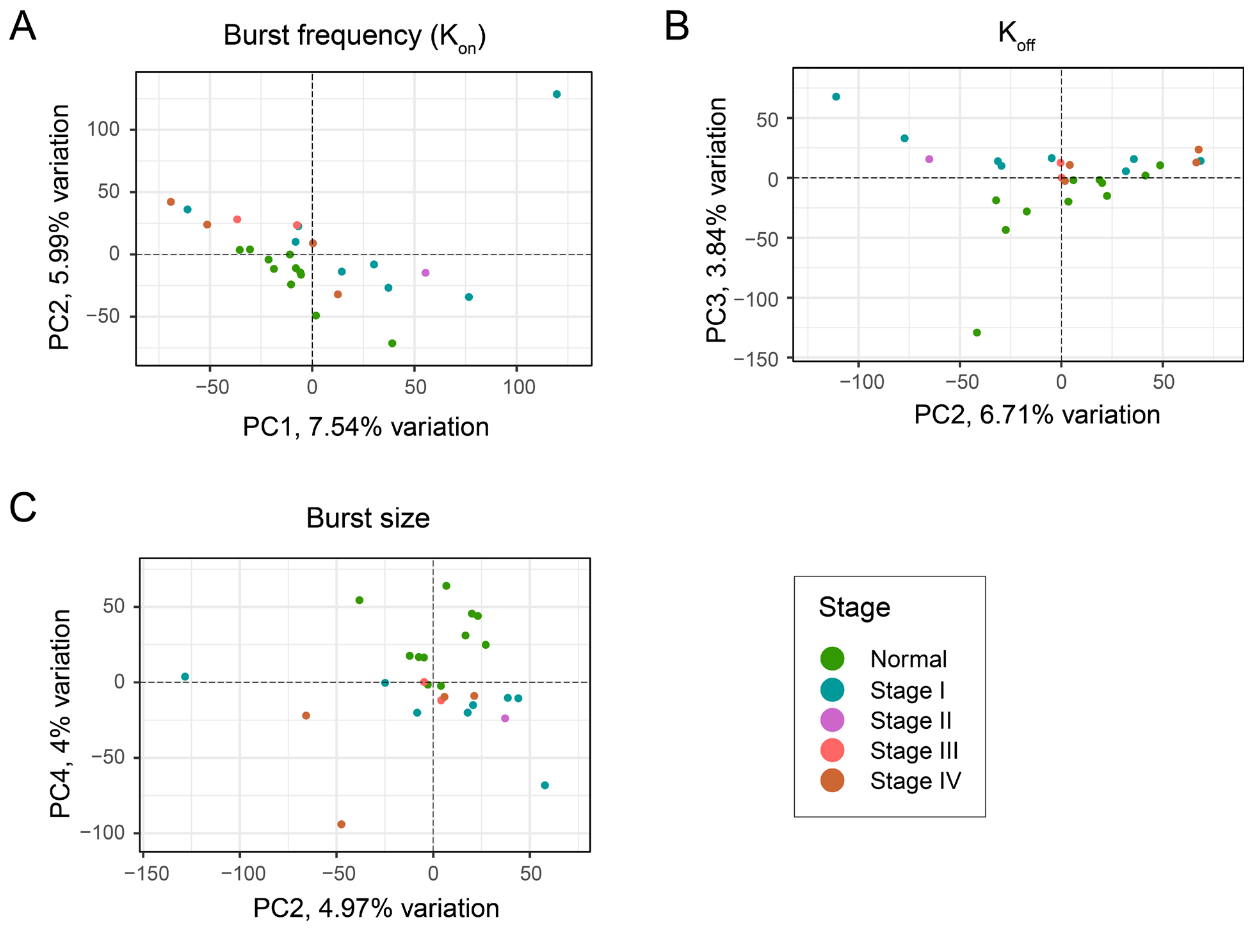


**Figure S31 -** Principal Component Analysis (PCA) of all samples based on burst frequency ($K_{on}$) **(A)**, $K_{off}$ **(B)**, and burst size **(C)** shows separation of normal and cancer samples in T-NK cells.

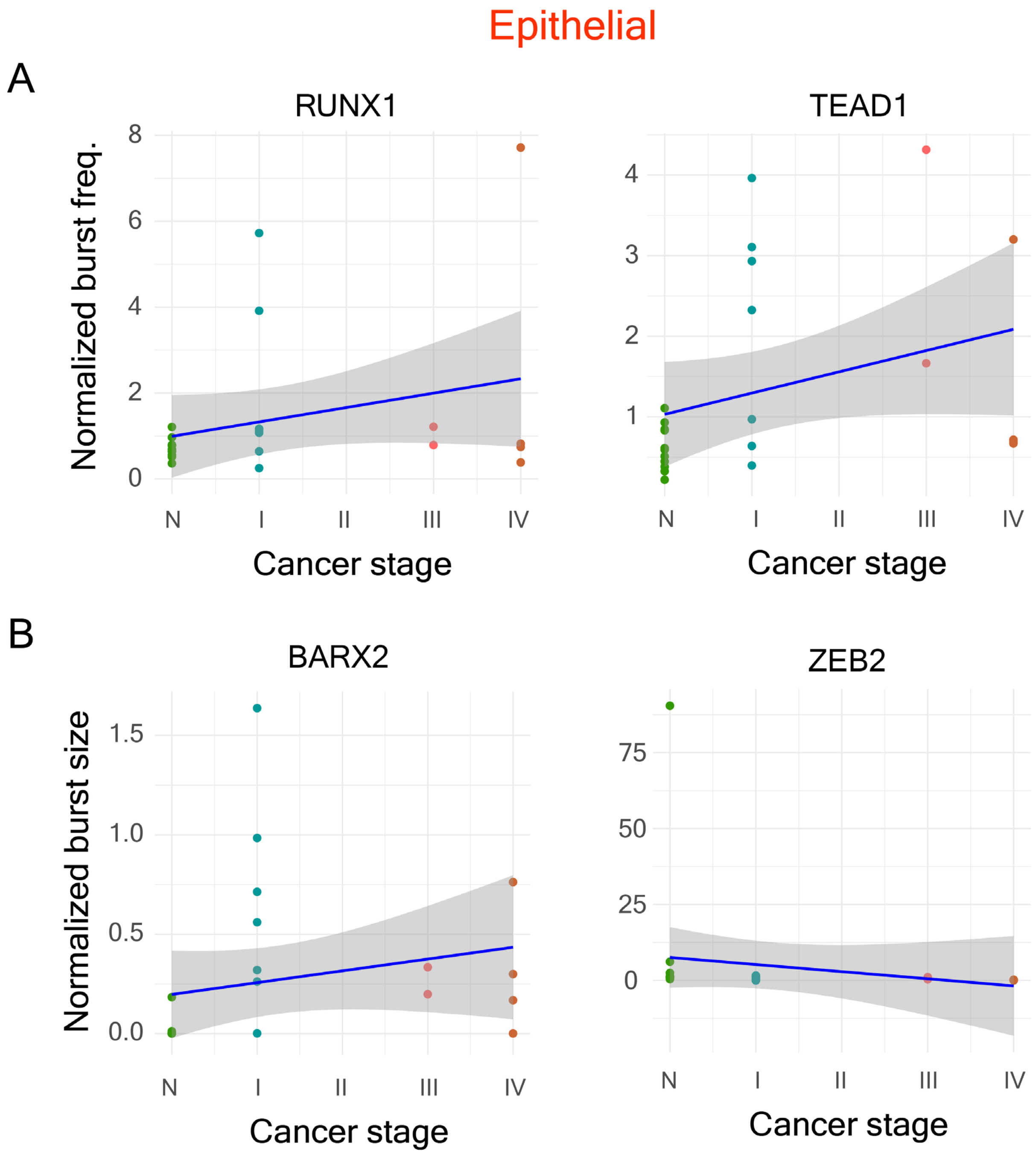


**Figure S32 –** Changes in burst frequency ($K_{on}$) **(A)**, and burst size **(B)** of some of the representative genes in epithelial cells with cancer progression.

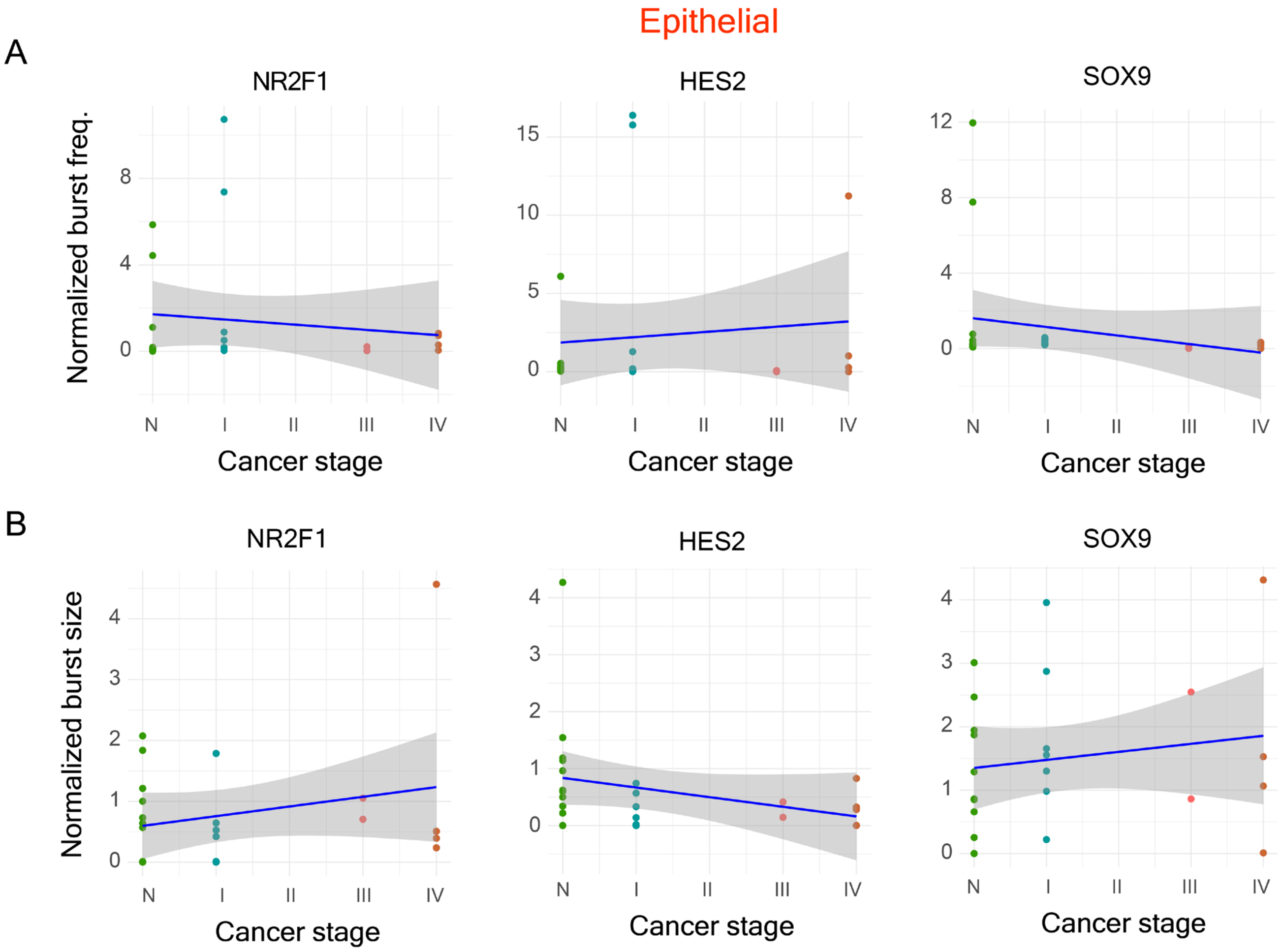


**Figure S33 –** Burst frequency ($K_{on}$) **(A)**, and burst size **(B)** of the hub TFs (identified from correlation trend analysis) in epithelial cells in normal and cancer samples.

#### Myeloid TFs

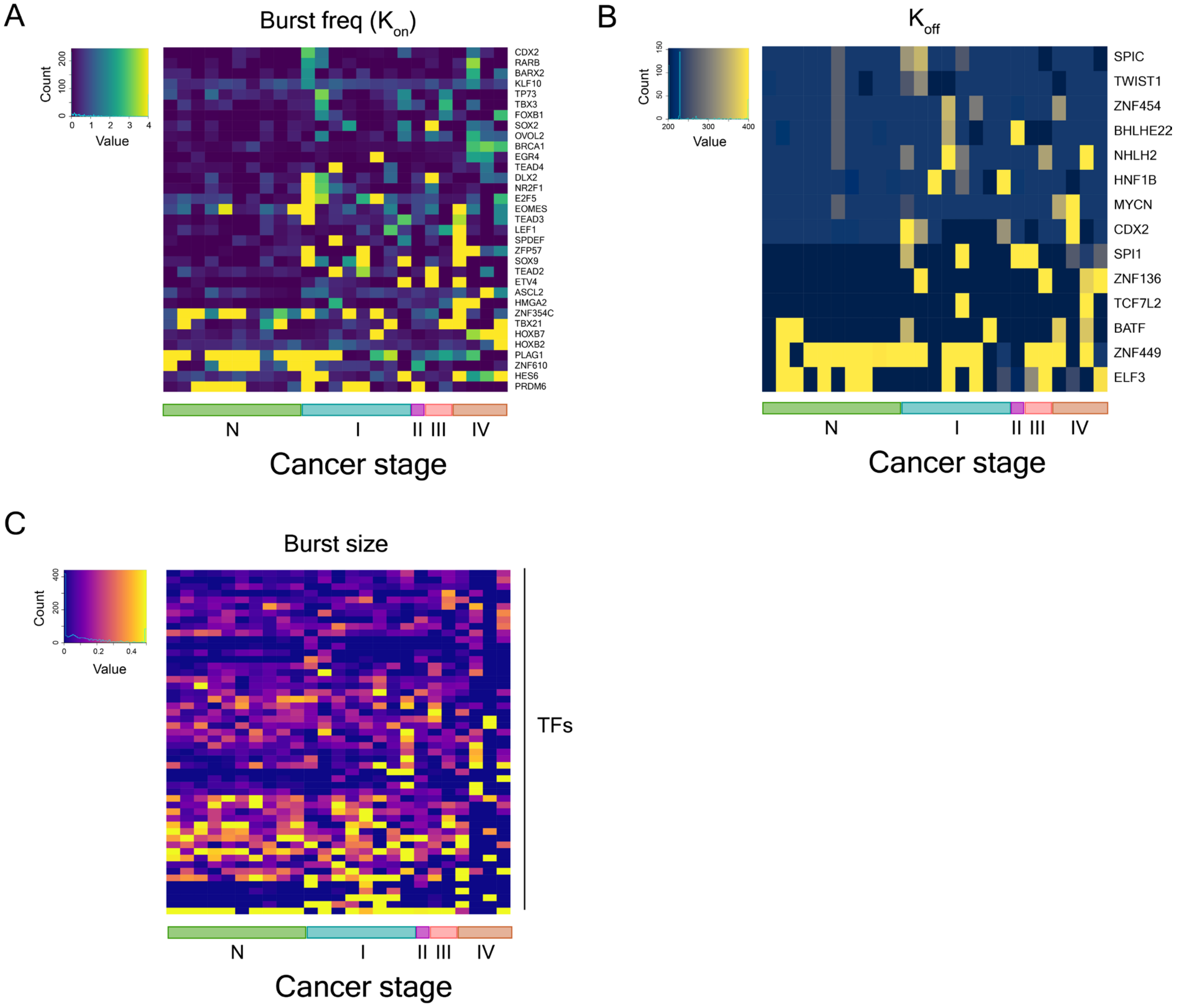


**Figure S34 -** Heatmaps showing the changes in burst frequency **(A)**, $K_{off}$ **(B)**, and burst size **(C)** in myeloid cells of the TFs identified to have different burst kinetics in cancer samples compared to normal samples.

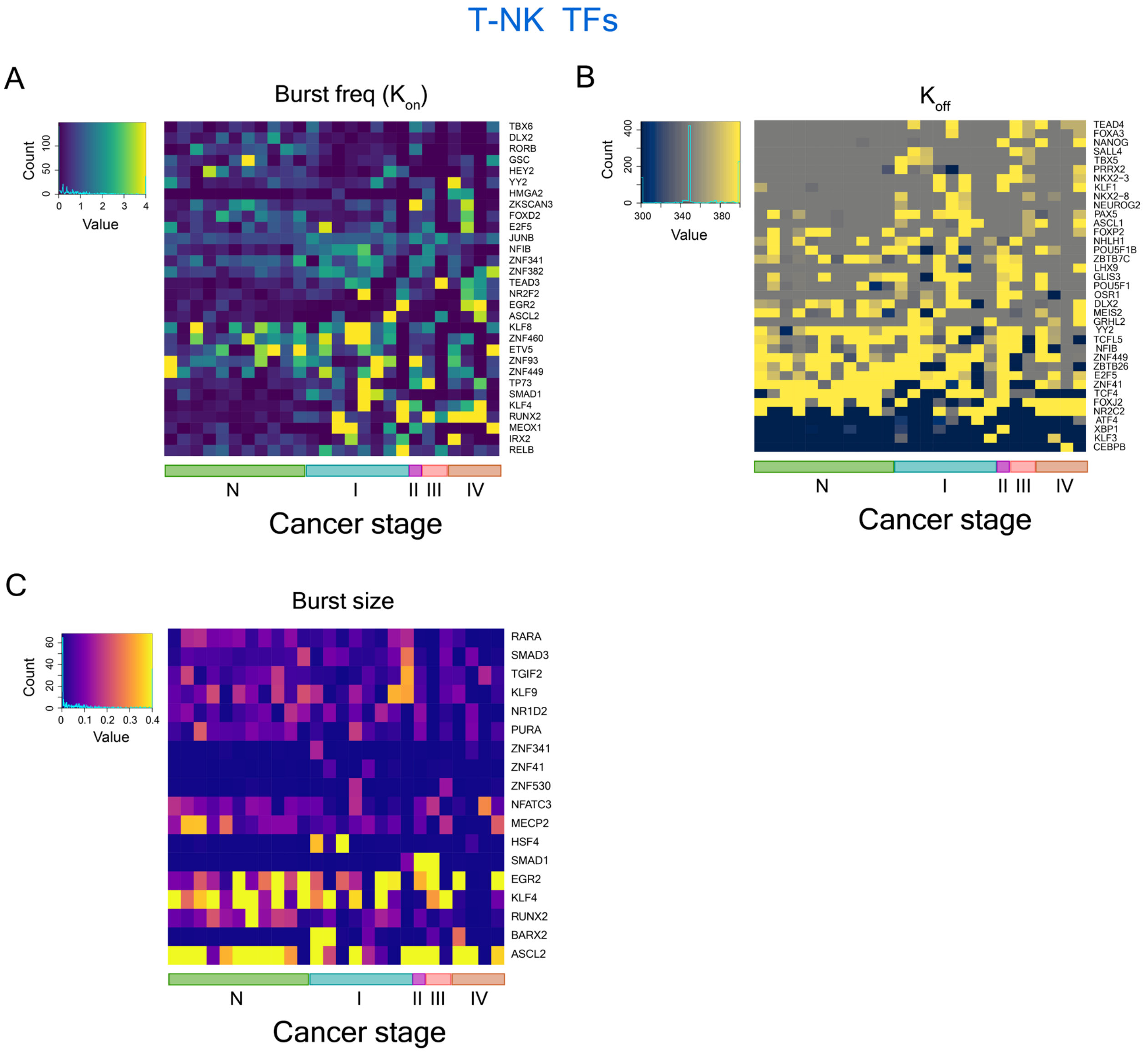


**Figure S35 -** Heatmaps showing the changes in burst frequency **(A)**, $K_{off}$ **(B)**, and burst size **(C)** in T-NK cells of the TF genes identified to have different burst kinetics in cancer samples compared to normal samples.

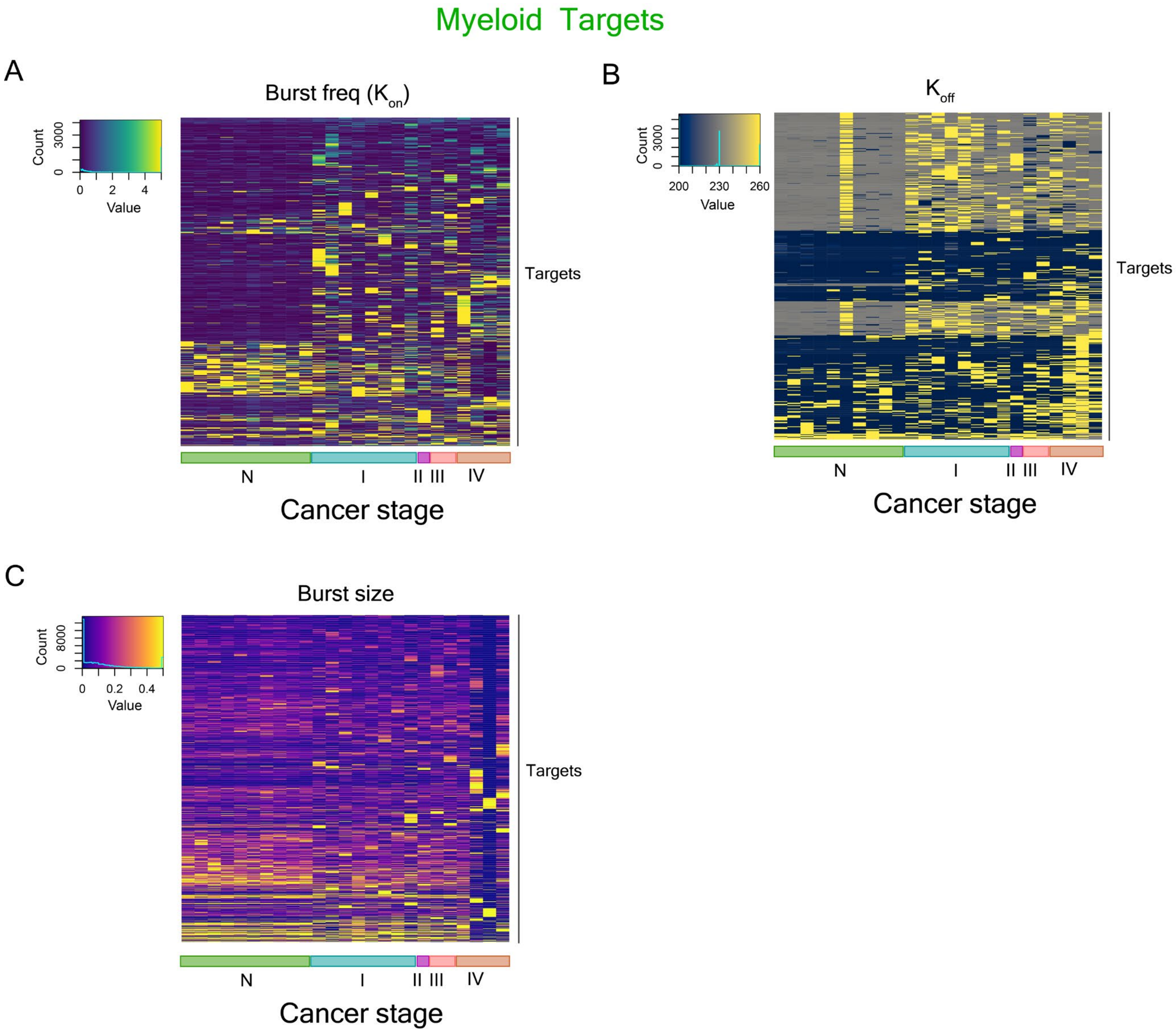


**Figure S36 -** Heatmaps showing the changes in burst frequency **(A)**, $K_{off}$ **(B)**, and burst size **(C)** in myeloid cells of the target genes identified to have different burst kinetics in cancer samples compared to normal samples. Only the target genes for which TF-target map was known were considered.

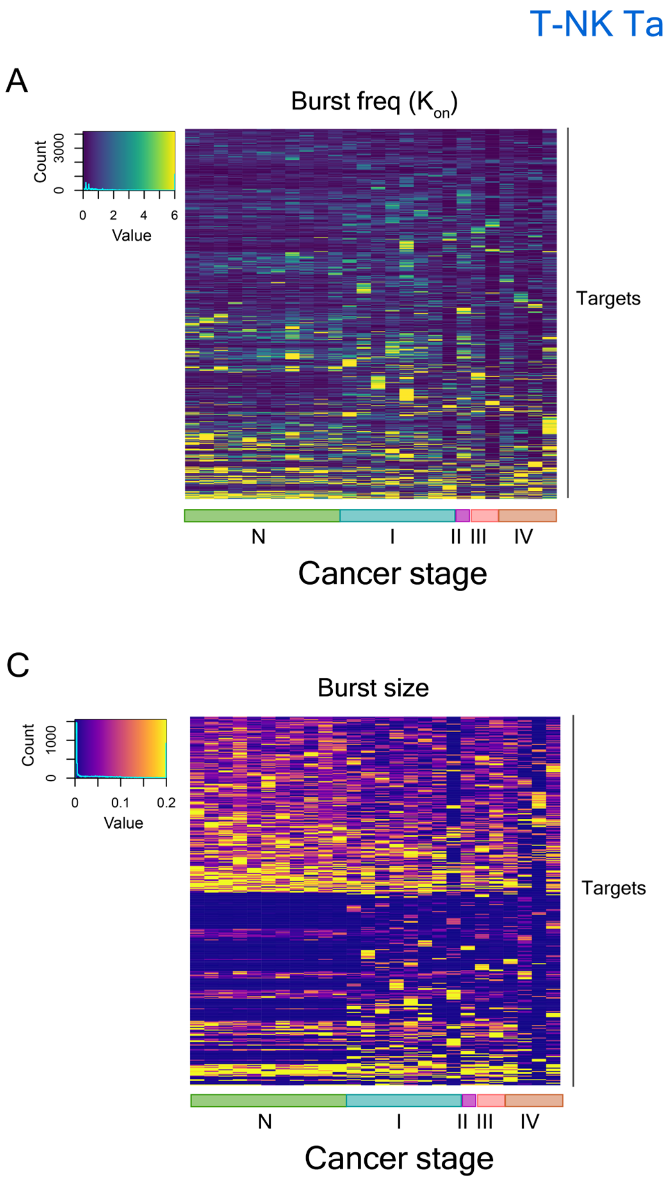


**Figure S37 -** Heatmaps showing the changes in burst frequency **(A)**, $K_{off}$ **(B)**, and burst size **(C)** in T-NK cells of the target genes identified to have different burst kinetics in cancer samples compared to normal samples. Only the target genes for which TF-target map was known were considered.

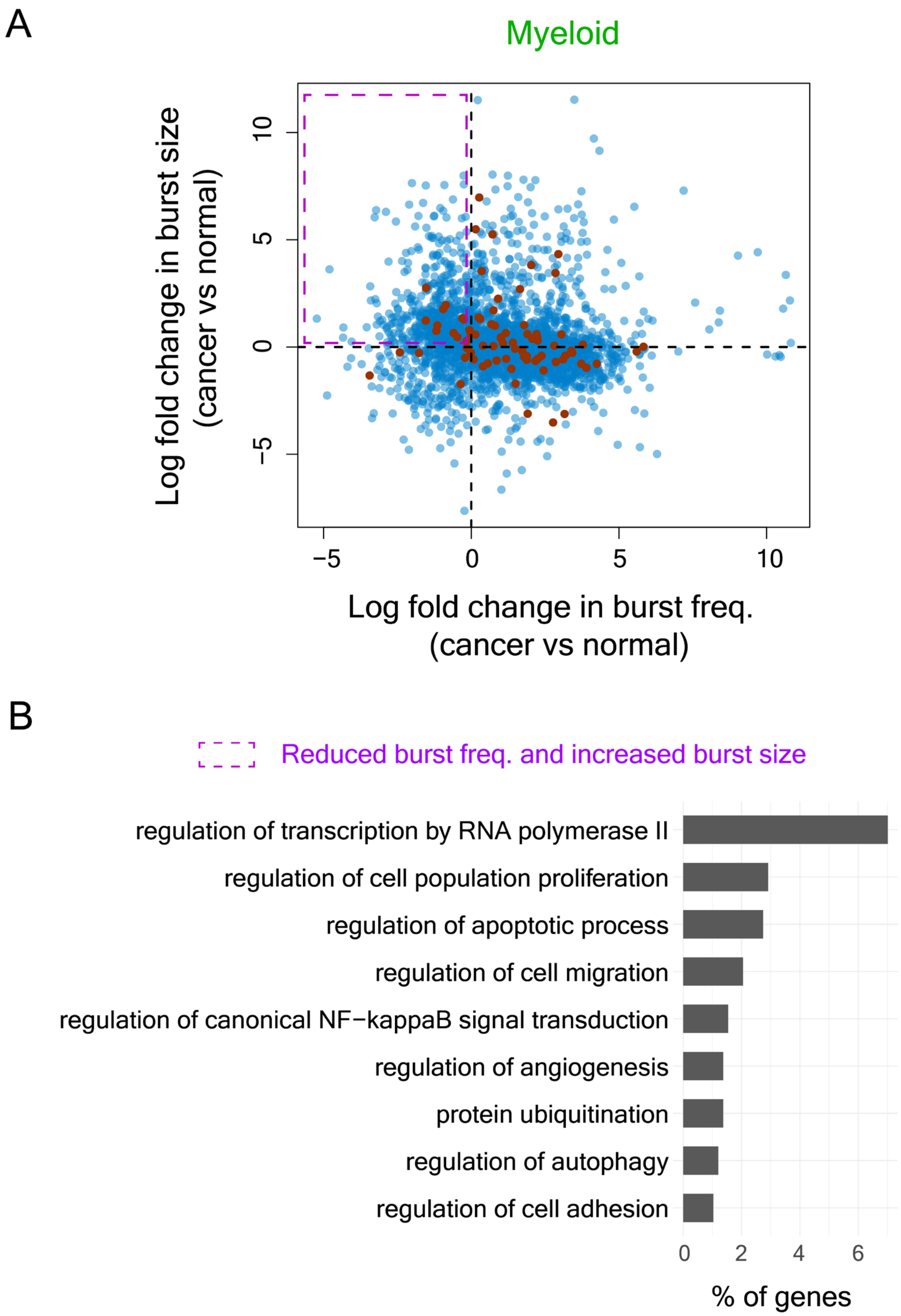


**Figure S38 – (A)** Log fold change in burst frequency vs log fold change in burst size in myeloid cells for all genes included in the analysis. **(B)** Functional association with biological processes in Gene Ontology (GO) of genes that show decreased burst frequency and increased burst size in myeloid cells of cancer samples compared to normal myeloid cells.

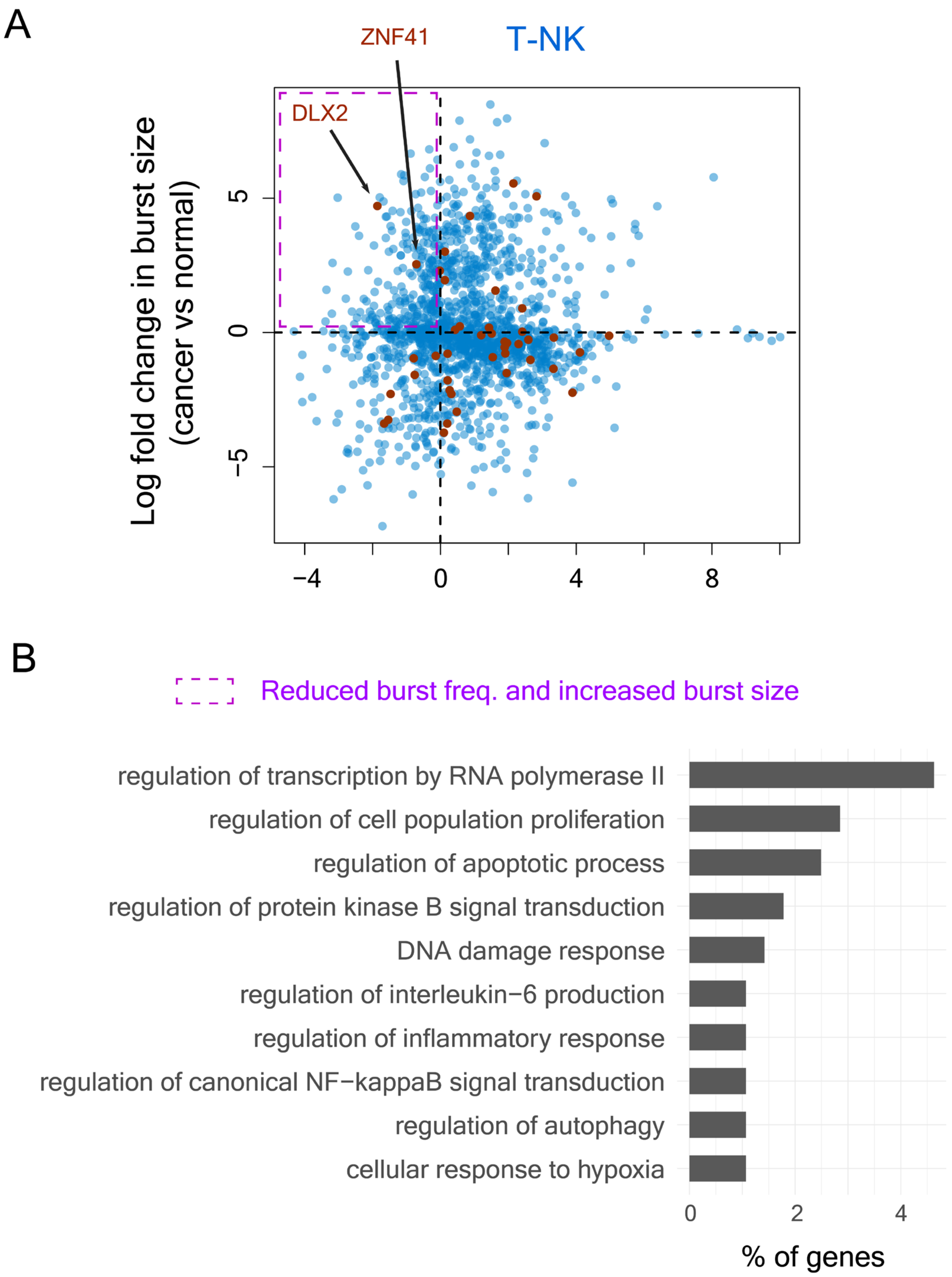


**Figure S39 - (A)** Log fold change in burst frequency vs log fold change in burst size in T-NK cells for all genes included in the analysis. **(B)** Functional association with biological processes in Gene Ontology (GO) of genes that show decreased burst frequency and increased burst size in T-NK cells of cancer samples compared to normal T-NK cells.

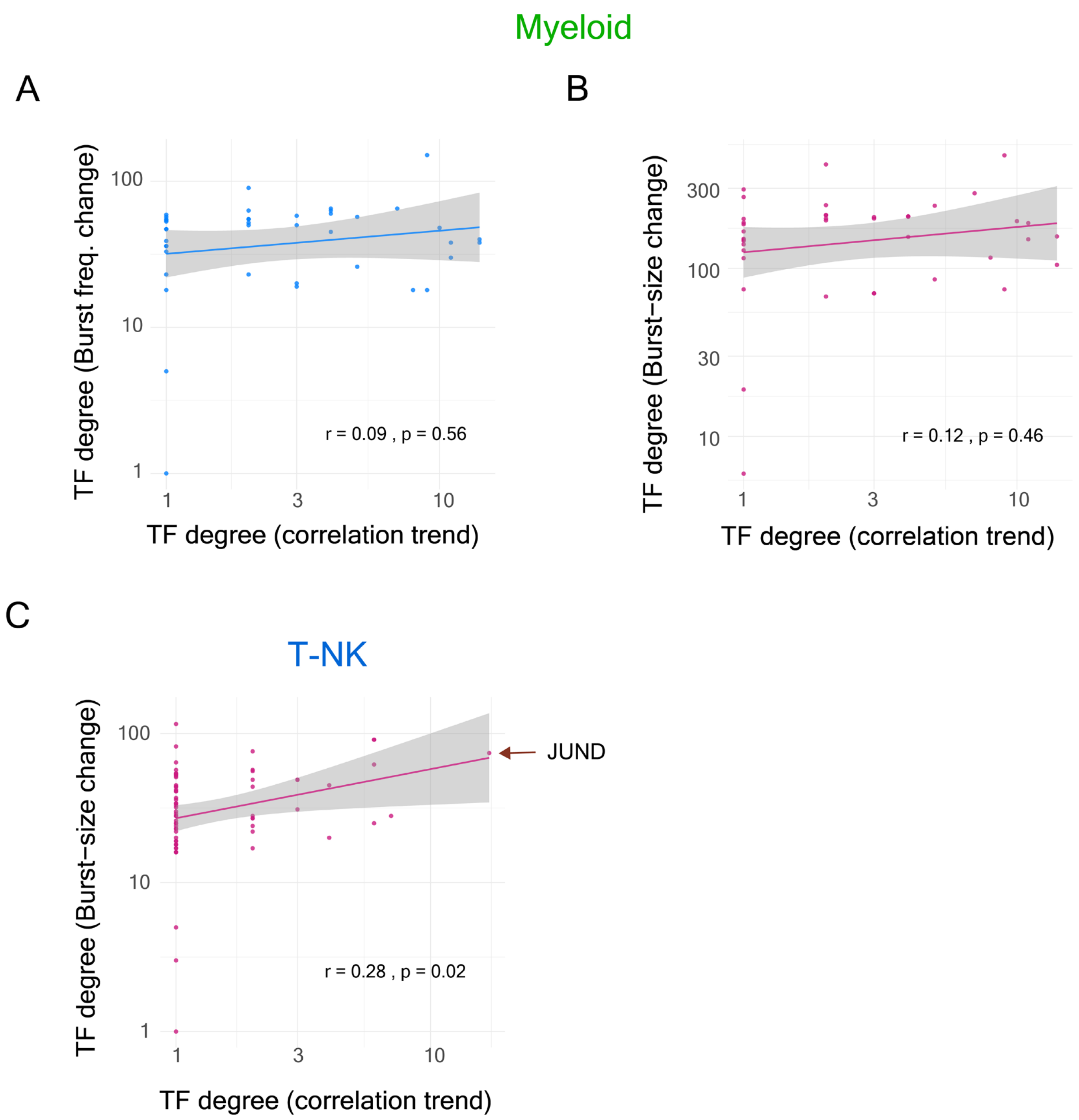


**Figure S40 – (A)** Correlation between TF degree identified from correlation trend analysis and TF degree identified from burst frequency change analysis in myeloid cells. **(B)** Correlation between TF degree identified from correlation trend analysis and TF degree identified from burst size change analysis in myeloid cells. **(C)** Correlation between TF degree identified from correlation trend analysis and TF degree identified from burst size change analysis in T-NK cells.

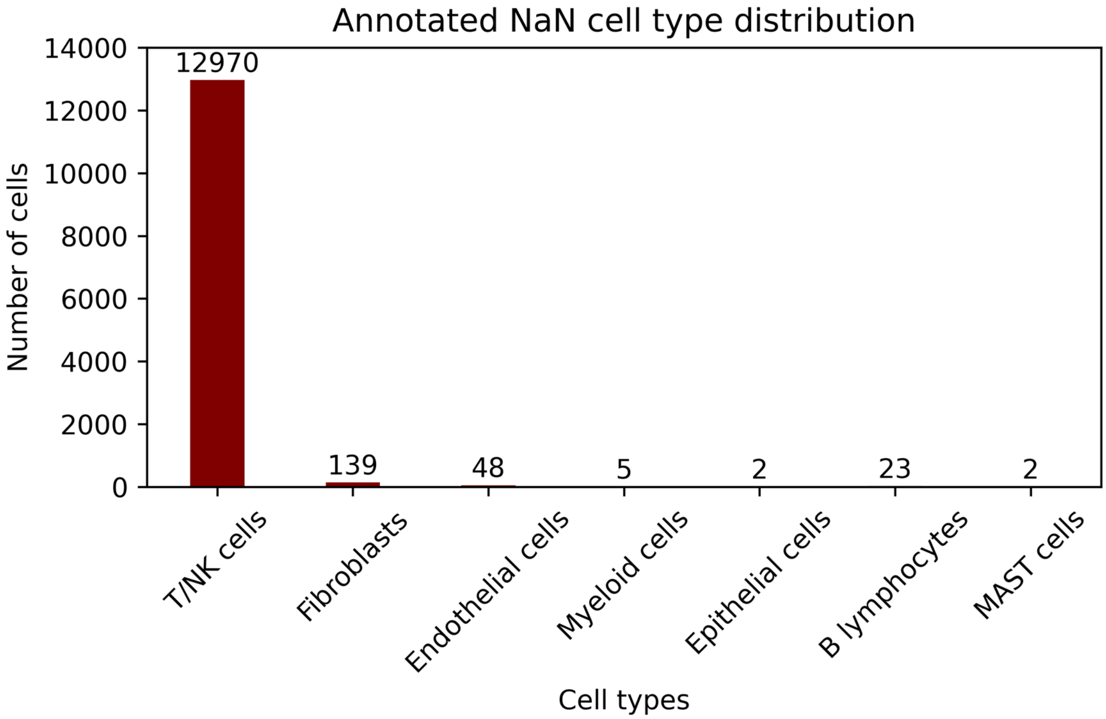


**Figure S41** – Annotation of non-annotated cells into specific cell types by Seurat v4 (Hao et al., 2021) and SciBet (Li et al., 2020). Cells that were assigned same cell type from both these methods were included in the analysis of which 98.3% were annotated as T-NK cells.

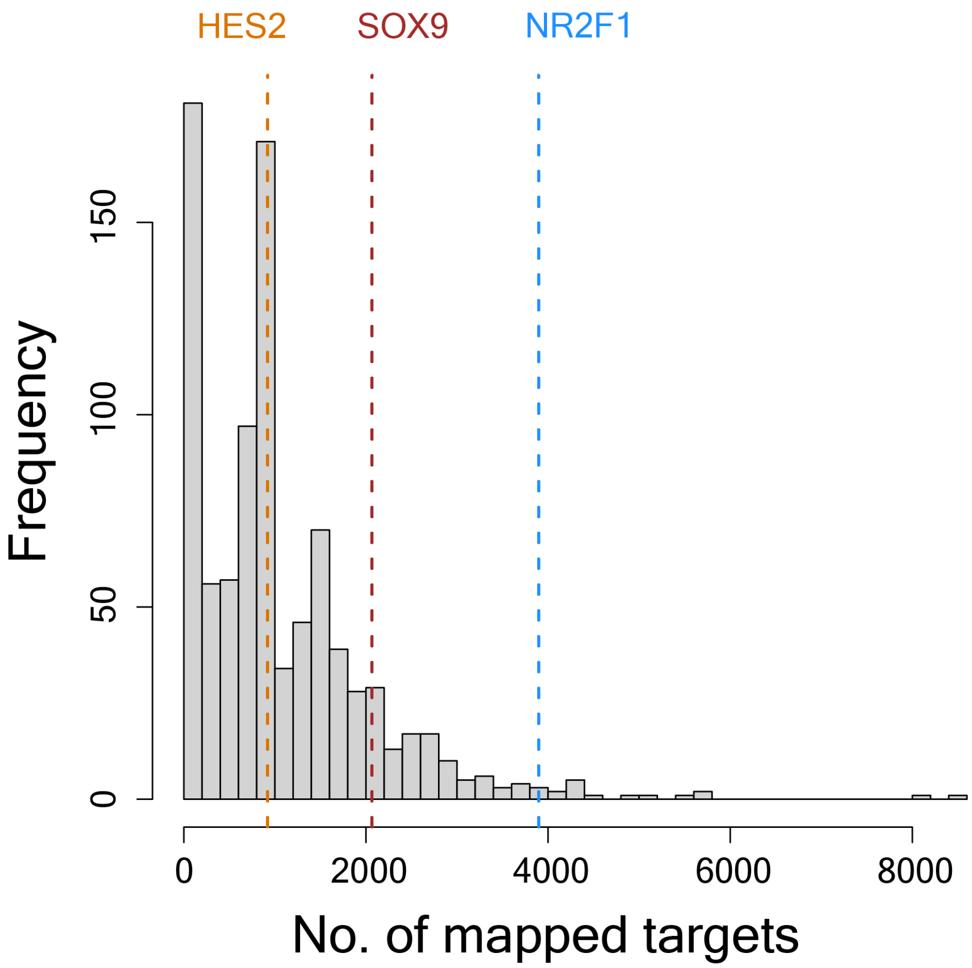


**Figure S42 –** Histogram showing the frequency of the TFs with given number of target genes (shown in the x-axis) in our TF-target map. The dotted lines show the number of targets of the TFs NR2F1, HES2 and SOX9 in our TF-target map.

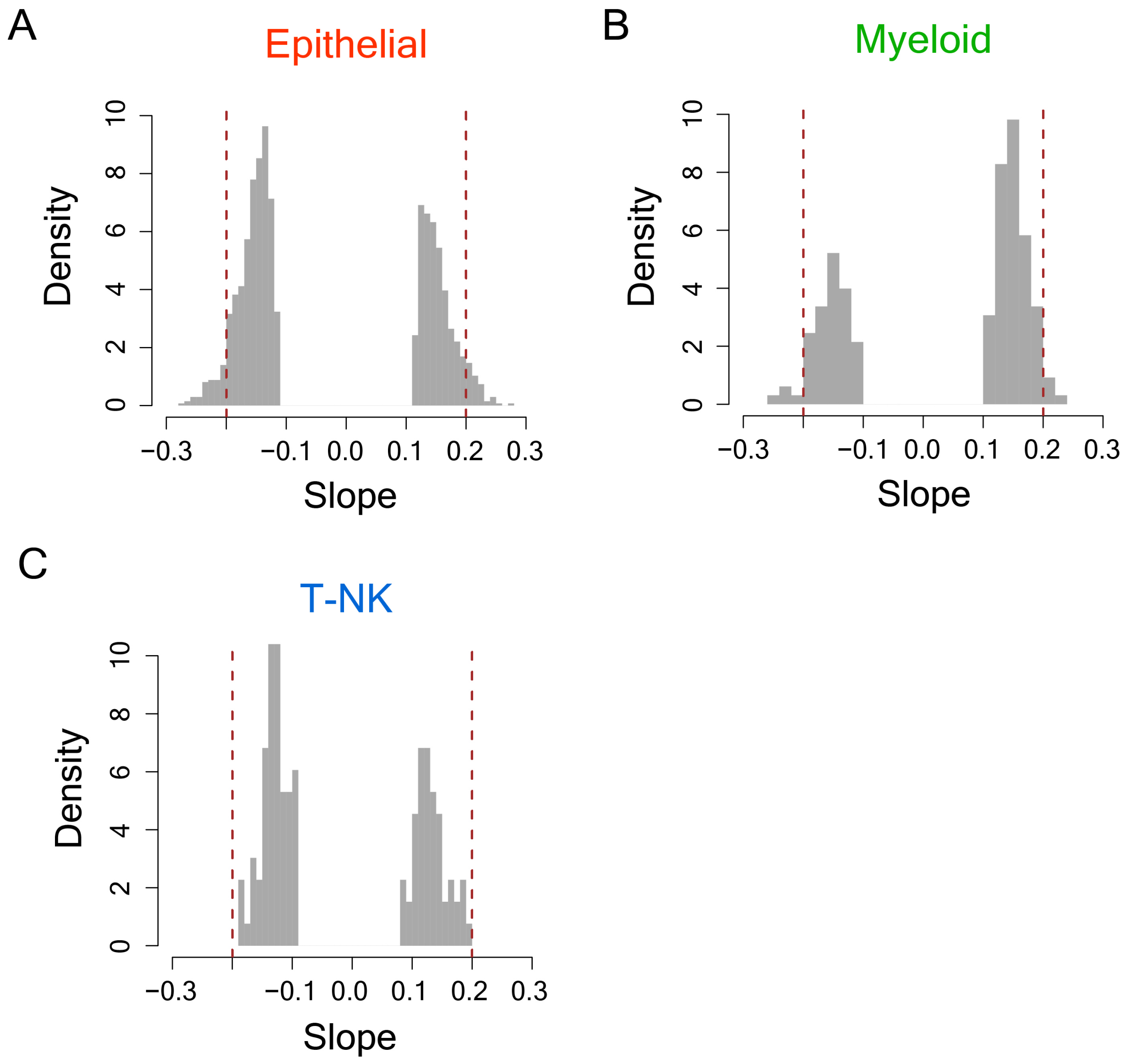


**Figure S43 -** Distribution of slope values of the regression lines for the filtered TF-target connections from correlation trend analysis in epithelial cells **(A)**, myeloid cells **(B)** and T-NK cells **(C)**.
