## Supplementary figures and images for "Gene regulatory network transitions reveal the central transcription factors in lung adenocarcinoma progression"

### File S1

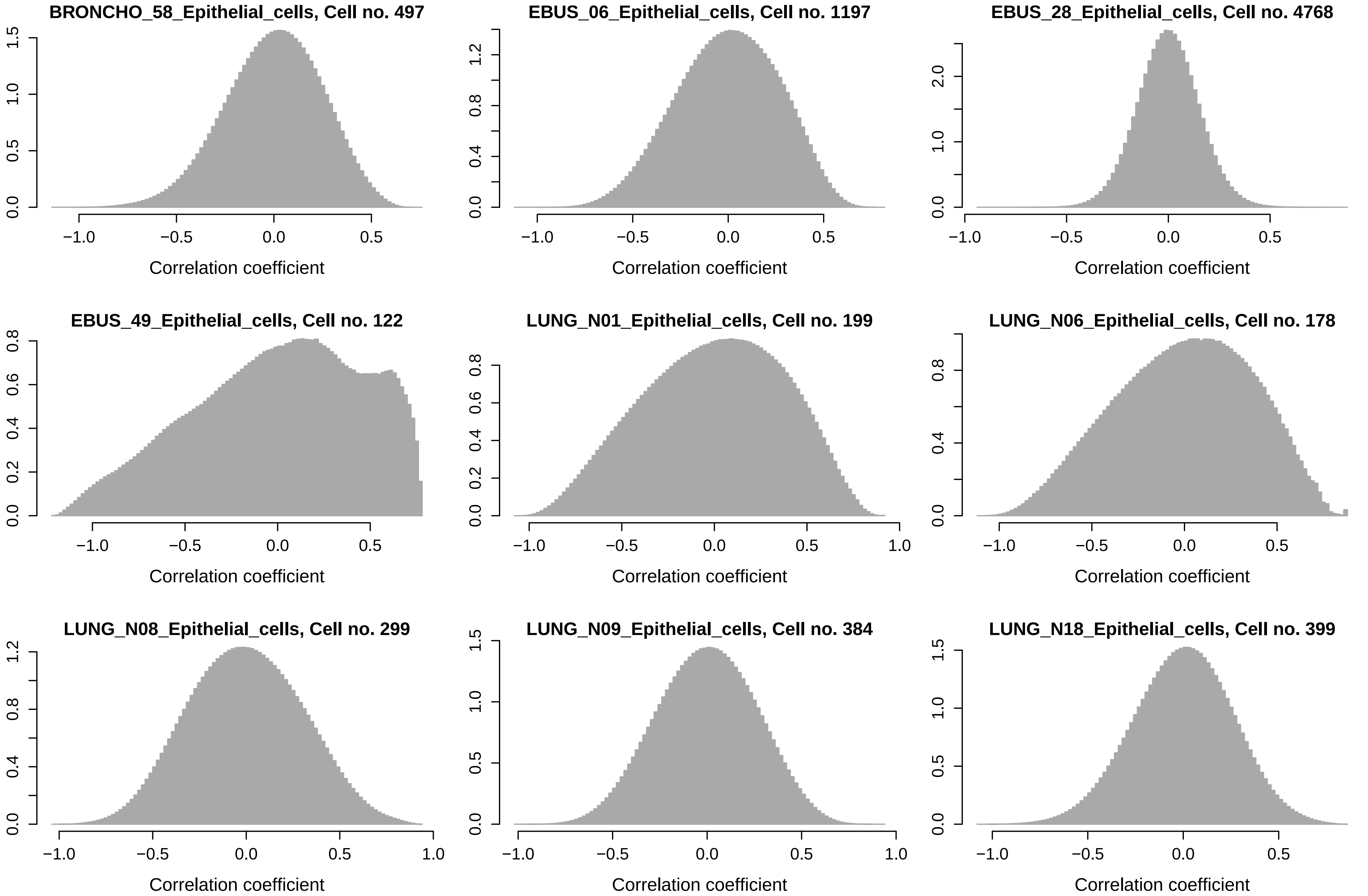

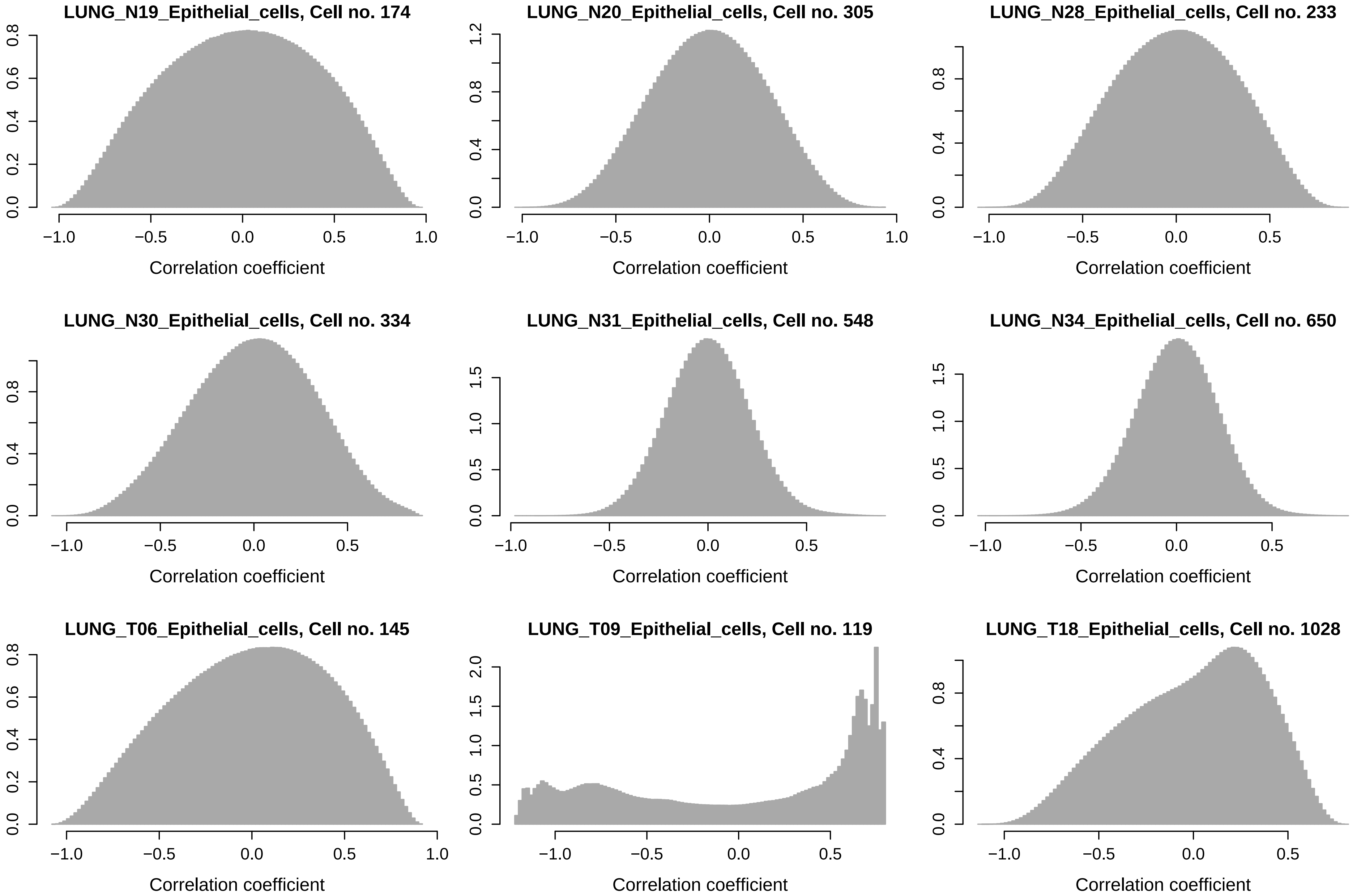

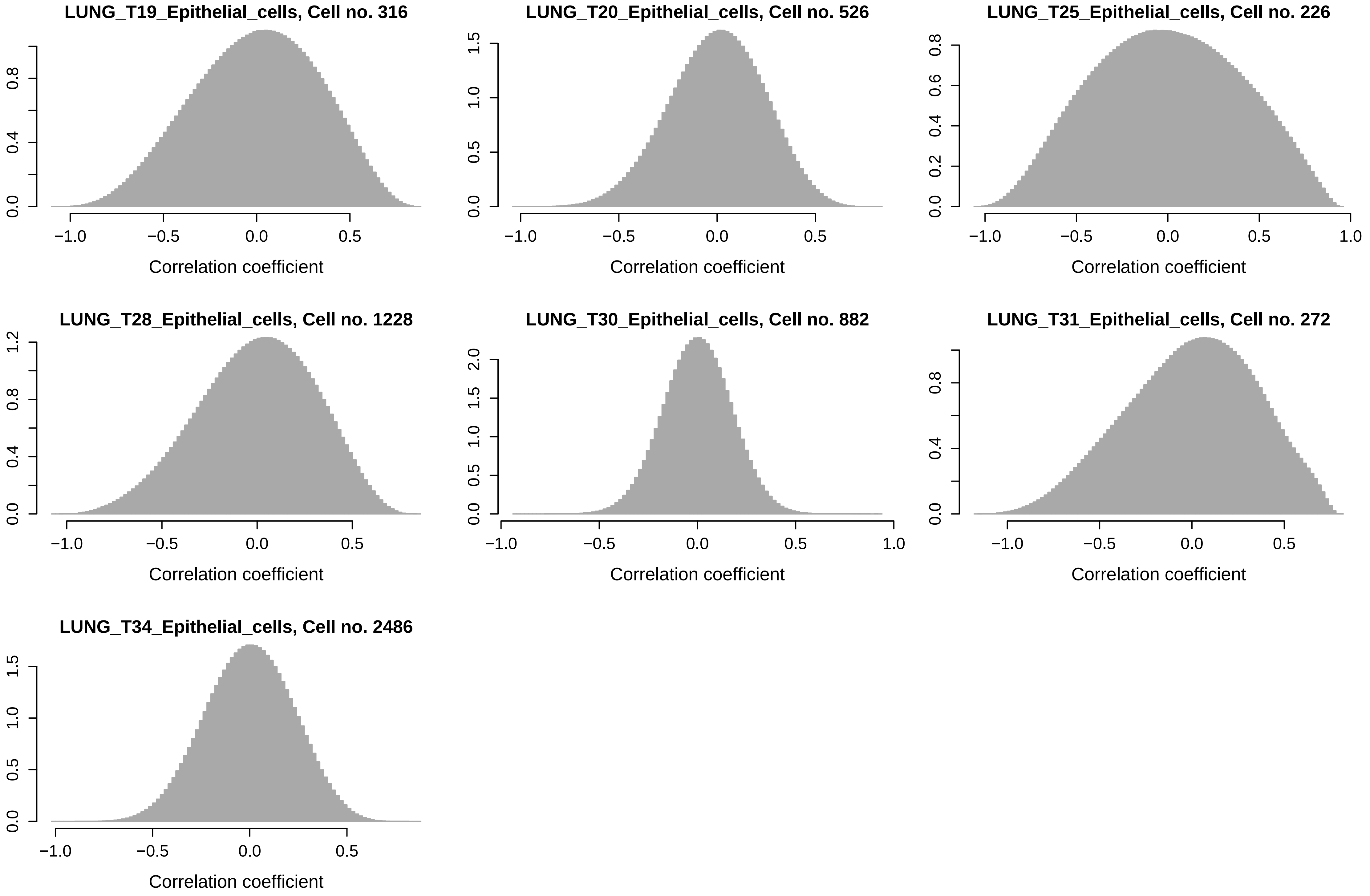

### File S2

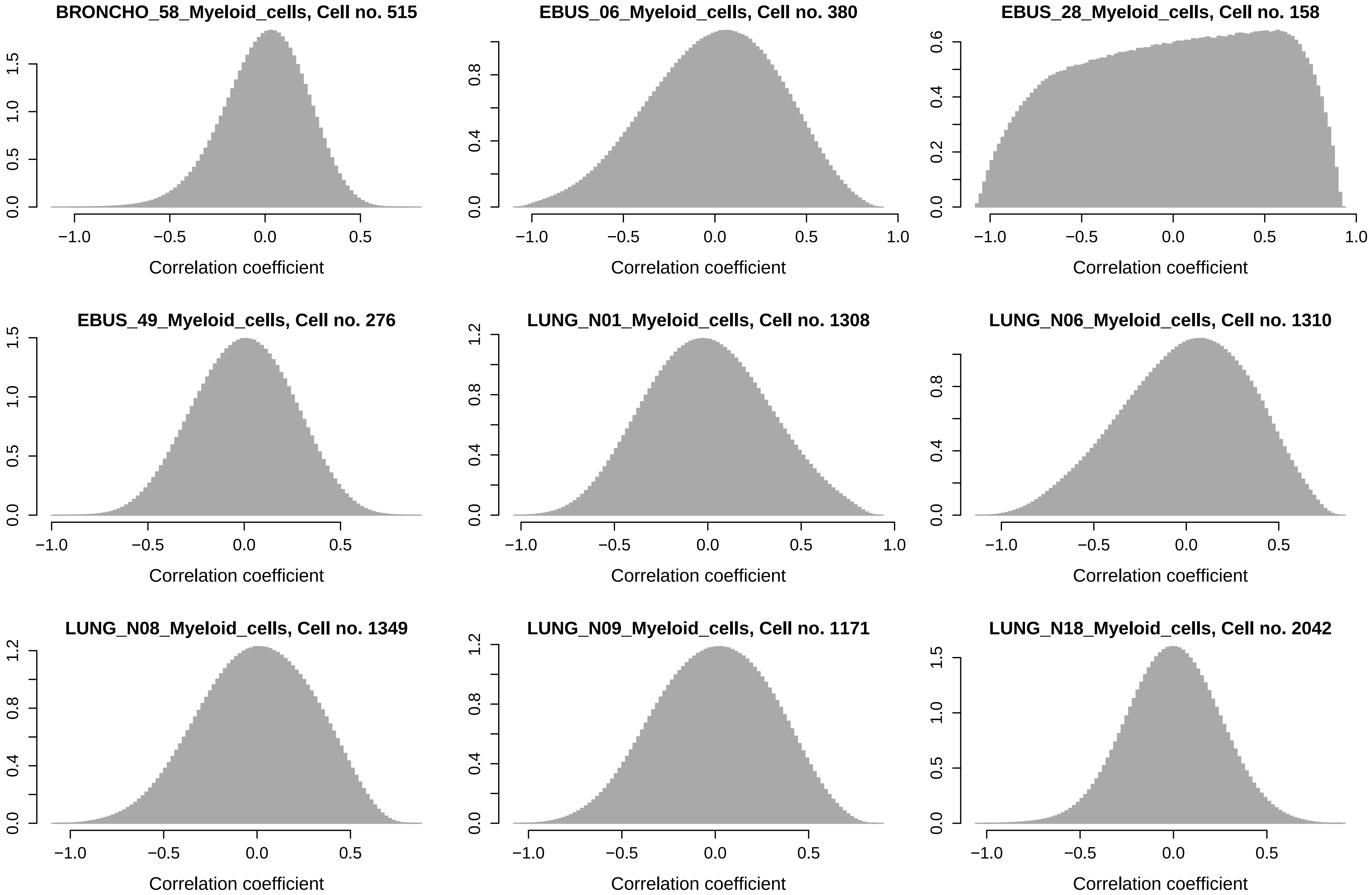

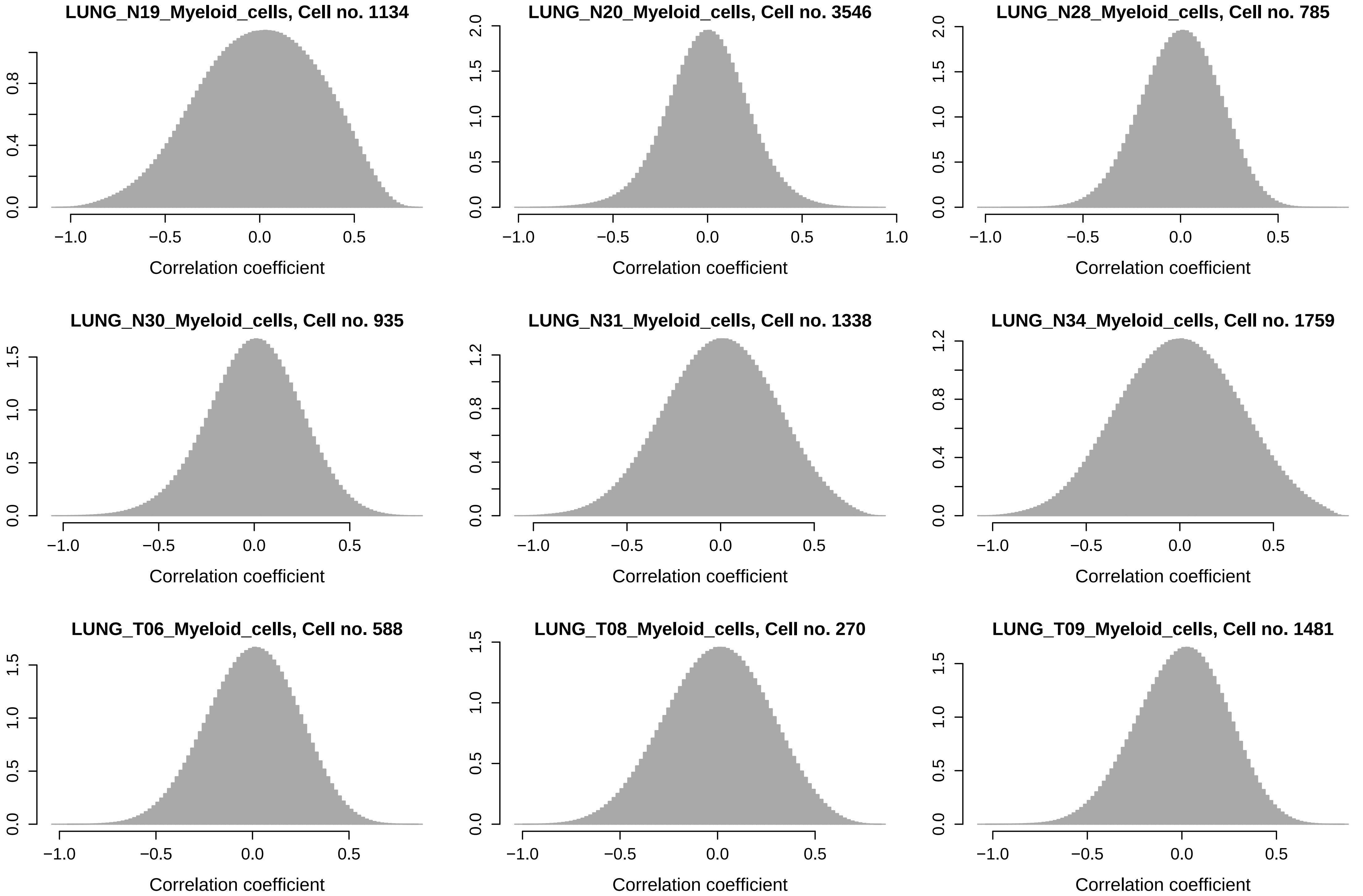

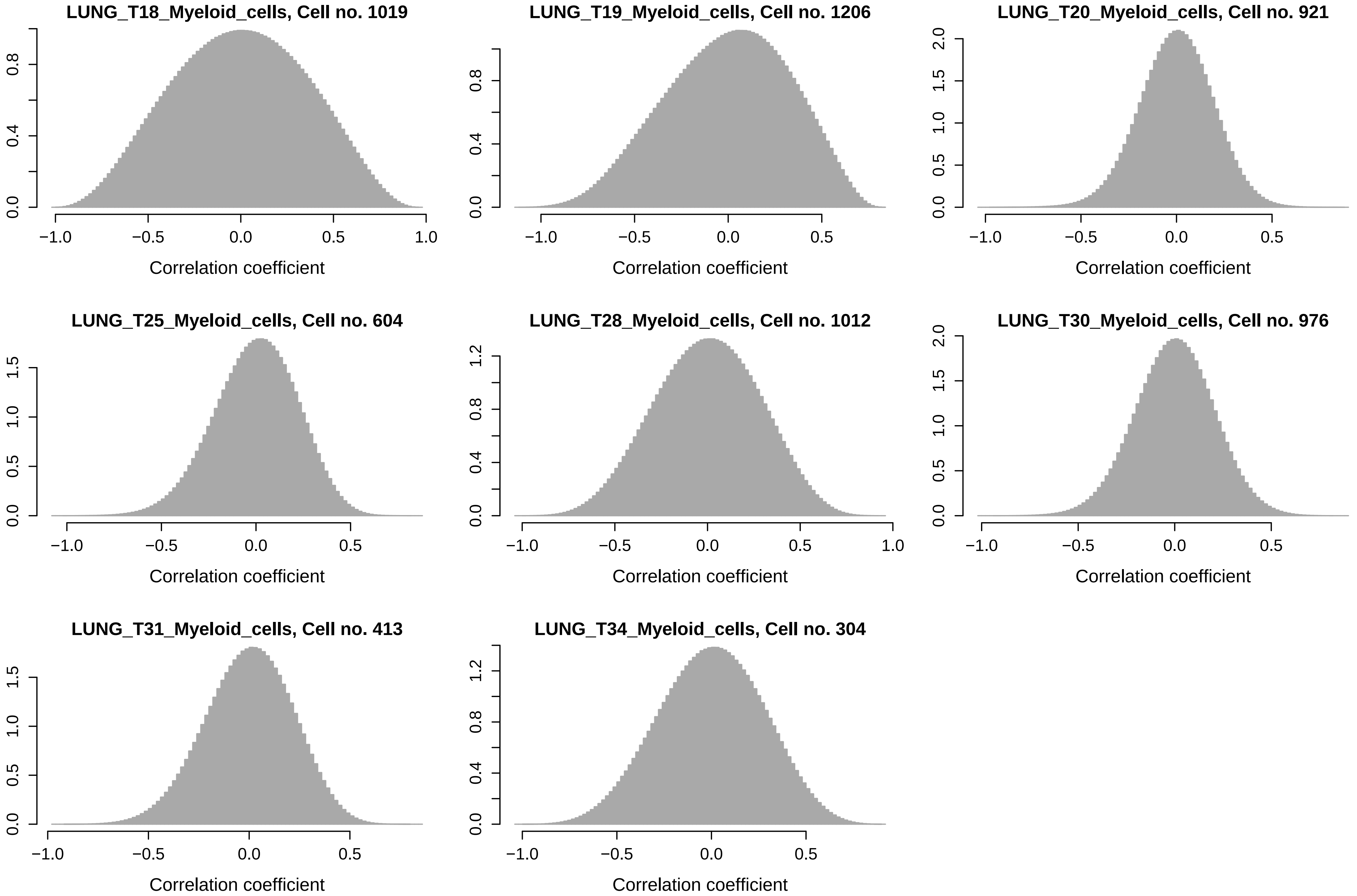

### File S3

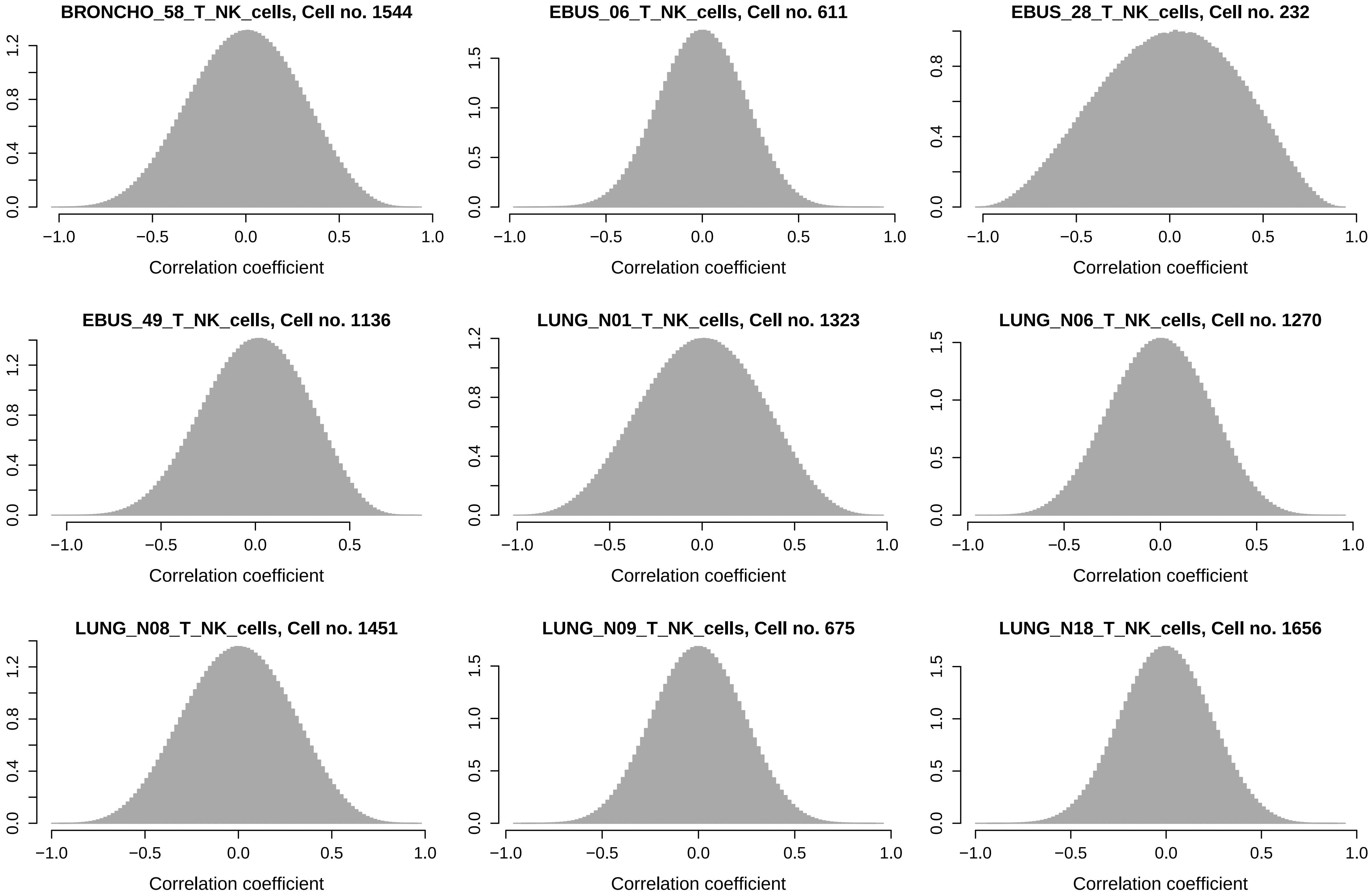

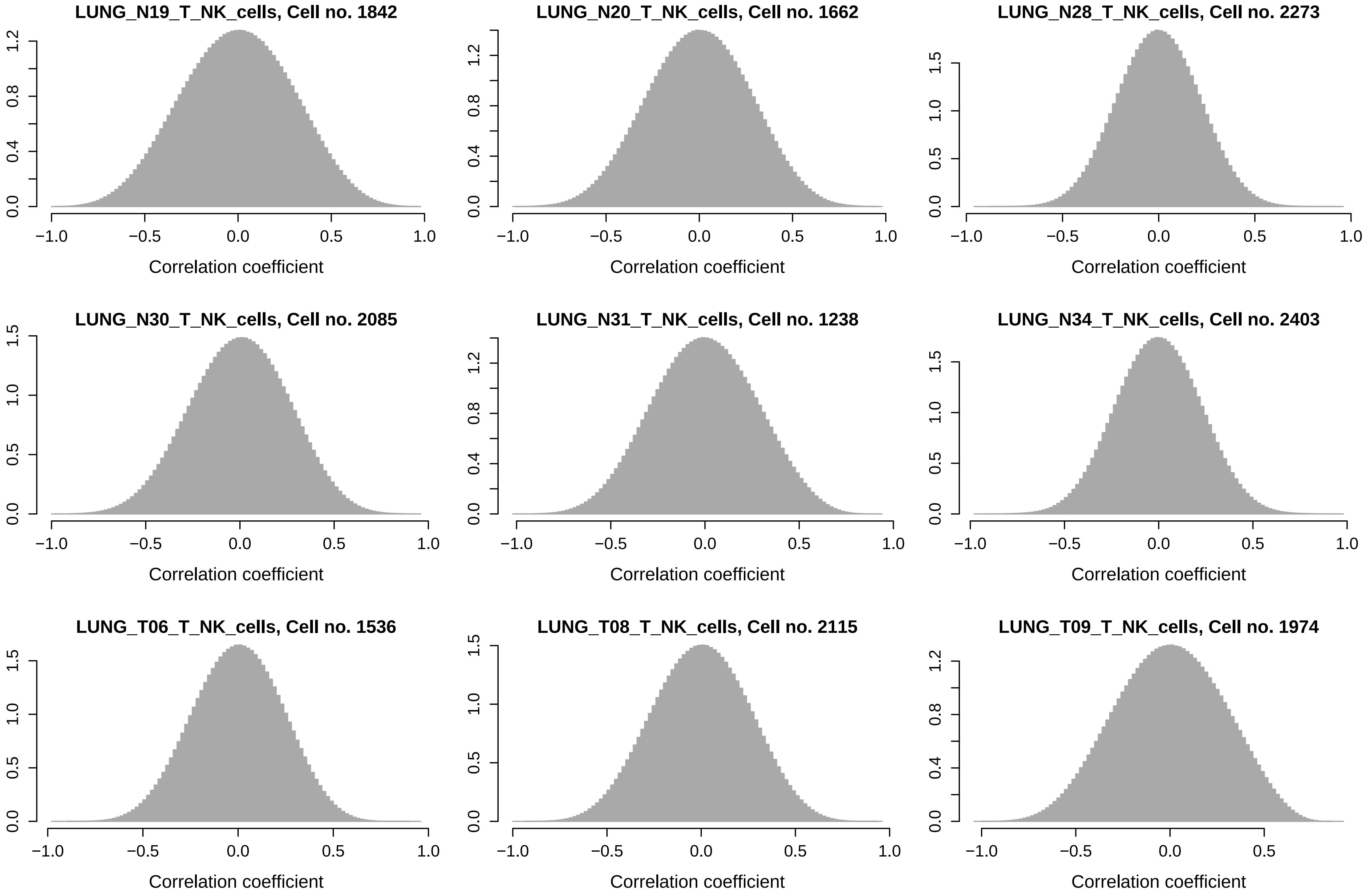

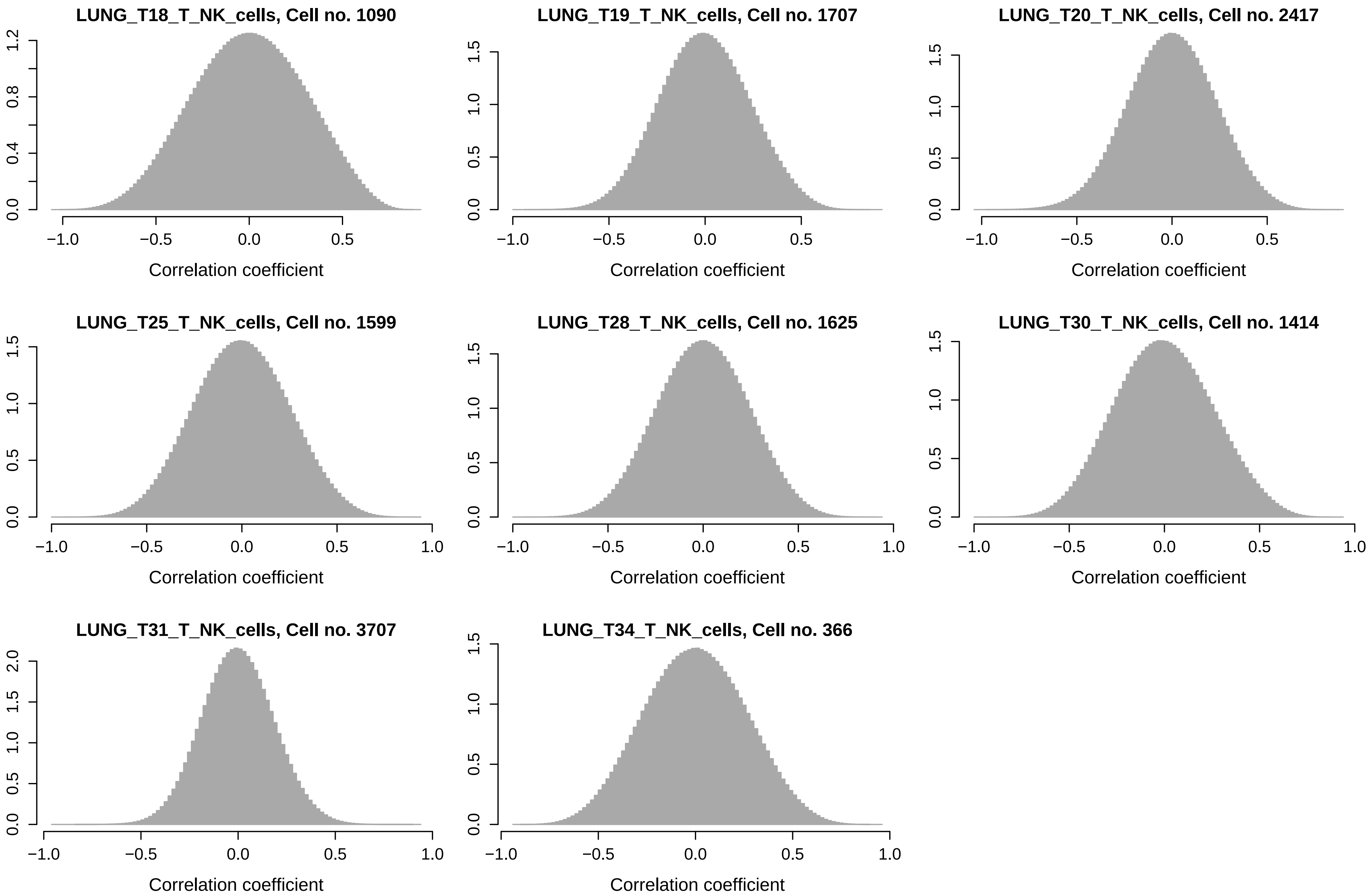
